# The whale shark genome reveals how genomic and physiological properties scale with body size

**DOI:** 10.1101/443036

**Authors:** Seung Gu Park, Victor Luria, Jessica A. Weber, Sungwon Jeon, Hak-Min Kim, Yeonsu Jeon, Youngjune Bhak, Jehun Jun, Sang Wha Kim, Won Hee Hong, Semin Lee, Yun Sung Cho, Amir Karger, John W. Cain, Andrea Manica, Soonok Kim, Jae-Hoon Kim, Jeremy S. Edwards, Jong Bhak, George M. Church

**Affiliations:** Korean Genomics Industrialization and Commercialization Center (KOGIC), Ulsan National Institute of Science and Technology (UNIST), Ulsan 44919, Republic of Korea; Department of Biomedical Engineering, School of Life Sciences, Ulsan National Institute of Science and Technology (UNIST), Ulsan 44919, Republic of Korea; Department of Systems Biology, Harvard Medical School, Boston, MA 02115, USA; Department of Genetics, Harvard Medical School, Boston, MA 02115, USA; Department of Biology, University of New Mexico, Albuquerque, NM 87131, USA; Clinomics Inc., Ulsan 44919, Republic of Korea; Laboratory of Aquatic Biomedicine, College of Veterinary Medicine and Research Institute for Veterinary Science, Seoul National University, Seoul 08826, Republic of Korea; Hanwha Marine Biology Research Center, Jeju 63642, Republic of Korea; IT - Research Computing, Harvard Medical School, Boston, MA 02115, USA; Department of Mathematics, Harvard University, Cambridge, MA 02138, USA; Department of Zoology, University of Cambridge, Downing Street, Cambridge CB2 3EJ, UK; National Institute of Biological Resources, Incheon 37242, Republic of Korea; College of Veterinary Medicine and Veterinary Medical Research Institute, Jeju National University, Jeju 63243, Korea; Department of Chemistry and Chemical Biology, UNM Comprehensive Cancer Center, University of New Mexico, Albuquerque, NM 87131, USA

**Keywords:** Whale shark, Lifespan, body size, metabolic rate, neural genes

## Abstract

The endangered whale shark (*Rhincodon typus*) is the largest fish on Earth and is a long-lived member of the ancient Elasmobranchii clade. To characterize the relationship between genome features and biological traits, we sequenced and assembled the genome of the whale shark and compared its genomic and physiological features to those of 81 animals and yeast. We examined scaling relationships between body size, temperature, metabolic rates, and genomic features and found both general correlations across the animal kingdom and features specific to the whale shark genome. Among animals, increased lifespan is positively correlated to body size and metabolic rate. Several genomic features also significantly correlated with body size, including intron and gene length. Our large-scale comparative genomic analysis uncovered general features of metazoan genome architecture: GC content and codon adaptation index are negatively correlated, and neural connectivity genes are longer than average genes in most genomes. Focusing on the whale shark genome, we identified multiple features that significantly correlate with lifespan. Among these were very long gene length, due to large introns highly enriched in repetitive elements such as CR1-like LINEs, and considerably longer neural genes of several types, including connectivity, activity, and neurodegeneration genes. The whale shark’s genome had an expansion of gene families related to fatty acid metabolism and neurogenesis, with the slowest evolutionary rate observed in vertebrates to date. Our comparative genomics approach uncovered multiple genetic features associated with body size, metabolic rate, and lifespan, and showed that the whale shark is a promising model for studies of neural architecture and lifespan.

---

The relationships between body mass, longevity, and basal metabolic rate (BMR) across diverse habitats and taxa have been researched extensively over the last century, and led to generalized rules and scaling relationships that explain many physiological and genetic trends observed across the tree of life. While studies of endothermic aquatic mammals have shown that selection for larger body sizes is driven by the minimization of heat loss^1^, metabolic rate in ectothermic aquatic vertebrates is directly dependent on temperature, and decreased temperatures are correlated with decreased BMRs, decreased growth rates, longer generational times, and increased body sizes^2-4^. The whale shark (*Rhincodon typus*) is the largest extant fish, reaching lengths of 20 meters (m)^5^ and 42 tonnes (t) in mass^6^ and has a maximum lifespan estimated at 80 years^6^. Unlike the two smaller filter-feeding shark species (*Cetorhinus maximus*, *Megachasma pelagios*) that inhabit colder temperate waters with increased prey availability, whale sharks have a cosmopolitan tropical and warm subtropical distribution and have rarely been sighted in areas with surface temperatures less than 21°C^7-9^. However, recent GPS tagging studies have revealed that they routinely dive to mesopelagic (200-1,000 m) and bathypelagic (1,000-4,000 m) zones to feed, facing water temperatures of <4°C^10^. Observations of increased surface occupation following deeper dives led to the suggestion that thermoregulation is a primary driver for their occupation of the warmer surface waters^7,11^. Since larger body masses retain heat for longer periods of time, the large body mass of whale sharks may slow their cooling upon diving and maximize their dive times to cold depths, where food is abundant. Larger body mass could thus play a role in metabolic regulation.

Body size, environmental temperature, metabolic rate, and generation time are all correlated with variations in evolutionary rates^12,13^. Since many of these factors are interconnected, modeling studies have shown that observed evolutionary rate heterogeneity can be predicted by accounting for the impact of body size and temperature on metabolic rate^14^, suggesting these factors together drive the rate of evolution through their effects on metabolism. Consistent with these results, the coelacanth and elephant fish have the slowest reported evolutionary rates^15,16^. Moreover, genome size and intron size have also been linked to metabolic rate in multiple clades. Intron length varies between species and plays an important role in gene regulation and splice site recognition. In an analysis of amniote genomes, intron size was reduced in species with metabolically demanding powered flight and was correlated with overall reductions in genome size^17,18^. However, since most previous studies were limited by poor taxonomic sampling and absence of genome data for the deepest branches of the vertebrate tree, comprehensive comparative genomic analyses across gnathostomes are necessary to gain a deeper understanding of the evolutionary significance of the correlations between genome size, intron size and metabolic demands.

Here we sequenced and analyzed the genome of the whale shark and compared its genome and biological traits to those of 81 eukaryotic species, with a focus on gnathostomes such as fishes, birds, and mammals. In particular, we identified scaling relationships between body size, temperature, metabolic rates, and genomic features, and found general genetic and physiological correlations that span the animal kingdom. We also examined characteristics unique to the whale shark and its slow-evolving, large genome.

## The whale shark genome

The DNA of a *Rhincodon typus* individual was sequenced to a depth of 164× using a combination of Illumina short-insert, mate-pair, and TSLR libraries (Table S1 and S2), resulting in a 3.2 Gb genome with a scaffold N50 of 2.56 Mb (Tables S2, S5, and S6). A sliding window approach was used to calculate GC content and resulted in a genome-wide average of 42%, which is similar to the coelacanth and elephant fish (Fig. S2). Roughly, 50% of the whale shark genome is comprised of transposable elements (TEs), which were identified using both homology-based and *ab initio* approaches^19,20^. Of these, long interspersed nuclear elements (LINEs) made up 27% of the total TEs identified (Table S7). A combination of homology based and *ab initio* genome annotation methods^19,20^ resulted in a total of 28,483 predicted protein coding genes (Table S8).

## Correlation of physiological characteristics with genome features across 82 taxa

Body mass is intrinsically linked to physiological traits such as lifespan and basal metabolic rate (BMR)^21^. To better understand how genomic features correlate with physiological and ecological parameters such as body weight, lifespan, temperature, and metabolic rate, we compared the whale shark to 80 animals and yeast (Table S15-16) using physiological and genomic data (Fig.1, Fig. S3-6 and Table S16). Across the 81 animals examined, we found a strong positive correlation with significant *p*-values between the log transformed values for body weight and maximum lifespan (ρ = 0.79, Fig. 2A and Table S17) and BMR (ρ = 0.958, Fig. S9A, exponent B = 0.68, Fig. S25, and Table S17), consistent with previous reports^21^. Comparisons of physiological traits and genome characteristics across the 81 animals revealed several genetic features that also scaled with body weight. Among these, total gene length, intron length, and genome size all show a moderate statistical correlation with body mass, lifespan, and BMR (ρ = ~0.5) (Fig. 2B-E and Table S17). These results are consistent with previous findings of decreased intron size with increased metabolic rates. Furthermore, genome size and relative intron size are strongly correlated (ρ = 0.707) (Fig. 2B and Table S17), with the whale shark being a notable outlier. Moreover, genome size, measured as golden path length, scales with gene size, measured as summed length of exons and introns per gene (B = 1.32, Fig. S26). Additionally, we found that, unlike in bacteria^22^ and crustaceans^23^, genome size in Chordates scales positively with temperature (B = 0.77, Fig. S27).

**Fig. 1.**
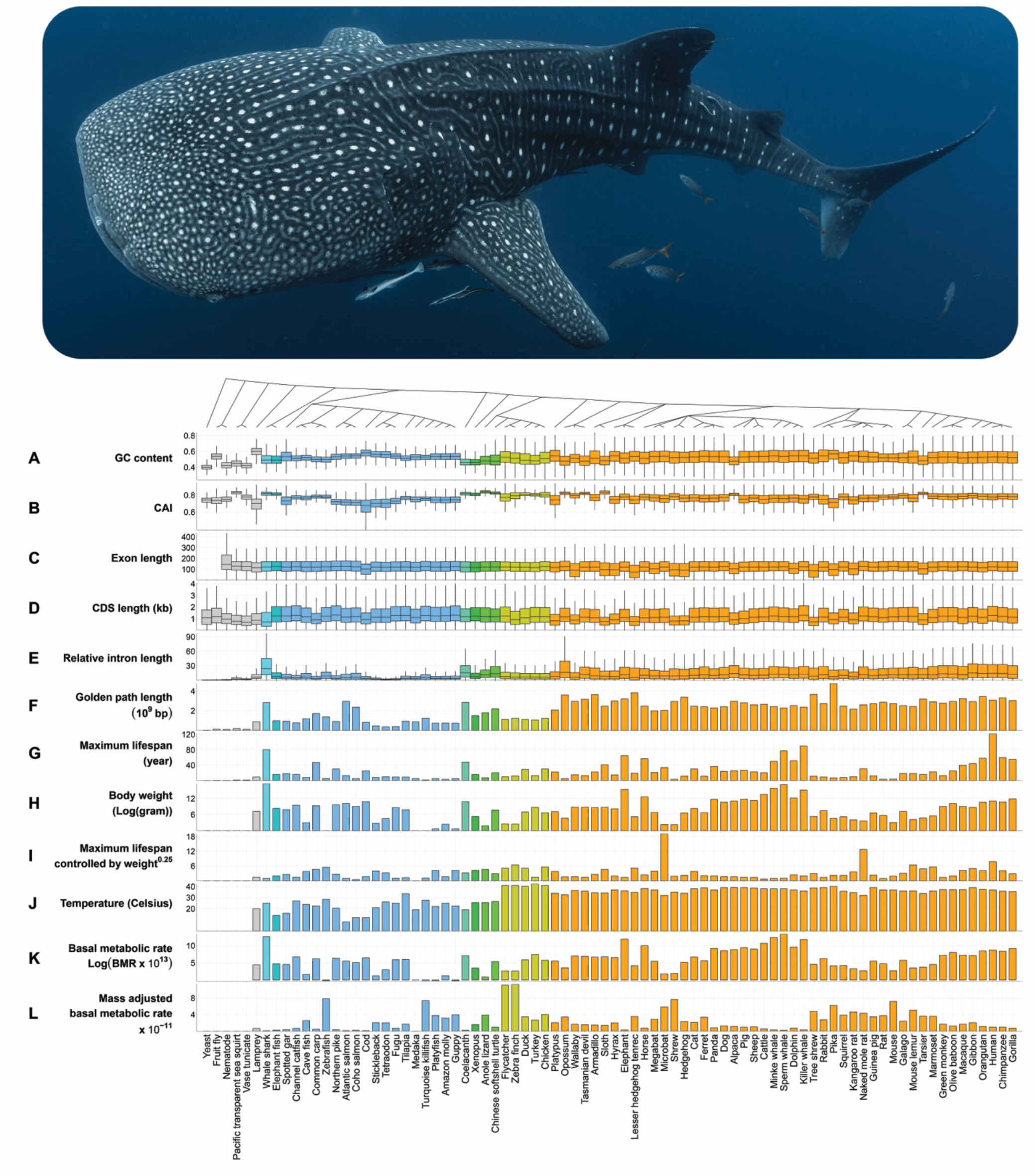
Comparative genomic analysis across 82 species reveals traits linked to lifespan and bodyweight. Top panel: image of a whale shark. Bottom panel: the phylogenetic tree was constructed using the NCBI common tree (https://www.ncbi.nlm.nih.gov/Taxonomy/CommonTree/wwwcmt.cgi) without divergence times. The second to the last rows show the following values in 82 species: five genomic contexts (**A-E**), golden path length (**F**), the maximum lifespan (**G**), body weight (**H**), maximum lifespan controlled by weight^0.25^ (**I**), body temperature (optimal temperature for cold-blooded animal) (**J**), basal metabolic rate (**K**), and basal metabolic rate adjusted by weight (**L**). The exon length (**C**) shows length of exons in coding region. Yeast and fruit fly exon length were removed due to their extremely long length (median exon lengths for yeast and fruit fly are 1,032 bp and 217 bp respectively). The relative intron length (**E**) was calculated by dividing the total intron length between first coding exon and last coding exon by the CDS length. The nine colors of boxes and bars indicate biological classification (gray: Hyperoartia, Ascidiacea, Chromadorea, Insecta and Saccharomycetes, turquoise: Chondrichthyes (the cyan color indicates whale shark), light blue: Actinopterygii, aquamarine: Sarcopterygii, dark green: Amphibia, light green: Reptilia, dark yellow: Aves, orange: Mammalia).

**Fig. 2.**
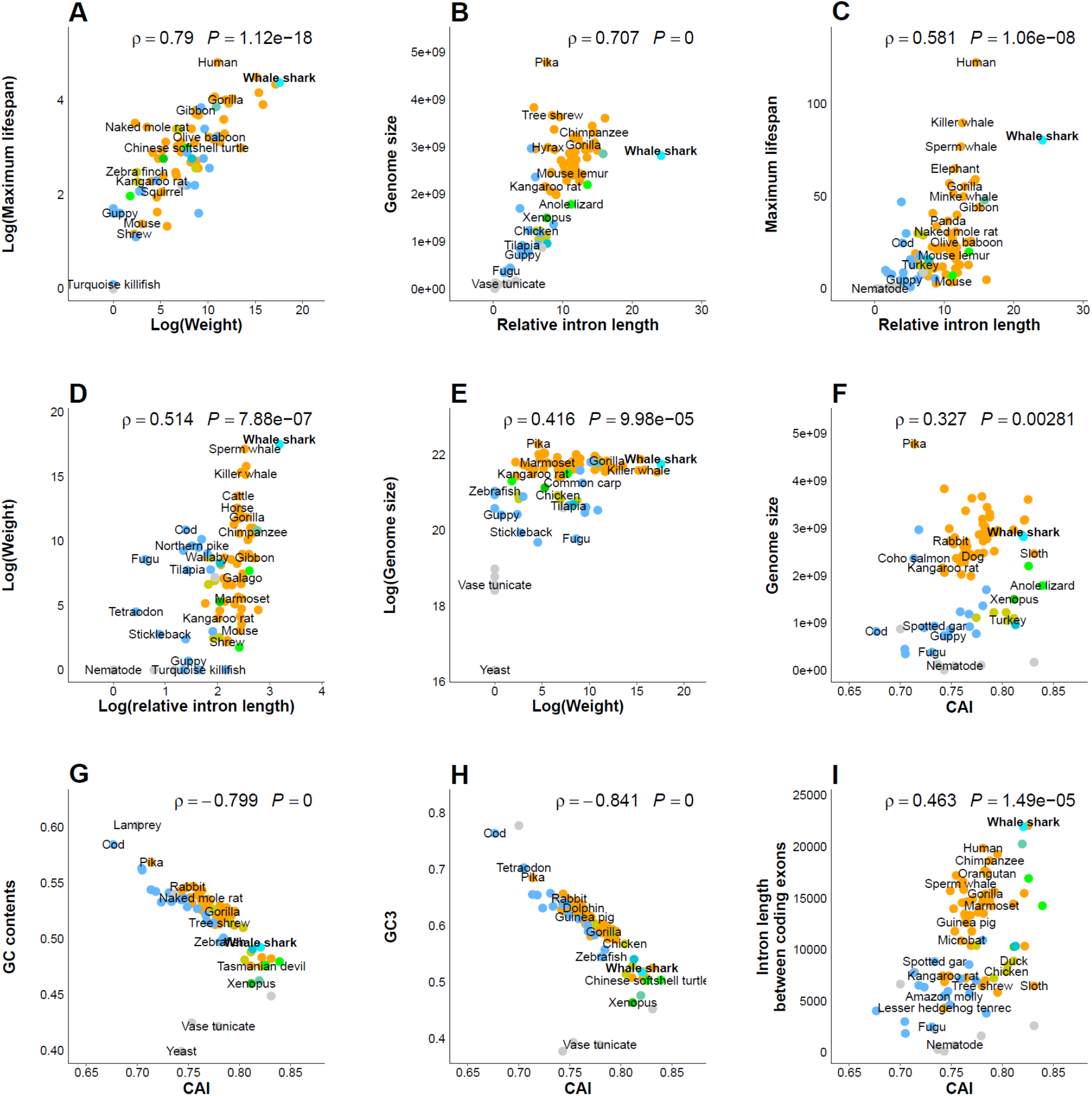
Scaling relationships between genomic and physiologic properties across 82 species. The properties on the x-axis and y-axis were used to calculate Spearman’s rank correlation coefficient for each plot. All *p*-values and rho values are shown at the top of each plot. Overlapping species names in the same layer were not plotted. The nine dot colors indicate biological classification (gray: Hyperoartia, Ascidiacea, Chromadorea, Insecta and Saccharomycetes, turquoise: Chondrichthyes (cyan is whale shark), light blue: Actinopterygii, aquamarine: Sarcopterygii, dark green: Amphibia, light green: Reptilia, dark yellow: Aves, orange: Mammalia).

Exon length is remarkably constant across animals, regardless of genome size or intron length (Fig. 1C and Fig. S4C). Early observations of this phenomenon across small numbers of taxa led to the suggestion that the splicing machinery imposes a minimum exon size while exon skipping begins to predominate when exons exceed ~500 nt in length^24^. Also of note is the tight correlation (ρ = 0.975) between the overall GC content and GC3, the GC content of the third codon position (Fig. S9B and Table S17), while both features are negatively correlated with the codon adaptation index (CAI) (ρ=−0.799 and ρ=−0.841, respectively; Fig. 2G-H and Table S17) in Eukaryota, and negatively correlated with the genome size in Mammalia (ρ = −0.434 and ρ = −0.473, respectively) (Table S17). These results are partially supported by previous research, which showed that GC3 is negatively correlated with body mass, genome size, and species longevity within 1,138 placental mammal orthologs^25^. However, our results using whole genome data do not support the GC3 correlation with body mass and longevity (ρ = 0.067 and ρ = 0.059; Table S17). Thus, exon and intron length may affect body mass and longevity through a strong association between GC content and coding sequence length^26^. Additionally, CAI and intron size are moderately positively correlated (ρ = 0.463; Fig. 2I and Table S17). Since the CAI and codon usage bias have an inverse relationship, this is consistent with the negative correlation between intron length and codon usage bias in multicellular organisms^27^.

## Whale shark longevity and genome characteristics

The allometric scaling relationships between longevity, mass, temperature, and metabolic rate are well established^21^, and the long lifespan of the whale shark can be explained by its large mass and the extremely low mass- and temperature-adjusted BMR (Fig. 1H and 1L). There has been considerable debate in the literature over the evolutionary causes and consequences of genome size, particularly as it relates to BMR. At 3.2 Gb, the whale shark has a genome that is significantly larger than those of other Chondrichthians (elephant fish), though both exon number and size are comparable. The whale shark is, however, a notable outlier, particularly among fish, for its long introns (Fig. 1E and S4E). Interestingly, the whale shark’s relative intron length (Fig. 1E and S4E) is significantly longer than any of the other 81 species (Fig. S5G and S6G). Analyses of single copy orthologous gene clusters did not reveal any large intron gains or losses in the whale shark (Fig. S10), though retrotransposon analyses revealed a significant expansion of CR1-like LINEs and Penelope-like elements in the introns (Fig. 3A and S11-15). The CR1-like LINEs are the dominant family of transposable elements (TEs) in non-avian reptiles and birds^28^. In the whale shark, the summed length of CR1-like LINE elements is 176 Mbp (Fig. S13C), which is eleven times longer than that of the anole lizard, a species known for expanded CR1-like LINEs. The total length of intronic repetitive elements is as great as in the opossum genome, known to be rich in repetitive elements^29^ (Fig. S12). In the whale shark genome, 38% of the CR1-like LINE, 39% of the CR1-Zenon like LINE, and 30% of the Penelope-like elements are located in intronic regions (Fig. S14). Strikingly, most genes (more than 88%) in the whale shark genome have the CR1-like LINE elements within their introns (Fig. S15) and 56% of genes also have LINE1 elements (Fig. S15). Thus, the whale shark’s relatively large genome and long introns are due to repetitive elements.

**Fig. 3.**
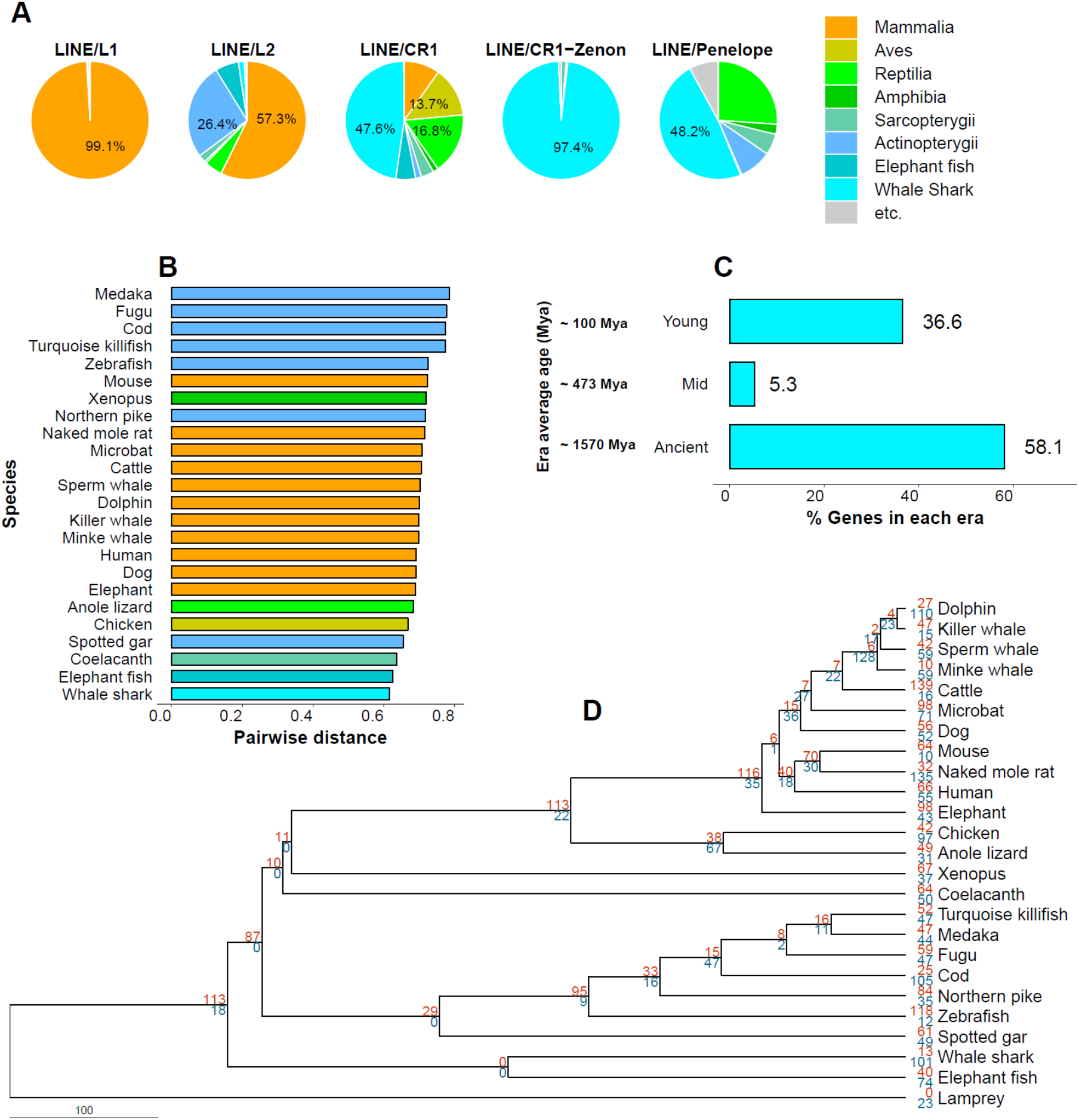
Repetitive elements, evolutionary rate model, and flow of genes in the whale shark genome. (**A**) Each pie chart summarizes the length of predicted intronic repetitive elements (labeled in the top of pie). Values from the 81 species (yeast excluded) were averaged across six Classes (Mammalia, Aves, Reptilia, Amphibia, Sarcopterygii, Actinopterygii), and the whale shark and elephant fish are listed separately (yeast was excluded from these analyses). (**B**) All pairwise distances from sea lamprey were calculated using the R-package ‘ape’ ^59^. The species were ordered by the pairwise distances. The eight bar colors indicate biological classification (turquoise: Chondrichthyes (the cyan color indicates whale shark), light blue: Actinopterygii, aquamarine: Sarcopterygii, dark green: Amphibia, light green: Reptilia, dark yellow: Aves, orange: Mammalia). (**C**) While most genes (~58%) in the whale shark genome are ancient, a few (~5%) are of intermediate age and a significant fraction (~37%) are relatively young. (**D**) Maximum likelihood phylogenetic tree. Red and blue numbers refer to the number of expanded and contracted gene families at each node compared to the common ancestor, respectively.

Previous research has shown that there is an association between codon usage and the evolutionary age of genes in metazoans^30^. Interestingly, two principal component analyses (PCA) of relative synonymous codon usage (RSCU) from 82 and 76 species (six species having distant codon usage patterns were excluded), respectively, revealed that the whale shark pattern of RSCU is most similar to that of the coelacanth; with well separated patterns of RSCU for each class (Fig. S16). While the whale shark genome has a relatively short exon length (smaller than that of 59 species), importantly, it has a smaller number of exons per gene than all but two species (the yeast and fruit fly have the smallest number of exons) (Fig. S3B and S4G). Thus, the whale shark CDS length is shorter than all but the yellow sea squirt genome (Fig. 1D and S4D).

## Evolutionary rate and historical demography

Analyses of the whale shark genome showed it is the slowest evolving vertebrate yet characterized. A relative rate test and two cluster analyses revealed that the whale shark has a slower evolutionary rate than those of the elephant fish and all other bony vertebrates examined, including coelacanth^16^ (Fig. 3B, S17 and Table S18-20). These results support the previous work which predicted a slow evolutionary rate in ectothermic, large-bodied species with relatively low body temperature (compared to similarly sized warm-blooded vertebrates)^14^. They are also consistent with previous studies of nucleotide substitution rates in elasmobranchs, which are significantly lower than those of mammals^31,32^.

A phylogenetic analysis of the 255 single-copy orthologous gene clusters from the whale shark and 24 other animal genomes (Fig. 3D) showed a divergence of the Elasmobranchii (sharks) and Holocephali (chimaeras) roughly 268 MYA and of the Chondrichthyes from the bony vertebrates about 457 MYA (Fig. 3D), consistent with previous estimates. To understand how many genes appeared in each evolutionary era within the whale shark genome, we evaluated the evolutionary age of whale shark protein-coding genes based on protein sequence similarity^33^. Grouping the whale shark genes into three broad evolutionary eras, we observed that while the majority (58%) of genes are ancient (older than 684 MYA), a few (~5%) are middle age (684-199 MYA), and many (36%) are young (199 MYA to present) (Fig. 3C). Normalizing the number of genes by evolutionary time suggests that gene turnover is highest near the present time (Fig. S18). Examining the age of genes shows many genes are ancient (PS 1) and many genes appear very young (PS 20) (Fig. S19), though the large number of young PS 20 genes may in part reflect the paucity of closely related species with fully sequenced genomes. These results highlight both the conservation of a large part of the genome as well as the innovative potential of the whale shark genome, since many new genes appeared within the last 200 million years.

## Gene family expansions and contractions in the whale shark

Gene family expansion and contraction analyses across 25 species identified 101 contracted gene families in the whale shark. Of these, nucleosome assembly (GO:0006334) and chromatin assembly (GO:0031497) were significantly decreased in the whale shark compared to the Chondrichthyes common ancestor (Table S21A). Interestingly, the whale shark genome has a smaller number of histone gene families (H1, H2A and H2Bs) than other bony fishes and mammals (Fig. S20). This small number of histone gene families, especially the H1 family which encodes the linkers important for higher order chromatin structures, may be related to the long length of whale shark introns^34^. We also identified 13 expanded gene families that are enriched for several metabolic pathways, including fatty acid metabolism, along with neurogenesis and nervous system development, and cardiac conduction system development (Table S21B).

## Gene length of neural genes and correlation with physiological features

Gene length has recently emerged as an important feature of neural genes, as long genes are preferentially expressed in neural tissues and their expression is under tight transcriptional and epigenetic control^35^. Within the 81 animal species, we compared the dimensions of average genes with those of ten categories of neural genes (neuronal connectivity, cell adhesion, olfactory receptors, ion channels, unfolded protein response associated genes, neuronal activity and memory, neuropeptides, homeobox genes, synaptic genes, and neurodegeneration) (Fig. 4A and S21). Interestingly, we found that neuronal connectivity genes are longer than average genes in most vertebrates, with the length increase being significant in whale shark and most mammals, as well as in coelacanth and platypus (Fig. 4A and S22A). Surprisingly, we found that neural genes are scaled to average genes with an exponent greater than 1 (B = 1.038, Fig. S28), with the whale shark showing an extreme lengthening of neural genes. Moreover, we found that cell adhesion, ion channels, homeobox genes, and neurodegeneration genes are increased in length in the whale shark (Fig. 4B). Thus, the organization of whale shark neural genes may reflect the need to maintain the shape, activity, identity, and resistance to neurodegeneration in a body that is both very large and long-lived. Finally, neuronal functions are enriched in long genes in more than 60 species (Fig. 4C and Additional File 1).

**Fig. 4.**
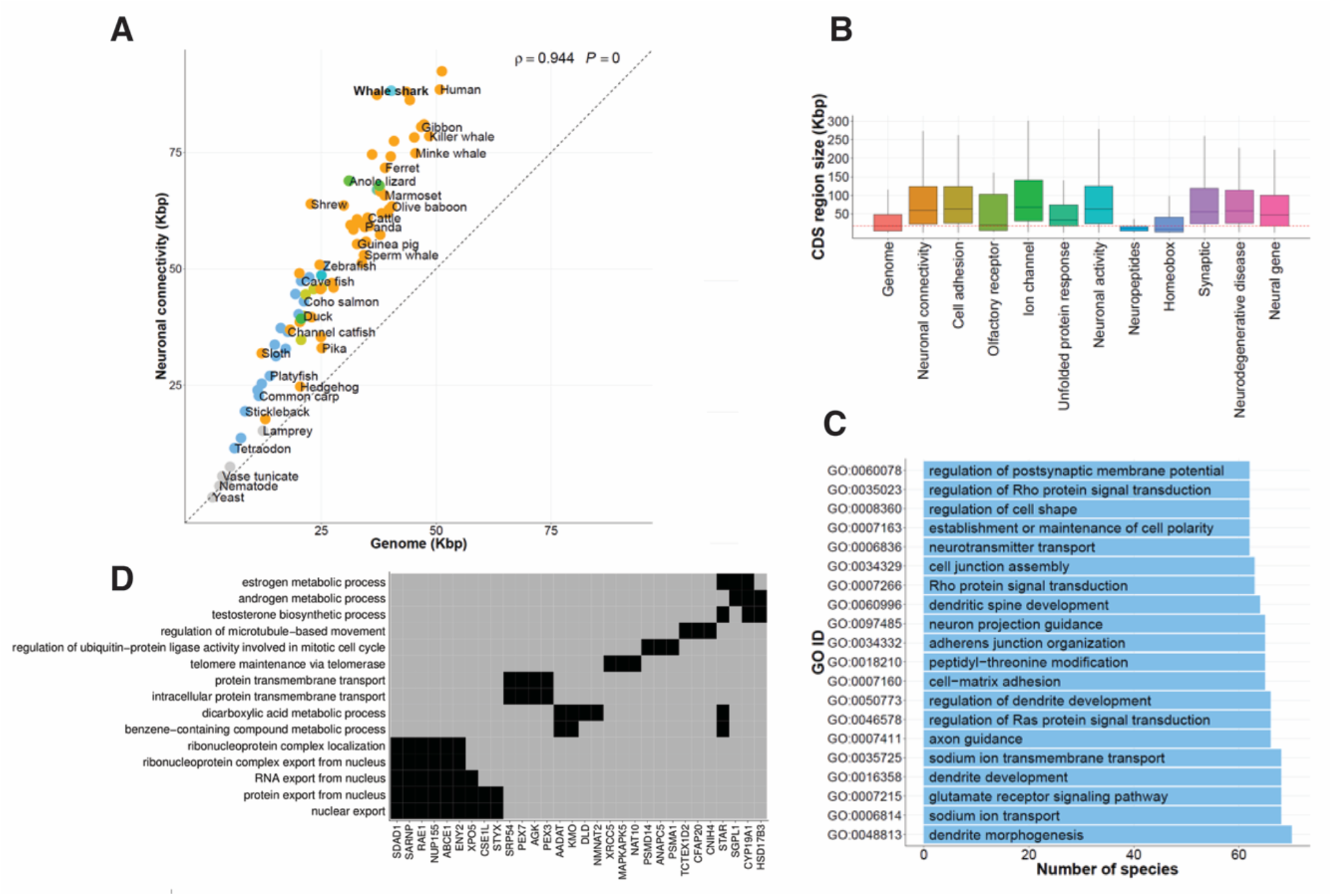
The relationship between gene length and neural genes, and single-copy orthologous gene families with correlations between gene length and maximum lifespan, weight, and BMR. (**A**) Neuronal connectivity genes are longer in 81 species (yeast excluded). The x- and y-axes show the average gene length and the gene length of neuronal connectivity-related genes, respectively. The dashed diagonal line represents ‘y = x’. Spearman’s rho correlation coefficient and *p-*value are shown in top right corner of the plot. (**B**) Of the 12 categories of neural genes we analyzed in the whale shark genome, several are longer than average genes. (**C**) Most common GO terms are relevant to neural function. GO terms are shown based on the number of species they were found in, and were computed with Gene Set Enrichment Analysis (GSEA). (**D**) Enriched GO functions in single-copy orthologous gene families in which relative intron length positively correlates with maximum lifespan. For each GO term, black boxes indicate representative human gene symbols representative of the family.

To determine whether physiological traits and genomic features are linked, we examined the correlation of gene size and maximum lifespan, body weight, and BMR (Fig. 2A-F). In 155 gene families, we found that gene length was significantly correlated to maximum lifespan, body weight, and BMR. Gene ontology analyses of this gene group showed statistical enrichment of biological processes such as telomere maintenance (GO:0007004: *XRCC5, MAPKAPK5*, and *NAT10*) and RNA and protein export from nucleus (GO:0006405 and GO:0006611: *SDAD1, SARNP, RAE1, NUP155, ABCE1, ENY2, XPO5, CSE1L* and *STYX;* Fig. 4D, Tables S22 and S23), both of which are associated with longevity and cancer^36,37^. Of the genes in which gene length is associated with lifespan, NUP210 (nucleoporin 210) and VWF (von Willebrand factor) are both associated with longevity^38^ (Fig. S24A and Table S24). Moreover, the genes correlated to BMR include *SNX14*, which is linked to protein metabolism and whose deficiency causes ataxia and intellectual disability^39^ (Fig. S24B and Table S24). The only gene previously correlated with body mass (*COX5B*; the terminal enzyme of the mitochondrial respiratory chain) is a subunit of Complex IV and is essential to energy production in the cell and ultimately to aging^40^ (Fig. S24C and Table S24). Taken together, these results suggest that there is an evolutionary relationship between gene size and physiological traits size such as body size, metabolic rate, and lifespan. This holds particularly among genes whose functions are essential for living long lives, such as telomere maintenance and energy production.

## Conclusions

We sequenced and assembled the genome of the whale shark (*Rhincodon typus*), an endangered species that is the largest extant fish on Earth. Its relatively large 3.2 Gb genome is the slowest evolving vertebrate genome found to date, and has a striking amount of CR1-like LINE transposable elements. In most genomes, we found that major genomic traits, including intron length and gene length, correlate with body size, temperature, and lifespan, and that GC content and codon adaptation index are negatively correlated. Unexpectedly, we found that neural connectivity genes are substantially longer than average genes. In the whale shark genome, specifically, we found that introns are longer than in most other species due to the presence of repetitive elements and that neural genes of several types, including neurodegeneration genes, are much longer than average genes of species with long lifespans. These results show the power of the comparative evolutionary approach to uncover both general and specific relationships that reveal how genome architecture is shaped by size and ecology.

## Methods

### Sample preparation and sequencing

Genomic DNA was isolated from heart tissue acquired from an approximately seven years old, 4.5 meter deceased male whale shark from the Hanwha Aquarium, Jeju, Korea. DNA libraries were constructed using a TruSeq DNA library kit for the short-read libraries and a Nextera Mate Pair sample prep kit for the mate pair libraries. Sequencing was performed using the Illumina HiSeq2500 platform. Libraries were sequenced to a combined depth of 164× (Tables S1 and S2).

### Genome assembly and annotation

Reads were quality filtered (Table S3) and the error corrected reads from the short insert size libraries (<1 Kb) and mate pair libraries (>1 Kb) were used to assemble the whale shark genome using SOAPdenovo2^41^. As the quality of assembled genome can be affected by the *K*-mer size, we used multi-K-mer value (minimum 45 to maximum 63) with the ‘all’ command in the SOAPdenovo2 package^41^. The gaps between the scaffolds were closed in two iterations with the short insert libraries (<1 Kb) using the GapCloser program in the SOAPdenovo2 package^41^. We then aligned the short insert size reads to the scaffolds using BWA-MEM^42^ with default options. Variants were identified using SAMtools^43^. Since at least one of alleles from the mapped reads of same individual as reference should be presented in the assembly, we corrected erroneous bases where both alleles were not present in the assembly by substituting the first variant alleles. Finally, we mapped the Illumina TruSeq synthetic long reads (TSLR) to the assembly and corrected the gaps covered by the synthetic long reads to reduce erroneous gap regions in the assembly (Table S5).

The GC distribution of the whale shark genome was calculated using a sliding window approach. We employed 10 Kb sliding windows to scan the genome to calculate the GC content. Tandem repeats were predicted using the Tandem Repeats Finder program (version 4.07)^44^. Transposable elements (TEs) were identified using both homology-based and *ab initio* approaches. The Repbase database (version 19.02)^45^ and RepeatMasker (version 4.0.5)^19^ were used for the homology-based approach, and RepeatModeler (version 1.0.7)^20^ was used for the *ab initio* approach. All predicted repetitive elements were merged using in-house Perl scripts. Two candidate gene sets were built to predict the protein coding genes in the whale shark genome; using AUGUSTUS^46^ and Evidence Modeler (EMV)^47^, respectively (Supplementary Text 1.7).

### Genomic context calculations

From the 82 species (Table S15), we computed the following genomic factors: GC3 (GC content at third codon position), CAI (codon adaptation index), number and length of coding exon(s), and relative intron length between first and last exon (or coding exon). CDS sequences with premature stop codons and lengths that were multiples of three were excluded. The relative intron length was calculated by dividing the total intron length between first and last exon (or coding exon) by the CDS length (or mRNA length). GC3 was computed from concatenated third codon nucleotides using the canonical method^48^. We measured relative synonymous codon usage (RSCU) using the method from Sharp *et al.*^49^ and the codon adaptation index (CAI) in a CDS using Sharp and Li’s method^50^ for each of the 82 species. The principle component analysis (PCA) on RSCU was performed using the R packages (version 3.3.0)^51^ ggplot2^52^ and ggfortify^53^.

### Orthologous gene family clustering

To identify orthologous gene families among the whale shark and the other 82 species, we downloaded all pair-wise reciprocal BLASTP results using the ‘peptide align feature’ in the Ensembl comparative genomics resources^54^ (release 86). To generate pair-wise orthologous that were not available in the Ensembl resources, we performed reciprocal BLASTP^55^ with the ‘-evalue 1e-05-seg no-max_hsps_per_subject 1-use_sw_tback’ options. From the pair-wise reciprocal BLASTP results among the 82 species, we generated similarity matrixes by connecting possible orthologous pairs. To constrain the computational load, we did not join additional nodes when the number of node was bigger than 1500. The normalized weights for the similarity matrix were calculated using the OrthoMCL approach^56^. We identified orthologous gene families by using an in-house C++ script based on the MCL clustering algorithm^57^, with inflation index 1.3. A total of 1,461,312 genes were assigned to 225,530 clusters, including 192,174 of singletons.

### Gene age estimation

Phylostratigraphy uses BLASTP-scored sequence similarity to estimate the minimal age of every protein-coding gene. The NCBI non-redundant database is queried with a protein sequence to detect the most distant species in which a sufficiently similar sequence is present and posit that the gene is at least as old as the age of the common ancestor^33^. Using NCBI for every species, the timing of lineage divergence events is estimated with TimeTree^58^. To facilitate detection of protein sequence similarity, we use the e-value threshold of 10^−3^. We evaluate the age of all proteins with length equal or greater than 40 amino acids. First, we count the number of genes in each phylostratum, from the most ancient (PS 1) to the most recent (PS 20). Most genes are ancient (PS 1-2) and a substantial number appear young (PS 20) (Fig. S19). Second, to understand broad evolutionary patterns, we aggregate the counts from several phylostrata into three broad evolutionary eras: ancient (PS 1-7, cellular organisms to Deuterostomia, 4,204 Mya - 684 Mya), middle (PS 8-14, Chordata to Selachii, 684 Mya - 199 Mya) and young (PS 15-20, Galeomorpha to *Rhincodon typus*, 199 Mya to present). To understand the gene flow per time unit, we normalized the number of genes by the age and the duration of the evolutionary era.

### Correlation tests in orthologous gene families

From the 82 species, we selected 6,929 single-copy orthologous gene families which are found in at least 40 species to calculate the correlation between gene length, i.e., exon + intron length between first and last coding exon and three physiological properties (the maximum lifespan, body weight, and BMR). We identified gene families which had significant correlations between the gene length and the maximum lifespan (2,882 genes), body mass (2,193 genes), and the BMR (2,627 genes) by Spearman’s rho correlation coefficient and Benjamini & Hochberg adjustment (adjust *p-*value ≤ 0.01). All these gene families were subject to alignment filtering criterion of containing more than 50% of conserved exon-exon boundaries (intron position) in their CDS alignments. This step reduces the effect of gene length change due to intron gain or loss and increases the accuracy of multiple sequence alignments (Fig. S23). Finally, we acquired four sets of correlated gene families between the gene length and the three properties: 1) 25 gene families with the maximum lifespan only (Table S24), 2) one gene family with the body weight only (Table S24), 3) seven gene families with the BMR only, and 4) 155 gene families with all three physiological properties (Table S23).

## Acknowledgements

We thank Dr. Mark Erdmann for generously providing the whale shark photograph used in Figure

1. This work was supported by the Genome Korea Project in Ulsan (800 genome sequencing) Research Fund (1.180017.01) of UNIST (Ulsan National Institute of Science & Technology) and the Genome Korea Project in Ulsan (200 genome sequencing) Research Fund (1.180024.01) of UNIST (Ulsan National Institute of Science & Technology). V.L. thanks Marc W. Kirschner and gratefully acknowledges funding support from the National Institutes of Health of USA (R01 HD073104 and R01 HD091846 to M.W.K.).

## Author contributions

J.B. conceived and planned the project. J.A.W., Y.S.C., S.G.P, and J.B. coordinated the project. J.A.W., V.L., S.G.P., S.J., J.S.E., and J.B. wrote the manuscript. W.H.H., S.W.K., S.K., Y.S.C., H.M.K., and S.K. prepared the samples and performed the experiments. S.G.P., J.A.W., V.L., S.J., H.M.K., Y.J., Y.B., A.K. and J.J. performed in-depth bioinformatics and evolutionary analyses. S.G.P., J.A.W., V.L., S.J., H.M.K., Y.J., Y.B., J.J., S.W.K., W.H.H., S.L., Y.S.C., A.K., A.M., S.K., J.S.E., J.B., and G.M.C. reviewed the manuscript and discussed the work.

## Data and materials availability

The whale shark whole genome project has been deposited at DDBJ/ENA/GenBank under the accession QPMN00000000. The version described in this paper is version QPMN01000000. DNA sequencing reads have been uploaded to the NCBI Read Archive (SRP155581).

## 1. Whale shark genome sequencing and assembly

### 1.1 DNA sample preparation and sequencing

Genomic DNA was extracted from the heart tissue of a 4.5-meter, seven year old dead male whale shark (*Rhinocodon typus*, from the Hanwha Aquarium, Jeju, Republic of Korea). DNA libraries were constructed using a TruSeq DNA library kit for the short-read libraries and a Nextera Mate Pair sample prep kit for the mate pair libraries. Libraries were sequenced using the Illumina HiSeq2500 platform. We obtained roughly 164× of paired-end short reads with varying insert sizes including mate pair (Table S1) and 848,425 TSLRs (Table S2).

**Table S1.**
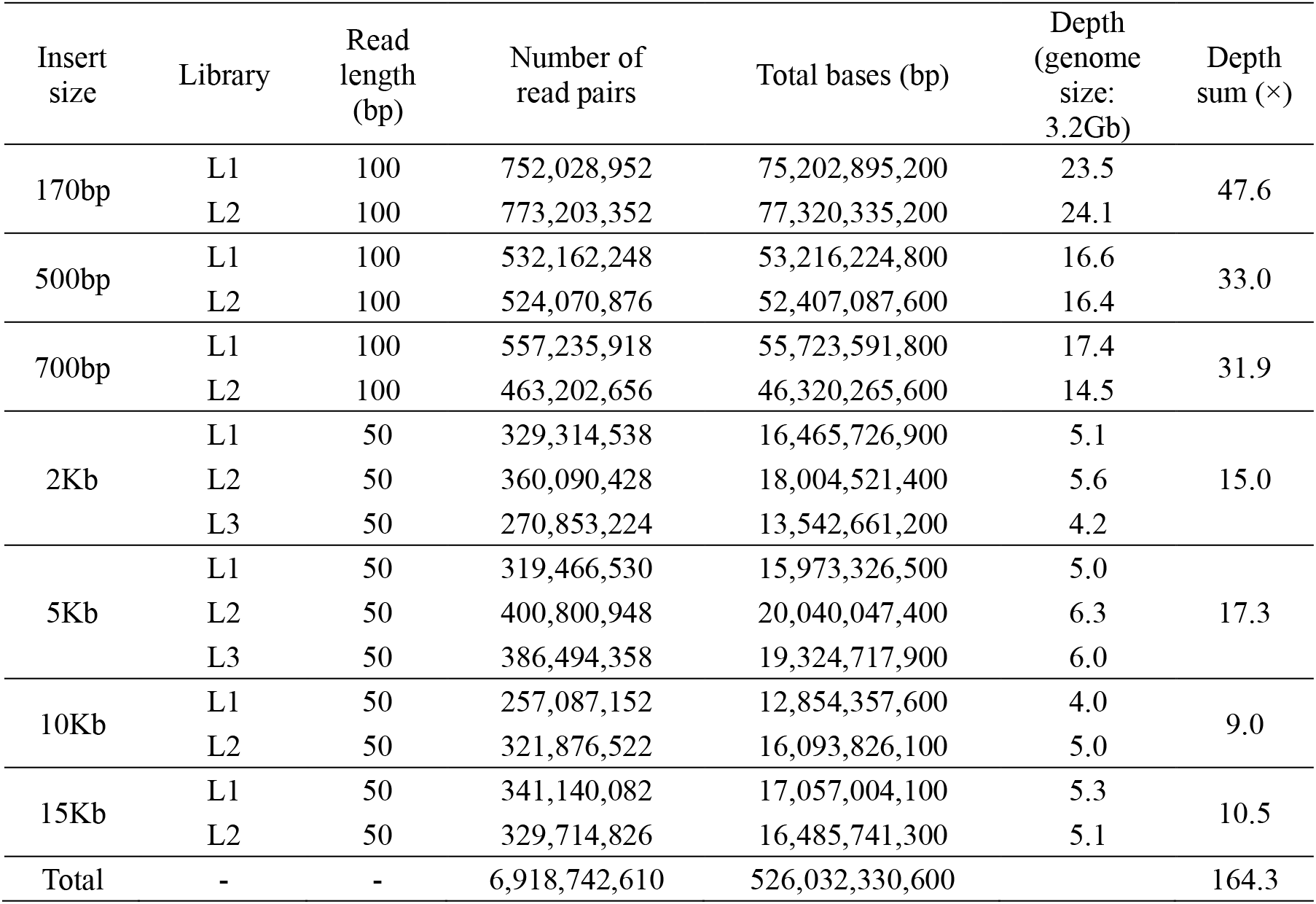
Short insert and mate pair library sequencing statistics

| Insert size | Library | Read length (bp) | Number of read pairs | Total bases (bp) | Depth (genome size: 3.2Gb) | Depth sum (×) |
| --- | --- | --- | --- | --- | --- | --- |
| 170bp | L1 | 100 | 752,028,952 | 75,202,895,200 | 23.5 | 47.6 |
|  | L2 | 100 | 773,203,352 | 77,320,335,200 | 24.1 |  |
| 500bp | L1 | 100 | 532,162,248 | 53,216,224,800 | 16.6 | 33.0 |
|  | L2 | 100 | 524,070,876 | 52,407,087,600 | 16.4 |  |
| 700bp | L1 | 100 | 557,235,918 | 55,723,591,800 | 17.4 | 31.9 |
|  | L2 | 100 | 463,202,656 | 46,320,265,600 | 14.5 |  |
| 2Kb | L1 | 50 | 329,314,538 | 16,465,726,900 | 5.1 | 15.0 |
|  | L2 | 50 | 360,090,428 | 18,004,521,400 | 5.6 |  |
|  | L3 | 50 | 270,853,224 | 13,542,661,200 | 4.2 |  |
| 5Kb | L1 | 50 | 319,466,530 | 15,973,326,500 | 5.0 | 17.3 |
|  | L2 | 50 | 400,800,948 | 20,040,047,400 | 6.3 |  |
|  | L3 | 50 | 386,494,358 | 19,324,717,900 | 6.0 |  |
| 10Kb | L1 | 50 | 257,087,152 | 12,854,357,600 | 4.0 | 9.0 |
|  | L2 | 50 | 321,876,522 | 16,093,826,100 | 5.0 |  |
| 15Kb | L1 | 50 | 341,140,082 | 17,057,004,100 | 5.3 | 10.5 |
|  | L2 | 50 | 329,714,826 | 16,485,741,300 | 5.1 |  |
| Total | - | - | 6,918,742,610 | 526,032,330,600 |  | 164.3 |

**Table S2.** Illumina TruSeq synthetic long read (TSLR) sequencing statistics

|  | All | > 1,500bp only |
| --- | --- | --- |
| Number of sequences | 848,425 | 588,325 |
| Number of bases (bp) | 3,774,313,129 | 3,547,450,453 |
| N50 (bp) | 8,407 | 8,750 |
| The largest length (bp) | 21,924 | 21,924 |
| Average length (bp) | 4,448 | 6,029 |

### 1.2 Raw data QC trimming and filtering

Low quality or contaminated reads were removed using the following filtering criteria:

1) PCR duplications (the reads were considered duplications if both paired end reads are identical).
2) Reads containing adapters. Left = “*GATCGGAAGAGCACACGTCTGAACTCCAGTCAC*” Right = “*GATCGGAAGAGCGTCGTGTAGGGAAAGAGTGT*”
3) Reads which had more than 5% ambiguous bases (N).
4) Reads with an average base quality below 20 (<Q20).
5) Reads which had junction adapters in the mate-pair libraries. Left = “*CTGTCTCTTATACACATCT*” Right = “*AGATGTGTATAAGAGACAG*”
6) Low-quality ends were trimmed for the short-insert libraries (2bp of 5’-end and 8bp of 3’-end).

Roughly 120× depth of coverage remained after filtering (Table S3).

**Table S3.** Post QC short insert and mate pair library sequencing statistics

| Insert size | Read length (bp) | Total Reads | Total Bases (bp) | Remained depth<br>(X, genome size:<br>3.2Gb) |
| --- | --- | --- | --- | --- |
| 170bp | 90 | 1,436,964,768 | 129,326,829,120 | 40.38963675 |
| 400bp | 90 | 561,405,924 | 50,526,533,160 | 15.77977543 |
| 500bp | 90 | 958,715,504 | 86,284,395,360 | 26.9471958 |
| 700bp | 90 | 830,451,564 | 74,740,640,760 | 23.34200375 |
| 2kb | 49 | 260,885,666 | 12,783,397,634 | 3.992340881 |
| 5kb | 49 | 160,898,212 | 7,884,012,388 | 2.462229985 |
| 10kb | 49 | 111,938,498 | 5,484,986,402 | 1.712998068 |
| 15kb | 49 | 103,019,726 | 5,047,966,574 | 1.576513842 |
| Total | - | 4,424,279,862 | 372,078,761,398 | 116.2026945 |

### 1.3 Estimation of genome size using K-mer analysis

The size of the whale shark genome was estimated by *K*-mer analysis (*K*=23) using the KmerFreq_HA command of the SOAPec program in the SOAPdenovo2 package^1^ (Figure S1). The genome size was calculated by dividing the total number of *K*-mers by a peak depth of *K*-mers (Table S4). The whale shark genome size was estimated to be approximately 3.14 Gb. Prior to genome assembly, the sequencing errors in the filtered reads were corrected using the *K*-mer frequency (*K*=23) information and the Corrector_HA command of the SOAPec program^1^ with a three-depth criterion for low-frequency *K*-mer cutoffs.

**Figure S1.**
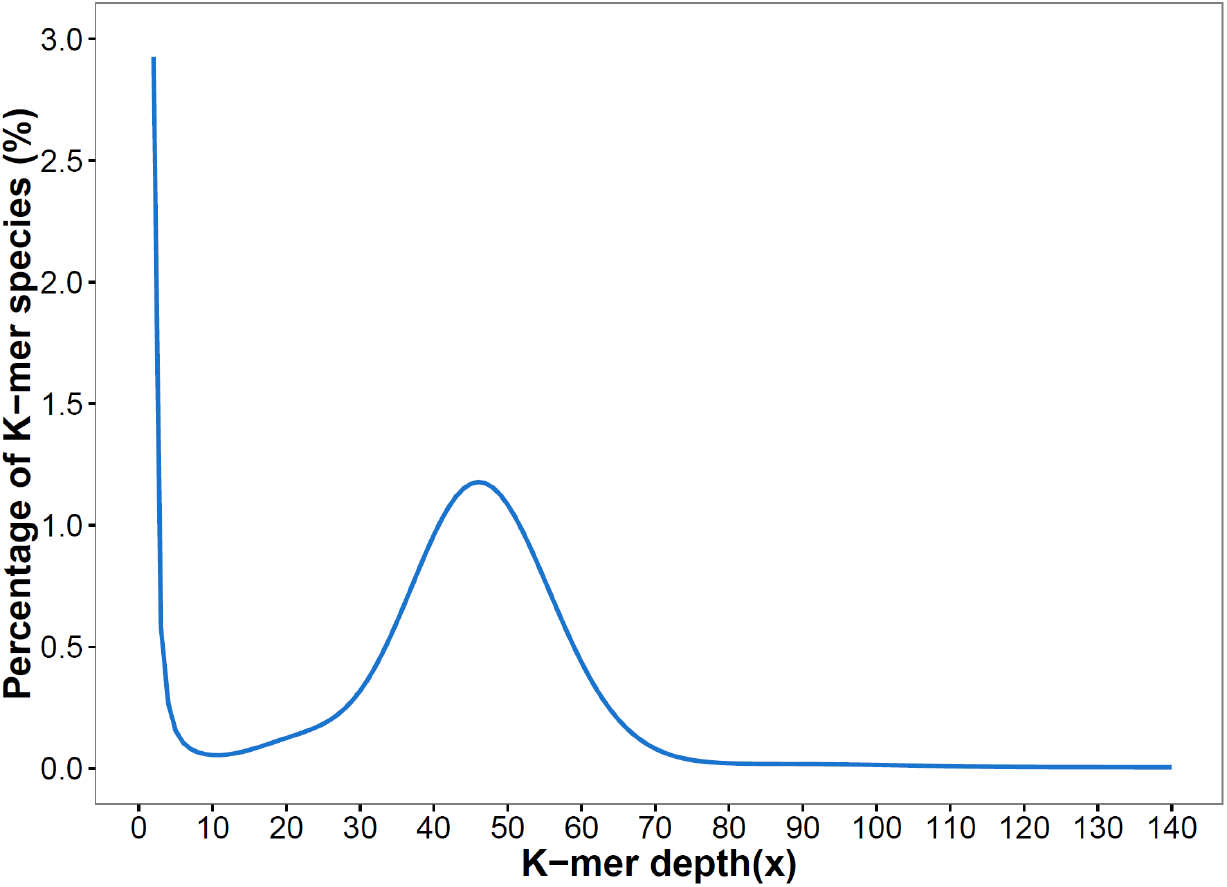
*K*-mer distribution frequency in the error-corrected reads, based on a 23-mer. The x-axis represents depth, and the y-axis represents proportion of *K*-mer species, as calculated by the frequency at a certain depth divided by the total frequency at all depths.

**Table S4.** Estimation of the whale shark genome size based on *K*-mer frequency using the error corrected reads

| **$K$-mer size** | **Total number of**<br>**$K$-mer** | **$K$ depth of**<br>**peak** | **Estimated genome**<br>**size** |
| --- | --- | --- | --- |
| 23 | 144,222,502,823 | 46 | 3,135,271,800 |

### 1.4 Genome assembly

The whale shark genome was assembled using the error-corrected reads from the short insert and mate pair libraries (>1 Kb) using SOAPdenovo2^1^. As the quality of assembled genome can be affected by the *K*-mer size, we used multi-*K*-mer values (minimum 45 to maximum 63) using the ‘all’ command in the SOAPdenovo2 package^1^. The gaps between the scaffolds were closed in two iterations with the short insert libraries (<1 Kb) using the GapCloser program in the SOAPdenovo2 package^1^. We then aligned the short insert size reads to the scaffolds using BWA-MEM^2^ with default options. Variants were identified using SAMtools^3^. At least one of the alleles from the self-mapping results should be the same as the reference. Thus, the erroneous bases of the assembly which are different from both alleles of the self-mapping results were changed to one of the alleles. We mapped the Illumina TruSeq synthetic long reads (TSLR) to the assembly and corrected the gaps covered by the synthetic long reads to reduce erroneous gap regions in the assembly (Table S5). The final length of the assembly is roughly 3.2 Gb with a scaffold N50 of 2.56 Mb and a contig N50 of 36 Kb (Table S6).

**Table S5.** Assembly statistics after removing erroneous gap regions with the TSLRs

|  | Before substitution | After substitution | Difference |
| --- | --- | --- | --- |
| Number of gaps | 188,018 | 172,567 | -15,451 |
| Number of monomer-gap | 127,356 | 112,865 | -14,491 |
| Bases (bp) | 3,202,752,364 | 3,201,980,496 | -771,868 |
| Sequences | 3,305,708 | 3,305,708 | 0 |

**Table S6.**
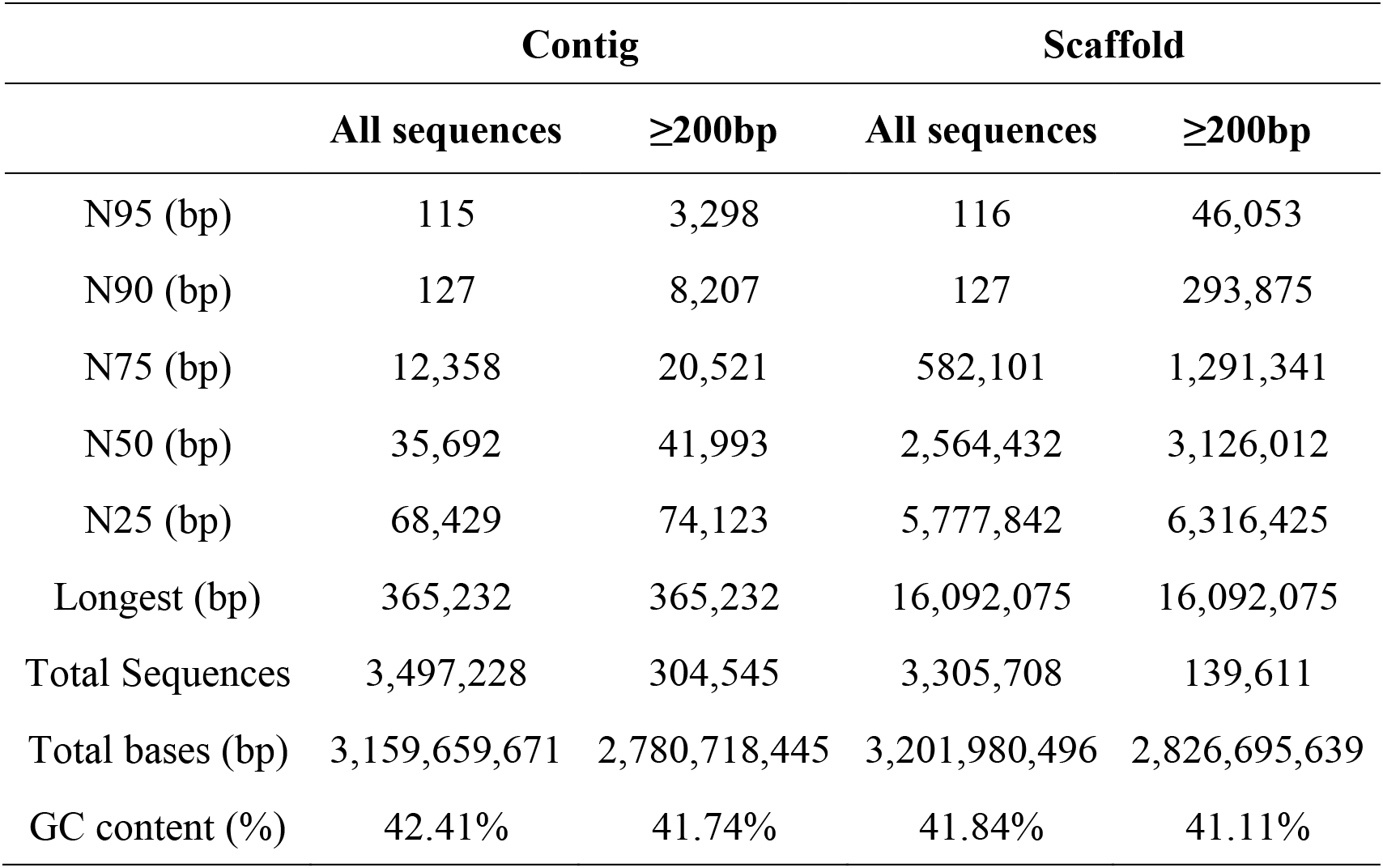
Final *de novo* assembly statistics

### 1.5 GC-content of whale shark genome

The GC distribution of the whale shark genome was calculated using a sliding window approach. We employed 10 Kb sliding windows to scan the genome to calculate the GC content. The average GC content of the whale shark is 41.6%, which is similar to that of coelacanth and elephant fish (Figure S2).

**Figure S2.**
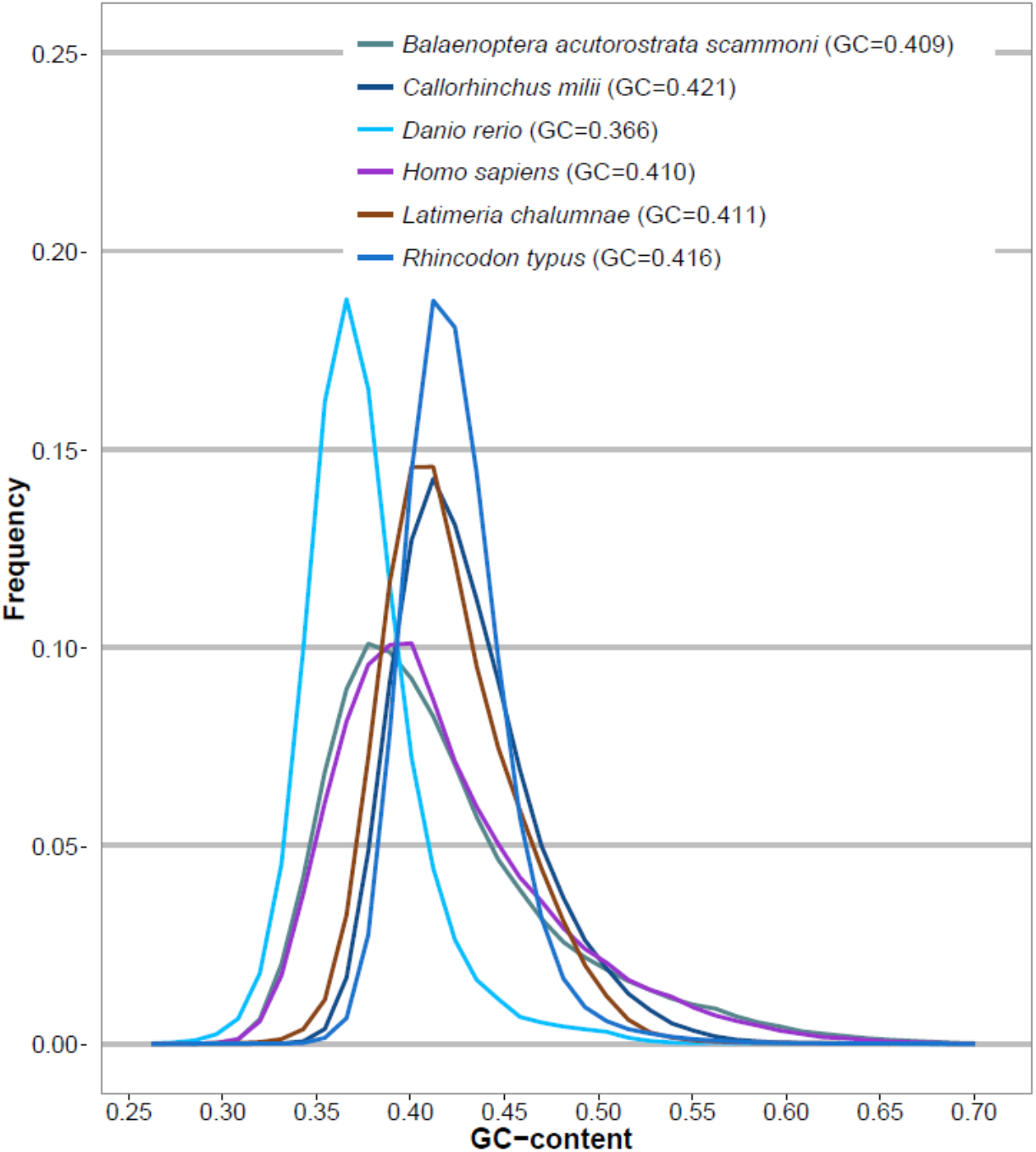
Genome-wide GC distribution. The x-axis represents GC content and the y-axis represents the proportion of the specified GC content. ‘GC’ in the legend indicates whole genome GC content of each species.

### 1.6 Annotation of repetitive elements

Tandem repeats were predicted using the Tandem Repeats Finder program (version 4.07)^4^. Transposable elements (TEs) were identified using both homology-based and *ab initio* approaches. The Repbase database (version 19.02)^5^ and RepeatMasker (version 4.0.5)^6^ were used for the homology-based approach, and RepeatModeler (version 1.0.7)^7^ was used for the *ab initio* approach,. All predicted repetitive elements were merged using in-house Perl scripts. In total, 49.55% of the whale shark genome is made of TEs (Table S7).

**Table S7.** Repetitive element statistics for the whale shark genome

| Type | *Ab initio*-based (bp) | Homology-based (bp) | Total (bp) | Percentage of genome |
| --- | --- | --- | --- | --- |
| DNA | 65,075,457 | 22,286,842 | 86,564,210 | 2.70% |
| LINE | 781,235,803 | 260,999,963 | 861,138,326 | 26.89% |
| LTR | 101,363,964 | 912,079 | 101,919,539 | 3.18% |
| Low complexity | 415,435 | 0 | 415,435 | 0.01% |
| SINE | 7,020,248 | 3,595,973 | 10,614,972 | 0.33% |
| Satellite | 7,341,297 | 18,859 | 7,350,548 | 0.23% |
| Simple repeat | 67,281,471 |  | 67,281,471 | 2.10% |
| Tandem repeat* |  |  | 249,559,685 | 7.79% |
| Unknown | 519,673,351 | 17,679 | 519,689,768 | 16.23% |
| Unspecified | 6,777,305 |  | 6,777,305 | 0.21% |
| **Total** | **1,508,223,137** | **287,829,289** | **1,586,543,783** | **49.55%** |

\*Tandem Repeat was separately predicted using TRF program.

### 1.7 Annotation of protein coding genes

Two candidate gene sets were built to predict the protein coding genes in the whale shark genome; using AUGUSTUS^8^ and Evidence Modeler (EMV)^9^, respectively.

1) For the AUGUSTUS^8^ prediction, we used both homology-based and *ab initio* approaches. For the homology-based gene prediction, homologous genes were identified by aligning the protein sequences from the elephant fish, zebrafish, medaka, human, mouse, and minke whale (from NCBI) to the cartilage fish protein database (from Uniprot) using GeneBlastA^10^ with e-value cutoff 10^−5^. Homologous genes with less than 40% coverage were filtered out. Homology-based gene models were constructed using Exonerate^11^. With the Homology-based gene model and three hints: cartilage fishes’ EST sequences from NCBI, transcriptomic hint from elephant fish (SRP013772), and nurse shark (SRP018197), the *ab initio* prediction of the whale shark genome was performed using AUGUSTUS 3.1 with ‘--species=zebrafish’ option^8^. We filtered out genes which contained <30 amino acids. Gene symbols were assigned by best hit to the SwissProt or Trembl databases^12^ using BLASTP^13^ with e-value cutoff 10^−5^. A total of 25,409 out of 34,708 genes were assigned. Finally, we removed possible retro-transposable single exon genes. The resulting gene model contained 28,483 protein coding genes (Table S8).
2) For the EVM approach, we performed homology-based gene prediction with additional species (Table S9) and combined the prediction results with the *ab initio* prediction results [AUGUTUS^8^, MAKER^14^] using EVM^9^ (the weights of intermediate gene models for EVM^9^ integration is noted in Table S10). We predicted 25,915 protein coding genes using the EVM^9^ approach (Table S11).

**Table S8.** Statistics of the AUGUSTUS predicted gene set

| Categories | Number or length (bp) |
| --- | --- |
| Number of genes | 28,483 |
| Average transcript length | 39,530.27 |
| Average number of CDSs per gene | 7.45 |
| Average CDS length per gene | 1,173.7 |
| Average CDS length per exon | 157.54 |

**Table S9.** List of species used in EVM homology-based gene prediction

| Common name | Scientific name | Number of protein sequences | Data Source | Assembly ID |
| --- | --- | --- | --- | --- |
| Human | *Homo sapiens* | 20,129 | Ensembl 86 | GRCh38.p7 |
| Mouse | *Mus musculus* | 22,294 | Ensembl 86 | GRCm38.p4 |
| Anole lizard | *Anolis carolinensis* | 18,520 | Ensembl 86 | AnoCar2.0 |
| Xenopus | *Xenopus tropicalis* | 18,000 | Ensembl 86 | JGI 4.2 |
| Coelacanth | *Latimeria chalumnae* | 19,198 | Ensembl 86 | LatCha1 |
| Guppy | *Poecilia reticulata* | 17,907 | NCBI Refseq | GCF_000633615.1 |
| Turquoise killifish | *Nothobranchius furzeri* | 21,100 | NCBI Refseq | GCF_001465895.1 |
| Medaka | *Oryzias latipes* | 18,937 | Ensembl 86 | HdrR |
| Tilapia | *Oreochromis niloticus* | 20,467 | Ensembl 86 | Orenil1.0 |
| Northern pike | *Esox lucius* | 21,396 | NCBI Refseq | GCF_000721915.3 |
| Zebrafish | *Danio rerio* | 24,309 | Ensembl 86 | GRCz10 |
| Spotted gar | *Lepisosteus oculatus* | 17,874 | Ensembl 86 | LepOcu1 |
| Channel catfish | *Ictalurus punctatus* | 22,463 | NCBI Refseq | GCF_001660625.1 |
| Elephant fish | *Callorhynchus milii* | 15,669 | NCBI Refseq | GCF_000165045.1 |
| Lamprey | *Petromyzon marinus* | 10,048 | Ensembl 86 | Pmarinus_7.0 |

**Table S10.** EVM weights for each gene model

| **Approach** | **Program** | **EVM weight** |
| --- | --- | --- |
| Homology based | exonerate:AnoleLizard | 3 |
| Homology based | exonerate:ChannelCatfish | 3 |
| Homology based | exonerate:Coelacanth | 3 |
| Homology based | exonerate:ElephantFish | 3 |
| Homology based | exonerate:Guppy | 3 |
| Homology based | exonerate:Human | 5 |
| Homology based | exonerate:Lamprey | 3 |
| Homology based | exonerate:Medaka | 3 |
| Homology based | exonerate:Mouse | 3 |
| Homology based | exonerate:NorthernPike | 3 |
| Homology based | exonerate:SpottedGar | 3 |
| Homology based | exonerate:Tilapia | 3 |
| Homology based | exonerate:TurquoiseKillifish | 3 |
| Homology based | exonerate:Xenopus | 3 |
| Homology based | exonerate:Zebrafish | 5 |
| *ab initio* | maker | 5 |
| *ab initio* | Augustus | 15 |

**Table S11.** Statistics of the EVM predicted gene set

| **Categories** | **Number or length (bp)** |
| --- | --- |
| Number of genes | 25,915 |
| Average transcript length | 37,878.36 |
| Average number of CDSs per gene | 7.29 |
| Average CDS length per gene | 1,179.53 |
| Average CDS length per exon | 161.69 |

### 1.8 Genome assembly quality assessment

Assembly quality was assessed by mapping the paired-end DNA reads and the synthetic long read to the final scaffolds using BWA-MEM^2^. The mapping rate was 99.85% for the short reads (Table S12) and 95.14% for TSLRs (Table S13). The genome assembly and completeness of the gene annotation were also assessed using the Benchmarking Universal Single-Copy Orthologs (BUSCO) approach^15^. The two annotation methods (AUGUSTUS^8^ and EVM^9^) had 88.2% and 84.3% complete BUSCO sets, respectively; which are both higher than the previously published draft genome assembly (Table S14).

**Table S12.** Assembly quality assessment using self-mapping of short reads

| **Library** | **Number of filtered reads** | **Number of mapped reads** | **Percentage of mapped reads** |
| --- | --- | --- | --- |
| 170bp | 1,394,746,573 | 1,394,426,151 | 99.98% |
| 500bp | 940,285,287 | 940,078,838 | 99.98% |
| 700bp | 801,359,278 | 801,149,022 | 99.97% |
| 2kb | 236,768,852 | 235,357,729 | 99.40% |
| 5kb | 136,682,871 | 134,810,545 | 98.63% |
| 10kb | 94,350,139 | 93,601,841 | 99.21% |
| 15kb | 85,609,884 | 84,778,286 | 99.03% |
| Total | 3,689,802,884 | 3,684,202,412 | 99.85% |

**Table S13.** Assembly quality assessment using self-mapping of TSLRs

| **Read length**<br>**(bp)** | **Number of reads** | **Number of $\geq 90\%$ covered**<br>**reads** |  | **Number of $\geq 50\%$**<br>**covered reads** |  |
| --- | --- | --- | --- | --- | --- |
| 0-999 | 178,106 | 142,280 | 79.89% | 161,219 | 90.52% |
| 1,000-1,999 | 142,565 | 122,562 | 85.97% | 135,653 | 95.15% |
| 2,000-2,999 | 88,352 | 78,300 | 88.62% | 85,482 | 96.75% |
| 3,000-3,999 | 66,397 | 58,688 | 88.39% | 64,159 | 96.63% |
| 4,000-4,999 | 51,371 | 45,225 | 88.04% | 49,726 | 96.80% |
| 5,000-5,999 | 42,530 | 37,009 | 87.02% | 41,120 | 96.68% |
| 6,000-6,999 | 38,767 | 33,442 | 86.26% | 37,472 | 96.66% |
| 7,000-7,999 | 36,124 | 31,037 | 85.92% | 34,900 | 96.61% |
| 8,000-8,999 | 39,523 | 33,747 | 85.39% | 38,211 | 96.68% |
| 9,000-9,999 | 82,517 | 70,444 | 85.37% | 79,902 | 96.83% |
| 10,000-10,999 | 56,755 | 48,176 | 84.88% | 54,852 | 96.65% |
| 11,000-11,999 | 21,202 | 17,840 | 84.14% | 20,481 | 96.60% |
| 12,000-12,999 | 3,954 | 3,271 | 82.73% | 3,818 | 96.56% |
| 13,000-13,999 | 166 | 118 | 71.08% | 157 | 94.58% |
| 14,000-14,999 | 26 | 9 | 34.62% | 25 | 96.15% |
| >15,000 | 70 | 6 | 8.57% | 53 | 75.71% |
| Total | 848,425 | 722,154 | 85.12% | 807,230 | 95.14% |

**Table S14.** Assessment of the genome assembly and gene completeness using the BUSCO approach, compared to the initial draft whale shark assembly

|  | Whale shark<br>(AUGUSTUS) | Whale shark<br>(EVM) | Whale shark<br>(GCA_001642345.2) |
| --- | --- | --- | --- |
| Complete (Gene) | 2,279 (88.2%) | 2,180 (84.3%) | 1,934 (74.7%) |
| Duplicated (Gene) | 51 (2.0%) | 88 (3.4%) | 84 (3.2%) |
| Fragmented (Gene) | 136 (5.3%) | 271 (10.5%) | 283 (10.9%) |
| Missing (Gene) | 171 (6.5%) | 135 (5.2%) | 369 (14.4%) |
| Number of<br>single-copy orthologous genes | 2,586 | 2,586 | 2,586 |

## 2. Comparative genomic studies

### 2.1 Data resources

Genome sequences and gene sets for 69 species were downloaded from Ensembl FTP (ftp://ftp.ensembl.org/pub/release-86/). An additional twelve species were added from NCBI FTP (ftp://ftp.ncbi.nlm.nih.gov/genomes/) (Table S15).

**Table S15.** List of 82 species and their data sources

| Common name | Species name | Class | Data Source | Assembly ID |
| --- | --- | --- | --- | --- |
| Gorilla | *Gorilla gorilla* | Mammalia | Ensembl 86 | gorGor3.1 |
| Chimpanzee | *Pan troglodytes* | Mammalia | Ensembl 86 | CHIMP2.1.4 |
| Human | *Homo sapiens* | Mammalia | Ensembl 86 | GRCh38.p7 |
| Orangutan | *Pongo abelii* | Mammalia | Ensembl 86 | PPYG2 |
| Gibbon | *Nomascus leucogenys* | Mammalia | Ensembl 86 | Nleu1.0 |
| Macaque | *Macaca mulatta* | Mammalia | Ensembl 86 | Mmul_8.0.1 |
| Olive baboon | *Papio anubis* | Mammalia | Ensembl 86 | PapAnu2.0 |
| Green monkey | *Chlorocebus sabaues* | Mammalia | Ensembl 86 | ChlSab1.1 |
| Marmoset | *Callithrix jacchus* | Mammalia | Ensembl 86 | C_jacchus3.2.1 |
| Tarsier | *Tarsius syrichta* | Mammalia | Ensembl 86 | tarSyr1 |
| Mouse Lemur | *Microcebus murinus* | Mammalia | Ensembl 86 | Mmur_2.0 |
| Galago | *Otolemur garnettii* | Mammalia | Ensembl 86 | OtoGar3 |
| Mouse | *Mus musculus* | Mammalia | Ensembl 86 | GRCm38.p4 |
| Rat | *Rattus norvegicus* | Mammalia | Ensembl 86 | Rnor_6.0 |
| Kangaroo rat | *Dipodomys ordii* | Mammalia | Ensembl 86 | dipOrd1 |
| Guinea pig | *Cavia porcellus* | Mammalia | Ensembl 86 | cavPor3 |
| Naked mole rat | *Heterocephalus glaber* | Mammalia | NCBI | HetGla_female_1.0 |
| Squirrel | *Ictidomys tridecemlineatus* | Mammalia | Ensembl 86 | spetri2 |
| Pika | *Ochotona princeps* | Mammalia | Ensembl 86 | OchPri2.0 |
| Rabbit | *Oryctolagus cuniculus* | Mammalia | Ensembl 86 | OryCun2.0 |
| Tree shrew | *Tupaia belangeri* | Mammalia | Ensembl 86 | tupBel1 |
| Killer whale | *Orcinus orca* | Mammalia | NCBI | GCF_000331955.2 |
| Dolphin | *Tursiops truncatus* | Mammalia | Ensembl 86 | turTru1 |
| Sperm whale | *Physeter catodon* | Mammalia | NCBI | GCF_000472045.1 |
| Minke whale | *Balaenoptera acutorostrata scammoni* | Mammalia | NCBI | GCF_000493695.1 |
| Cattle | *Bos taurus* | Mammalia | Ensembl 86 | UMD3.1 |
| Sheep | *Ovis aries* | Mammalia | Ensembl 86 | Oar_v3.1 |
| Pig | *Sus scrofa* | Mammalia | Ensembl 86 | Sscrofa10.2 |
| Alpaca | *Vicugna pacos* | Mammalia | Ensembl 86 | vicPac1 |
| Dog | *Canis familiaris* | Mammalia | Ensembl 86 | CanFam3.1 |
| Panda | *Ailuropoda melanoleuca* | Mammalia | Ensembl 86 | ailMe11 |
| Ferret | *Mustela putorius furo* | Mammalia | Ensembl 86 | MusPutFur1.0 |
| Cat | *Felis catus* | Mammalia | Ensembl 86 | Felis_catus_6.2 |
| Hedgehog | *Erinaceus europaeus* | Mammalia | Ensembl 86 | eriEur1 |
| Shrew | *Sorex araneus* | Mammalia | Ensembl 86 | sorAra1 |
| Microbat | *Myotis lucifugus* | Mammalia | Ensembl 86 | Myoluc2.0 |
| Megabat | *Pteropus vampyrus* | Mammalia | Ensembl 86 | pteVam1 |
| Horse | *Equus caballus* | Mammalia | Ensembl 86 | Equ Cab 2 |
| Lesser hedgehog tenrec | *Echinops telfairi* | Mammalia | Ensembl 86 | TENREC |
| Elephant | *Loxodonta africana* | Mammalia | Ensembl 86 | Loxafr3.0 |
| Hyrax | *Procavia capensis* | Mammalia | Ensembl 86 | proCap1 |
| Sloth | *Choloepus hoffmanni* | Mammalia | Ensembl 86 | choHof1 |
| Armadillo | *Dasypus novemcinctus* | Mammalia | Ensembl 86 | Dasnov3.0 |
| Tasmanian devil | *Sarcophilus harrisii* | Mammalia | Ensembl 86 | Devil_ref v7.0 |
| Wallaby | *Macropus eugenii* | Mammalia | Ensembl 86 | Meug_1.0 |
| Opossum | *Monodelphis domestica* | Mammalia | Ensembl 86 | monDom5 |
| Platypus | *Ornithorhynchus anatinus* | Mammalia | Ensembl 86 | OANA5 |
| Chicken | *Gallus gallus* | Aves | Ensembl 86 | Gallus_gallus-5.0 |
| Turkey | *Meleagris gallopavo* | Aves | Ensembl 86 | Turkey_2.01 |
| Duck | *Anas platyrhynchos* | Aves | Ensembl 86 | BGI_duck_1.0 |
| Zebra Finch | *Taeniopygia guttata* | Aves | Ensembl 86 | taeGut3.2.4 |
| Flycatcher | *Ficedula albicollis* | Aves | Ensembl 86 | FicAlb_1.4 |
| Chinese softshell turtle | *Pelodiscus sinensis* | Reptilia | Ensembl 86 | PelSin_1.0 |
| Anole lizard | *Anolis carolinensis* | Reptilia | Ensembl 86 | AnoCar2.0 |
| Xenopus | *Xenopus tropicalis* | Amphibia | Ensembl 86 | JGI 4.2 |
| Coelacanth | *Latimeria chalumnae* | Sarcopterygii | Ensembl 86 | LatCha1 |
| Guppy | *Poecilia reticulata* | Actinopterygii | NCBI | GCF_000633615.1 |
| Amazon molly | *Poecilia formosa* | Actinopterygii | Ensembl 86 | Poecilia_formosa-5.1.2 |
| Platyfish | *Xiphophorus maculatus* | Actinopterygii | Ensembl 86 | Xipmac4.4.2 |
| Turquoise killifish | *Nothobranchius furzeri* | Actinopterygii | NCBI | GCF_001465895.1 |
| Medaka | *Oryzias latipes* | Actinopterygii | Ensembl 86 | HdrR |
| Tilapia | *Oreochromis niloticus* | Actinopterygii | Ensembl 86 | Orenil1.0 |
| Fugu | *Takifugu rubripes* | Actinopterygii | Ensembl 86 | FUGU 4.0 |
| Tetraodon | *Tetraodon nigroviridis* | Actinopterygii | Ensembl 86 | TETRAODON 8.0 |
| Stickleback | *Gasterosteus aculeatus* | Actinopterygii | Ensembl 86 | BROAD S1 |
| Cod | *Gadus morhua* | Actinopterygii | Ensembl 86 | gadMor1 |
| Coho salmon | *Oncorhynchus kisutch* | Actinopterygii | NCBI | GCF_002021735.1 |
| Atlantic salmon | *Salmo salar* | Actinopterygii | NCBI | GCF_000233375.1 |
| Northern pike | *Esox lucius* | Actinopterygii | NCBI | GCF_000721915.3 |
| Zebrafish | *Danio rerio* | Actinopterygii | Ensembl 86 | GRCz10 |
| Common carp | *Cyprinus carpio* | Actinopterygii | NCBI | GCF_000951615.1 |
| Cave fish | *Astyanax mexicanus* | Actinopterygii | Ensembl 86 | AstMex102 |
| Spotted gar | *Lepisosteus oculatus* | Actinopterygii | Ensembl 86 | LepOcu1 |
| Channel catfish | *Ictalurus punctatus* | Actinopterygii | NCBI | GCF_001660625.1 |
| Elephant fish | *Callorhynchus milii* | Chondrichthyes | NCBI | GCF_000165045.1 |
| Whale shark | *Rhincodon typus* | Chondrichthyes | This study | This study |
| Lamprey | *Petromyzon marinus* | Hyperoartia | Ensembl 86 | Pmarinus_7.0 |
| Vase tunicate | *Ciona intestinalis* | Ascidacea | Ensembl 86 | KH |
| Pacific transparent sea squirt | *Ciona savignyi* | Ascidacea | Ensembl 86 | CSAV 2.0 |
| Nematode | *Caenorhabditis elegans* | Chromadorea | Ensembl 86 | WBcel235 |
| Fruit fly | *Drosophila melanogaster* | Insecta | Ensembl 86 | BDGP6 |
| Yeast | *Saccharomyces cerevisiae* | Saccharomycetes | Ensembl 86 | R64-1-1 |

### 2.2 Comparison of genomic factors

Due to the absence of transcriptome data for fourteen of the comparison species (cod, sloth, hyrax, elephant, lesser tenrec, megabat, shrew, hedgehog, dolphin, tree shrew, pika, kangaroo rat, tarsier, whale shark), we focused our analyses on the genomic features in translated region (Figure 1 and Figure S3-S6).

**Figure S3.**
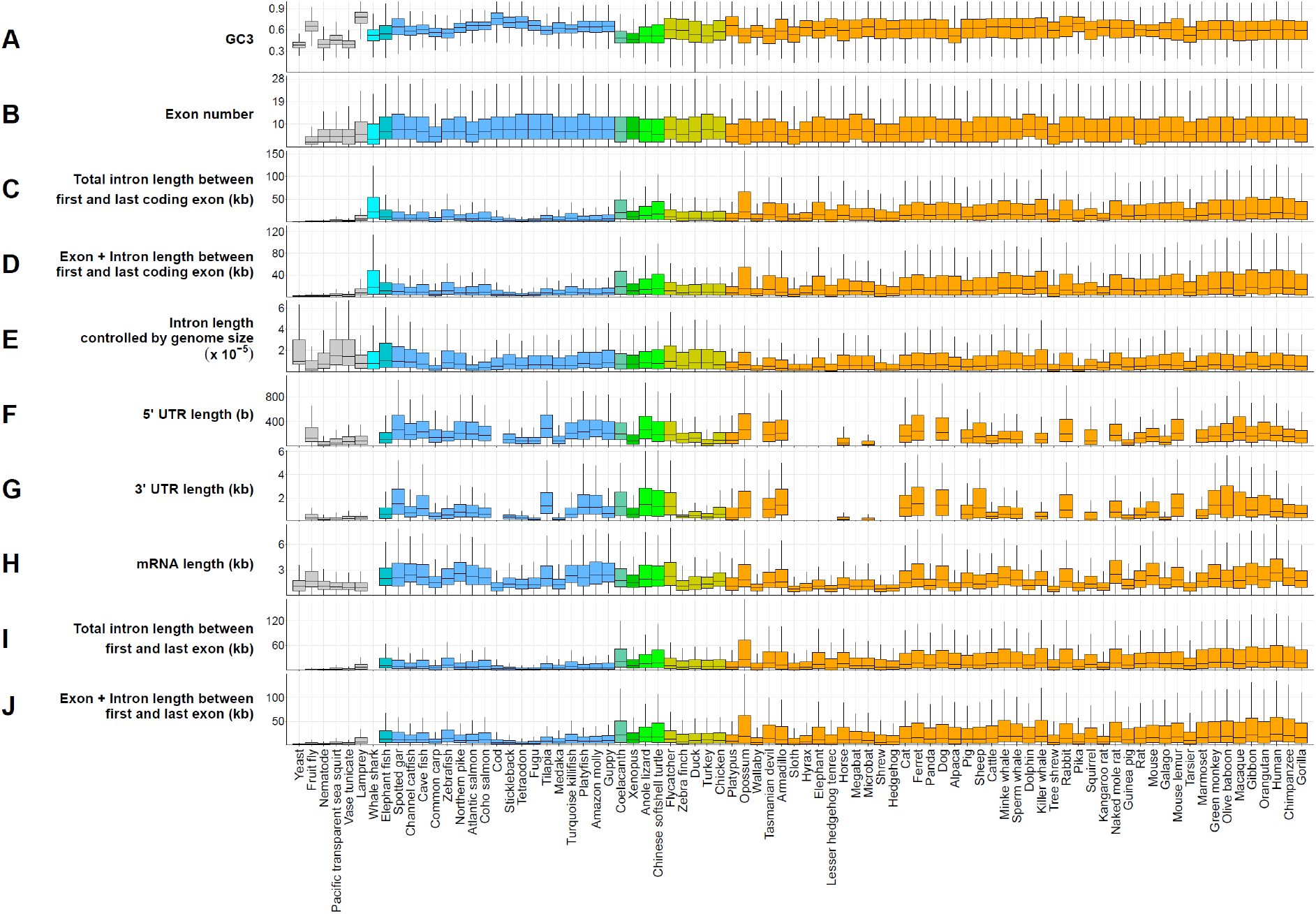
Comparative genomic analysis across 82 species. Extended data from Figure 1. The nine colors of boxes indicate biological classification (gray: Hyperoartia, Ascidiacea, Chromadorea, Insecta and Saccharomycetes, cyan: whale shark, dark turquoise: elephant fish, light blue: Actinopterygii, aquamarine: Sarcopterygii, dark green: Amphibia, light green: Reptilia, dark yellow: Aves, orange: Mammalia).

**Figure S4A.**
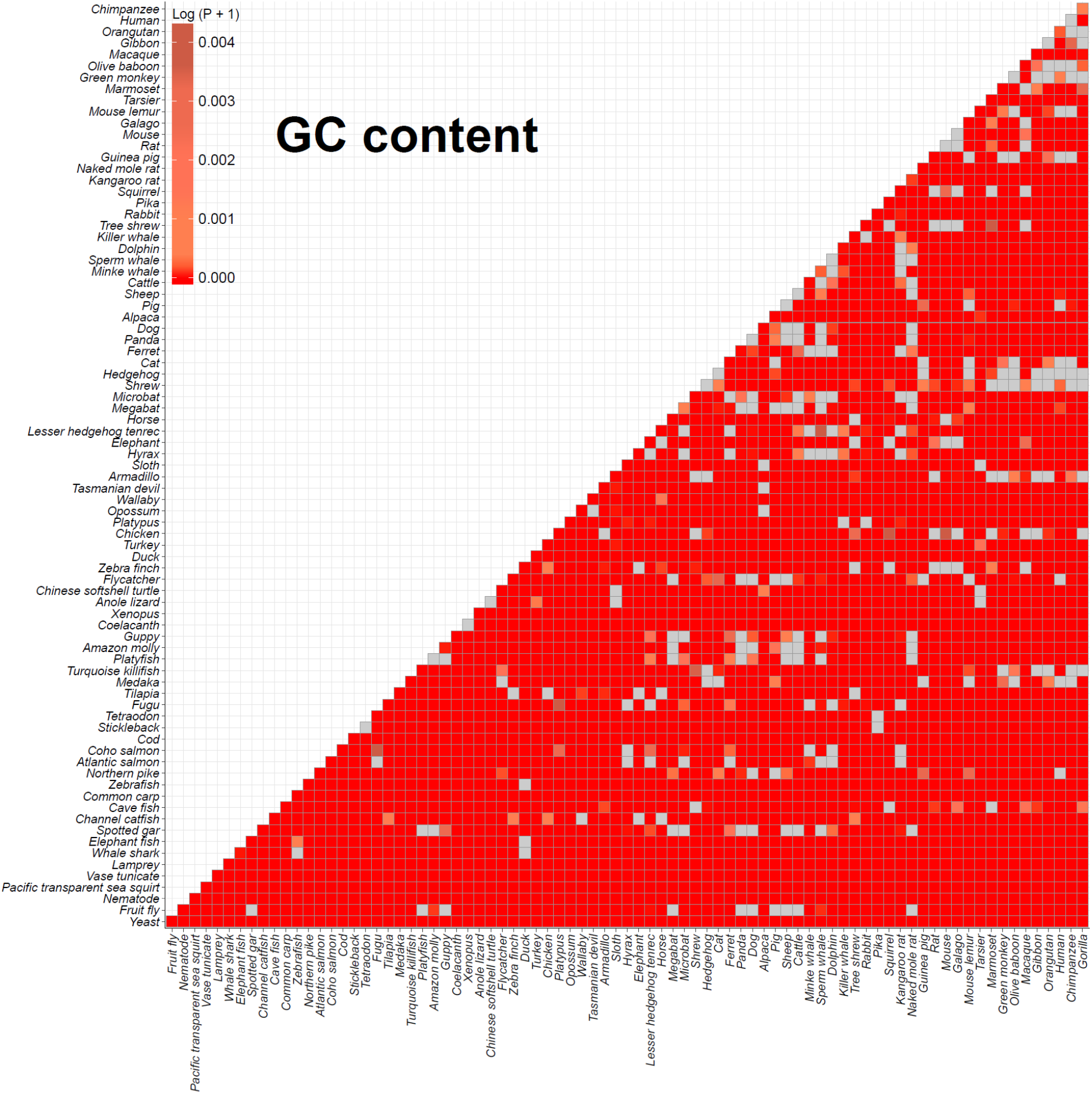
Comparison of GC content in the CDS by Wilcoxon rank sum test among 82 species. Two sided Wilcoxon rank sum tests were computed with the GC content among 82 species. All *p*-values were adjusted using the Benjamini-Hochberg procedure and log-transformed. Deep red indicates significant adjusted *p*-values. Gray boxes indicate *p*-values higher than 0.01.

**Figure S4B.**
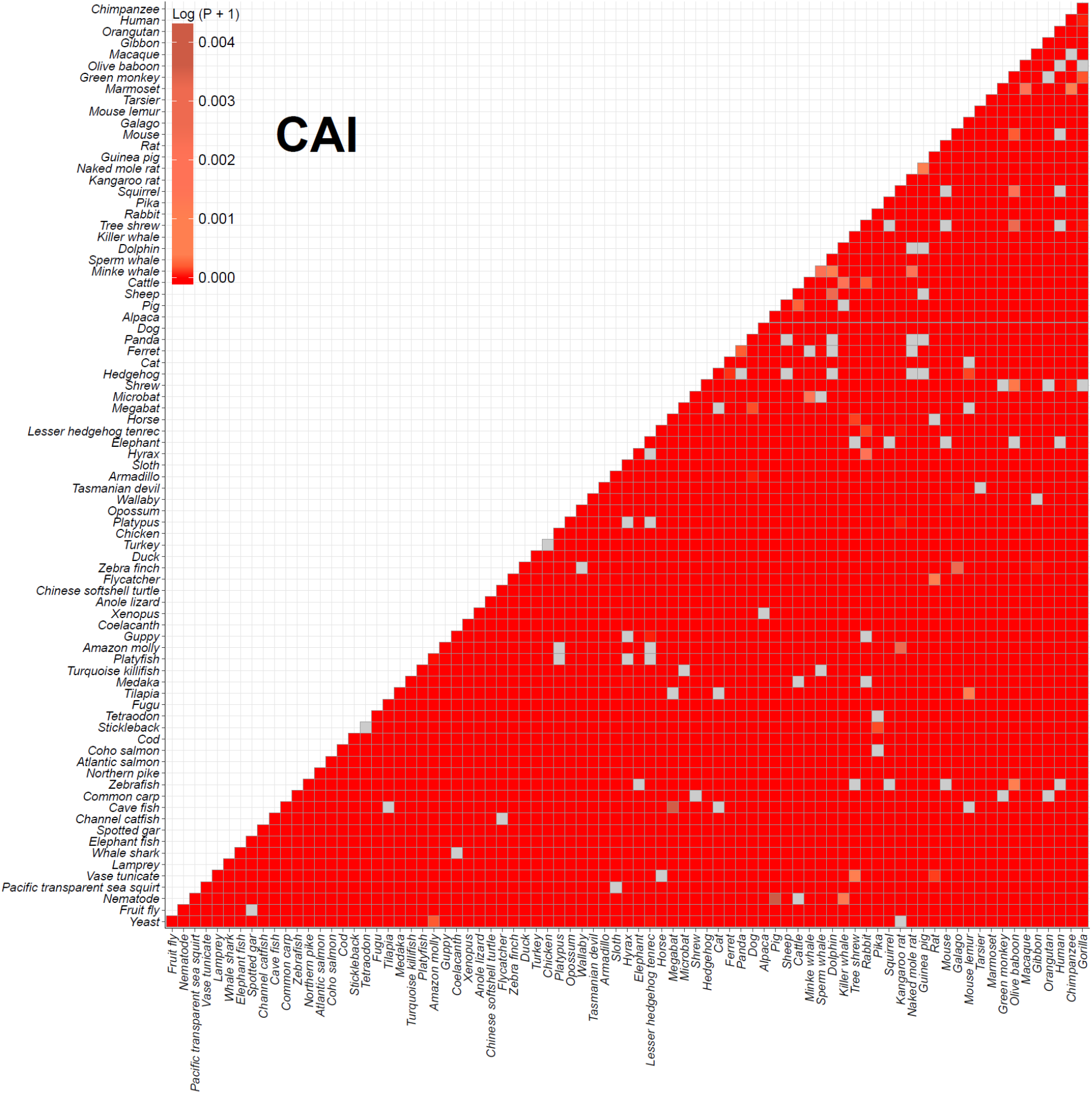
Comparison of CAI by Wilcoxon rank sum test among 82 species. Two sided Wilcoxon rank sum tests were computed with the codon adaptation index (CAI) among 82 species. All *p*-values were adjusted using the Benjamini-Hochberg procedure and log-transformed. Deep red indicates significant adjusted *p*-values. Gray boxes indicate *p*-values higher than 0.01.

**Figure S4C.**
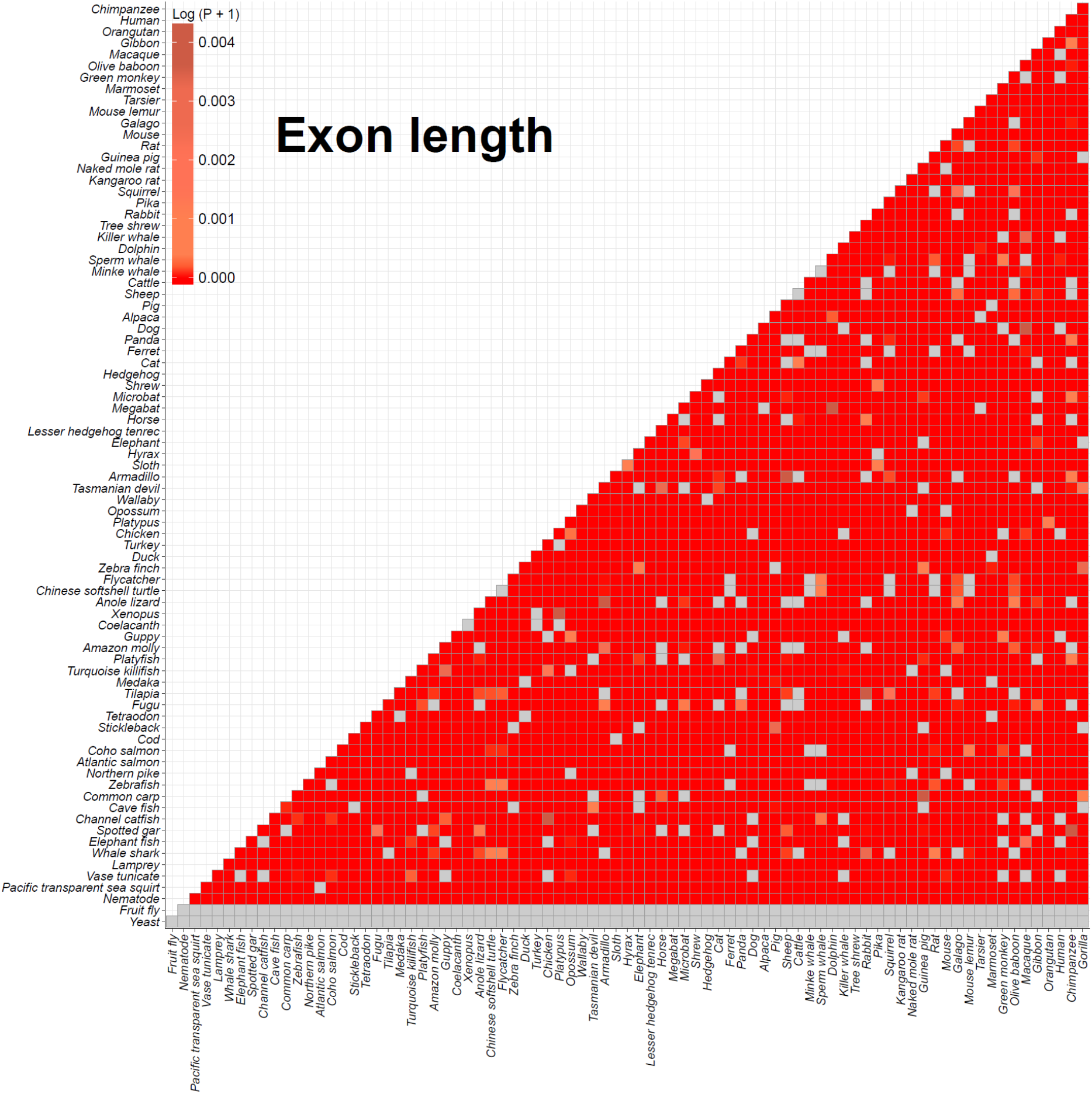
Comparison of exon length by Wilcoxon rank sum test among 82 species. Two sided Wilcoxon rank sum tests were computed with the length of exons between first and last coding exon among 82 species. All *p*-values were adjusted using the Benjamini-Hochberg procedure and log-transformed. Deep red indicates significant adjusted *p*-values. Gray boxes indicate *p*-values higher than 0.01 or NA value (Yeast and Fruit fly).

**Figure S4D.**
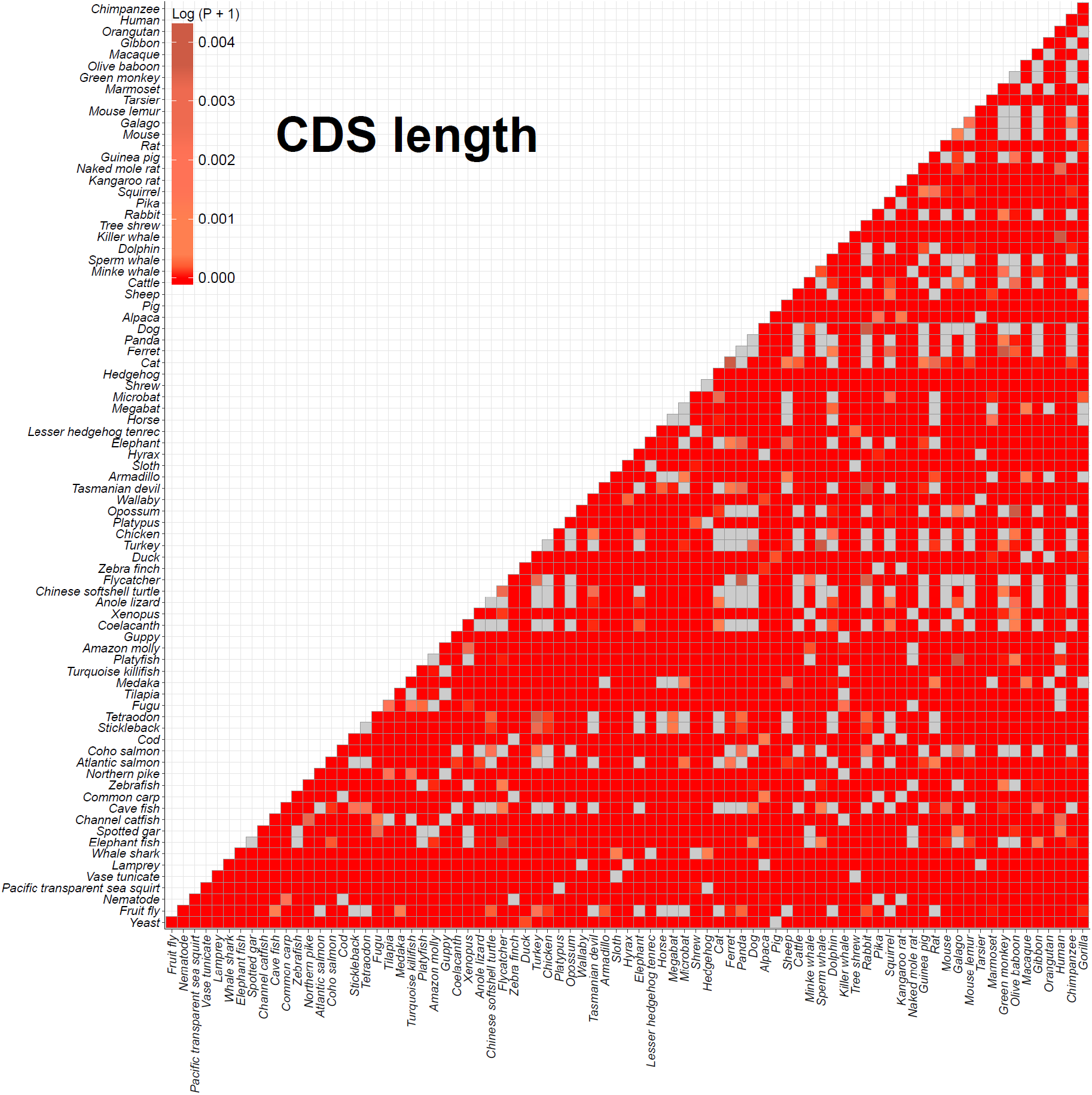
Comparison of CDS length by Wilcoxon rank sum test among 82 species. Two sided Wilcoxon rank sum tests were computed with the CDS length among 82 species. All *p*-values were adjusted using the Benjamini-Hochberg procedure and log-transformed. Deep red indicates significant adjusted *p*-values. Gray boxes indicate *p*-values higher than 0.01.

**Figure S4E.**
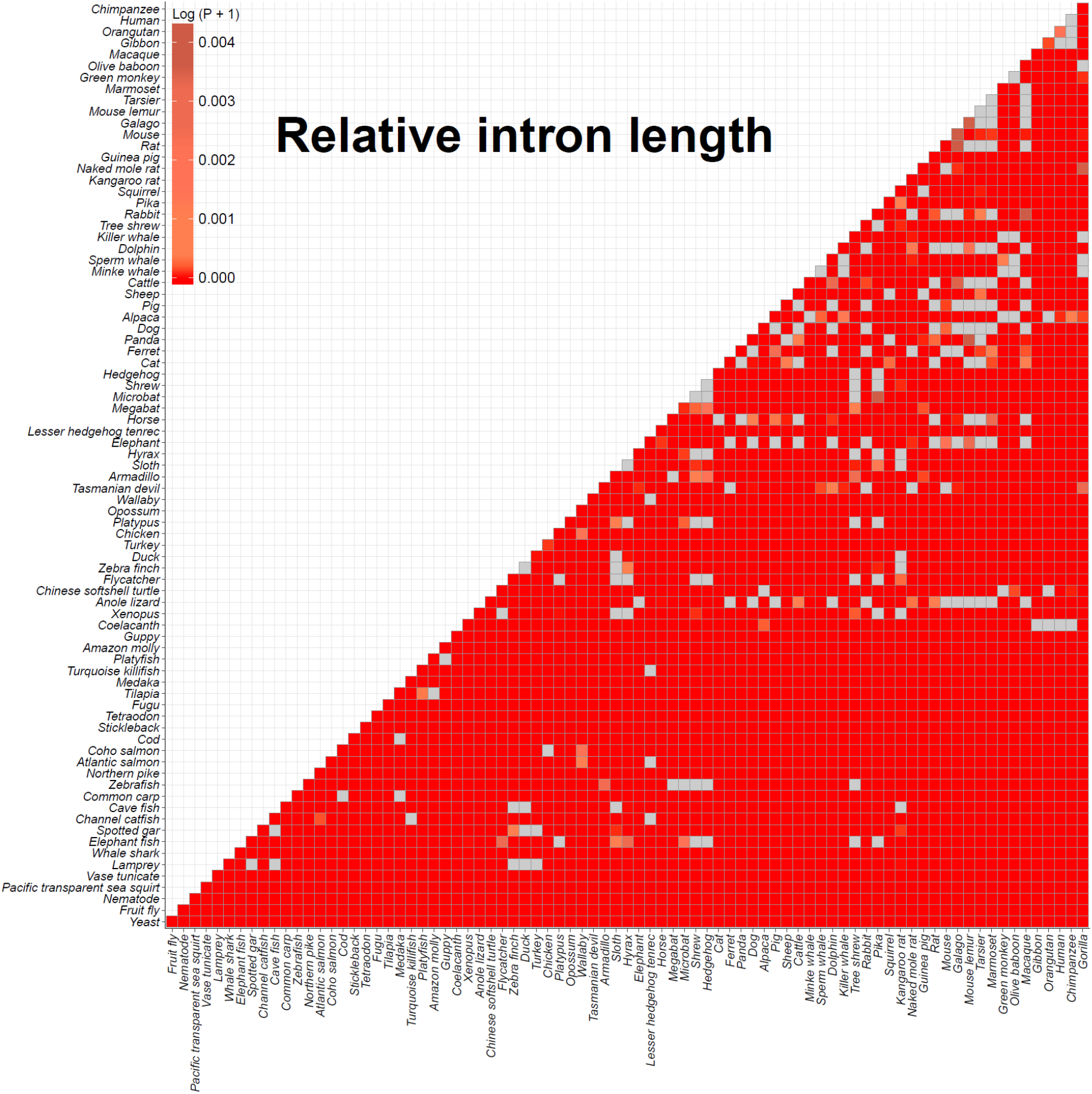
Comparison of relative introns length by Wilcoxon rank sum test among 82 species. Two sided Wilcoxon rank sum tests were computed with the relative intron length among 82 species. The relative intron length was calculated by dividing the total intron length between first and last coding exon by the CDS length. All *p*-values were adjusted using the Benjamini-Hochberg procedure and log-transformed. Deep red indicates significant adjusted *p*-values. Gray boxes indicate *p*-values higher than 0.01.

**Figure S4F.**
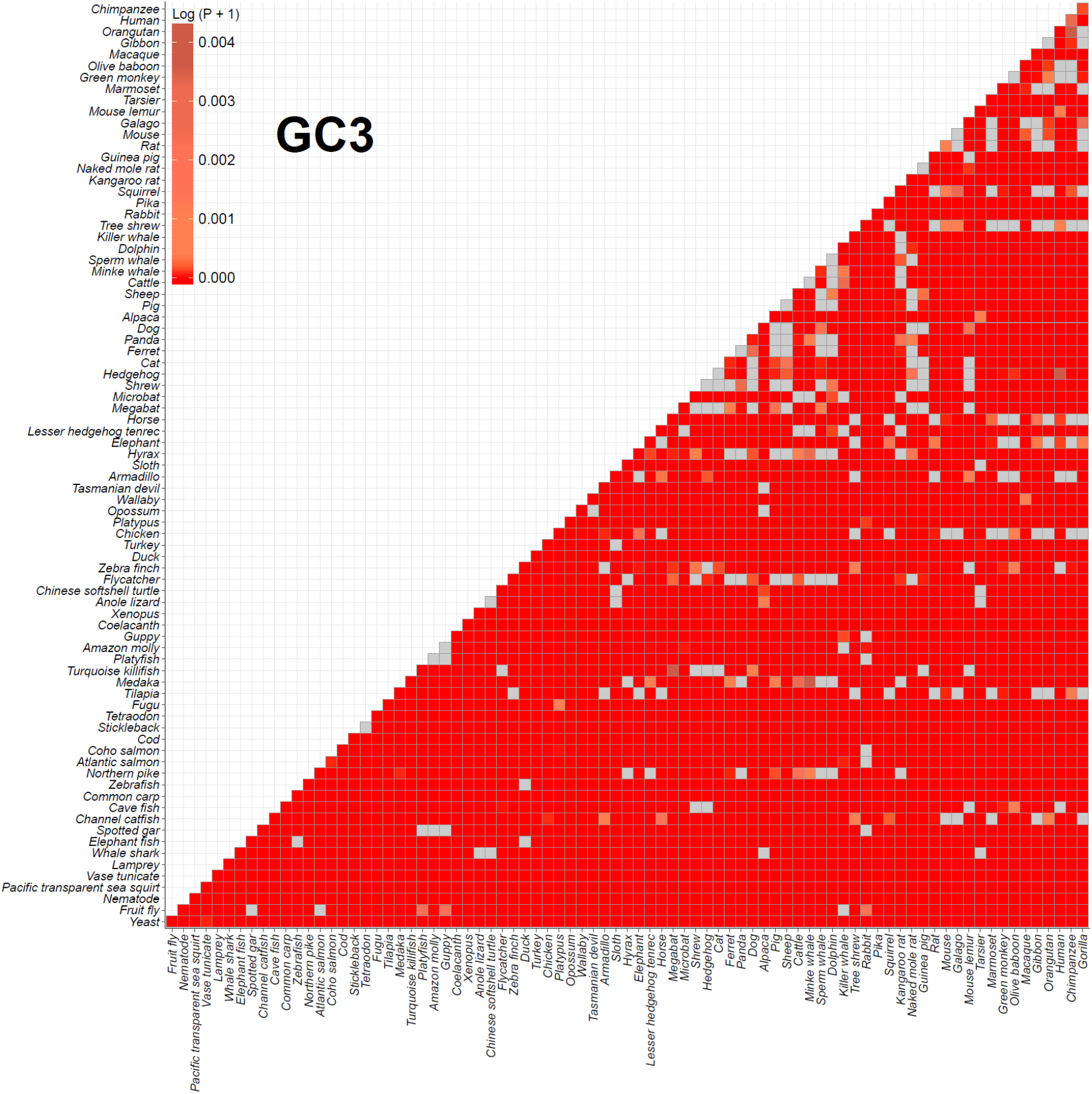
Comparison of GC3 by Wilcoxon rank sum test among 82 species. Two sided Wilcoxon rank sum tests were computed with the GC content at the third codon position (GC3) among 82 species. All *p*-values were adjusted using the Benjamini-Hochberg procedure and log-transformed. Deep red indicates significant adjusted *p*-values. Gray boxes indicate *p*-values higher than 0.01.

**Figure S4G.**
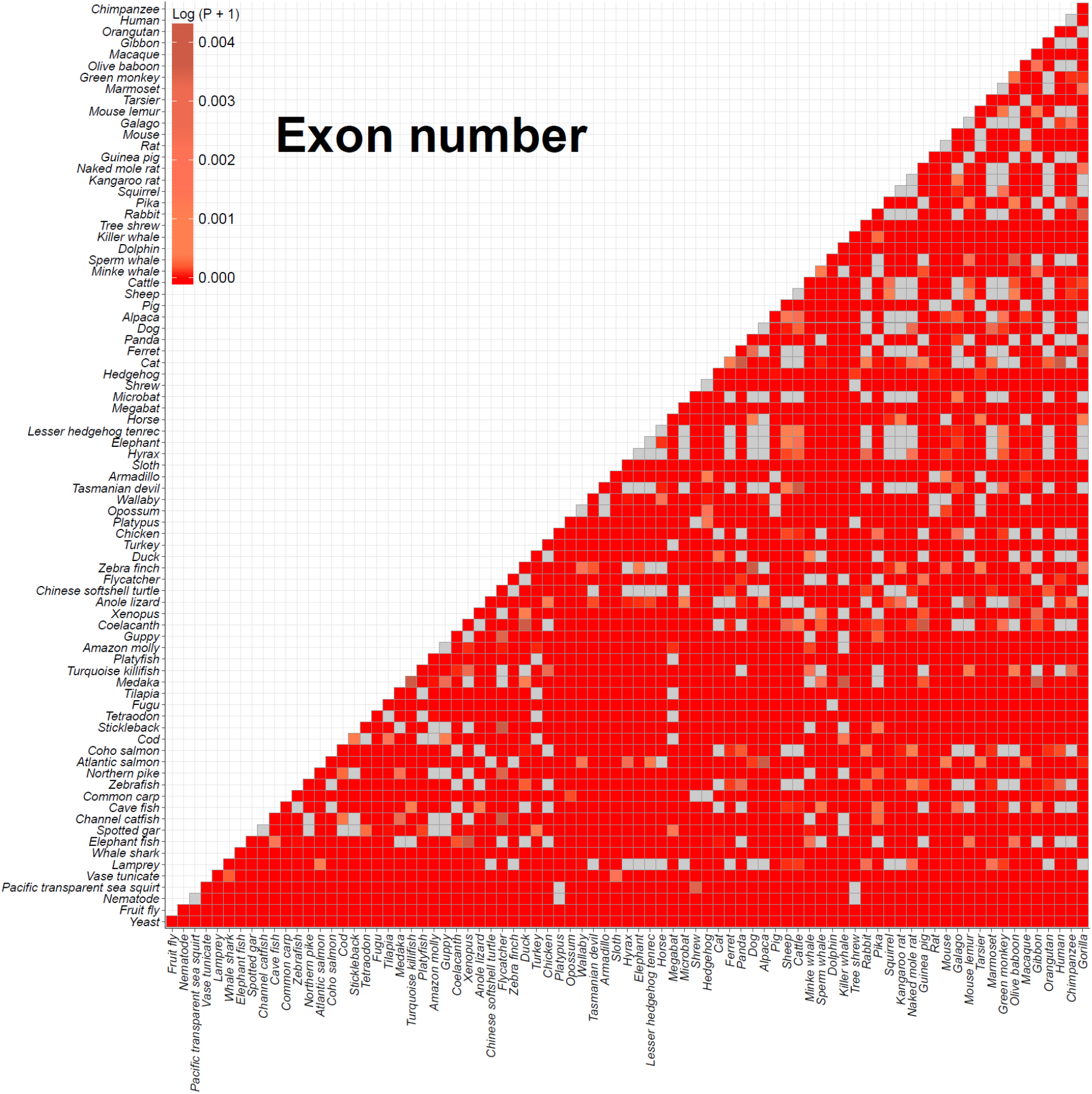
Comparison of exon number by Wilcoxon rank sum test among 82 species. Two sided Wilcoxon rank sum tests were computed with the number of coding exons among 82 species. All *p*-values were adjusted using the Benjamini-Hochberg procedure and log-transformed. Deep red indicates significant adjusted *p*-values. Gray boxes indicate *p*-values higher than 0.01.

**Figure S4H.**
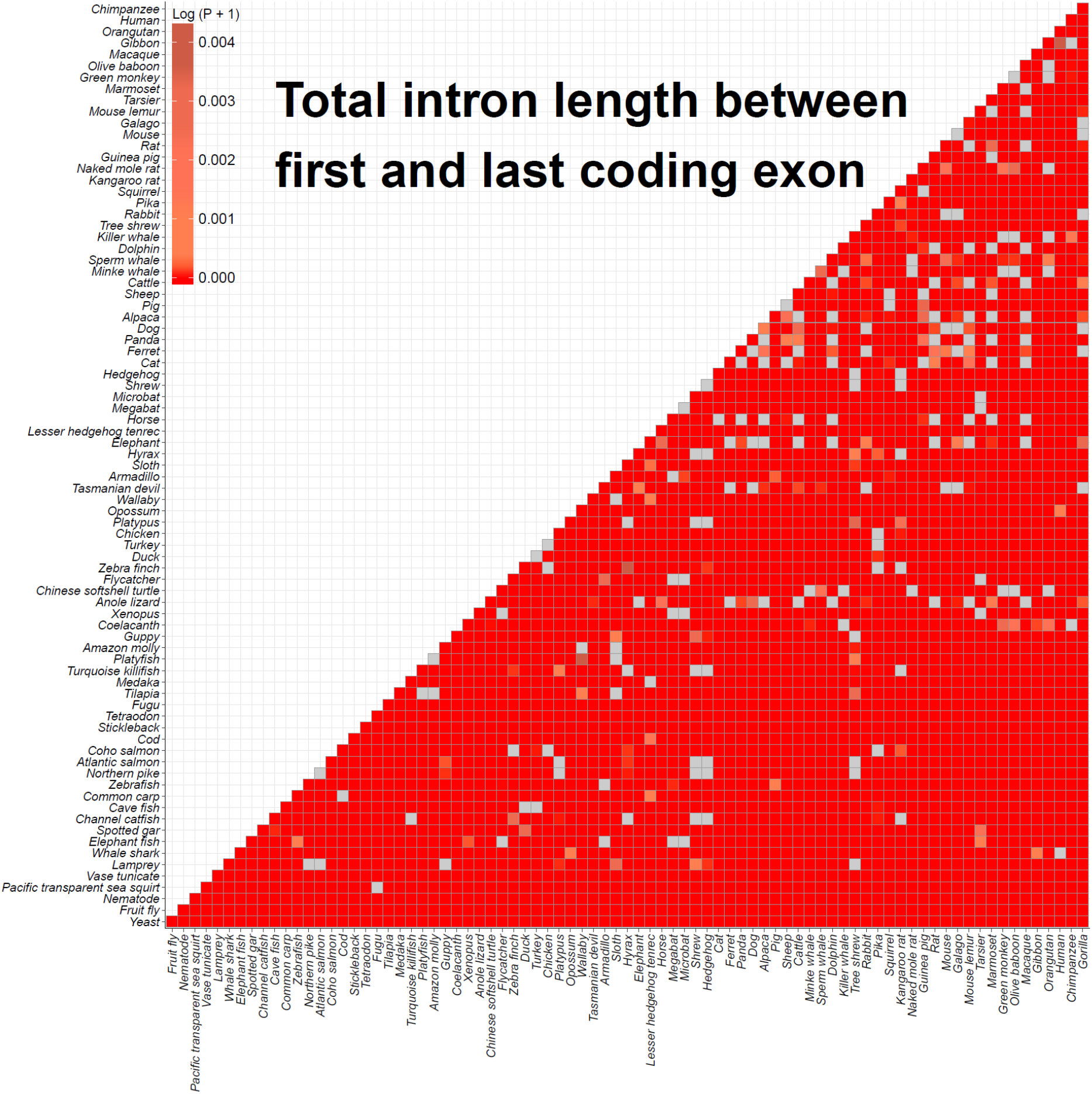
Comparison of total intron length by Wilcoxon rank sum test among 82 species. Two sided Wilcoxon rank sum tests were computed with the total length between first and last exon among 82 species. All *p*-values were adjusted using the Benjamini-Hochberg procedure and log-transformed. Deep red indicates significant adjusted *p*-values. Gray boxes indicate *p*-values higher than 0.01.

**Figure S4I.**
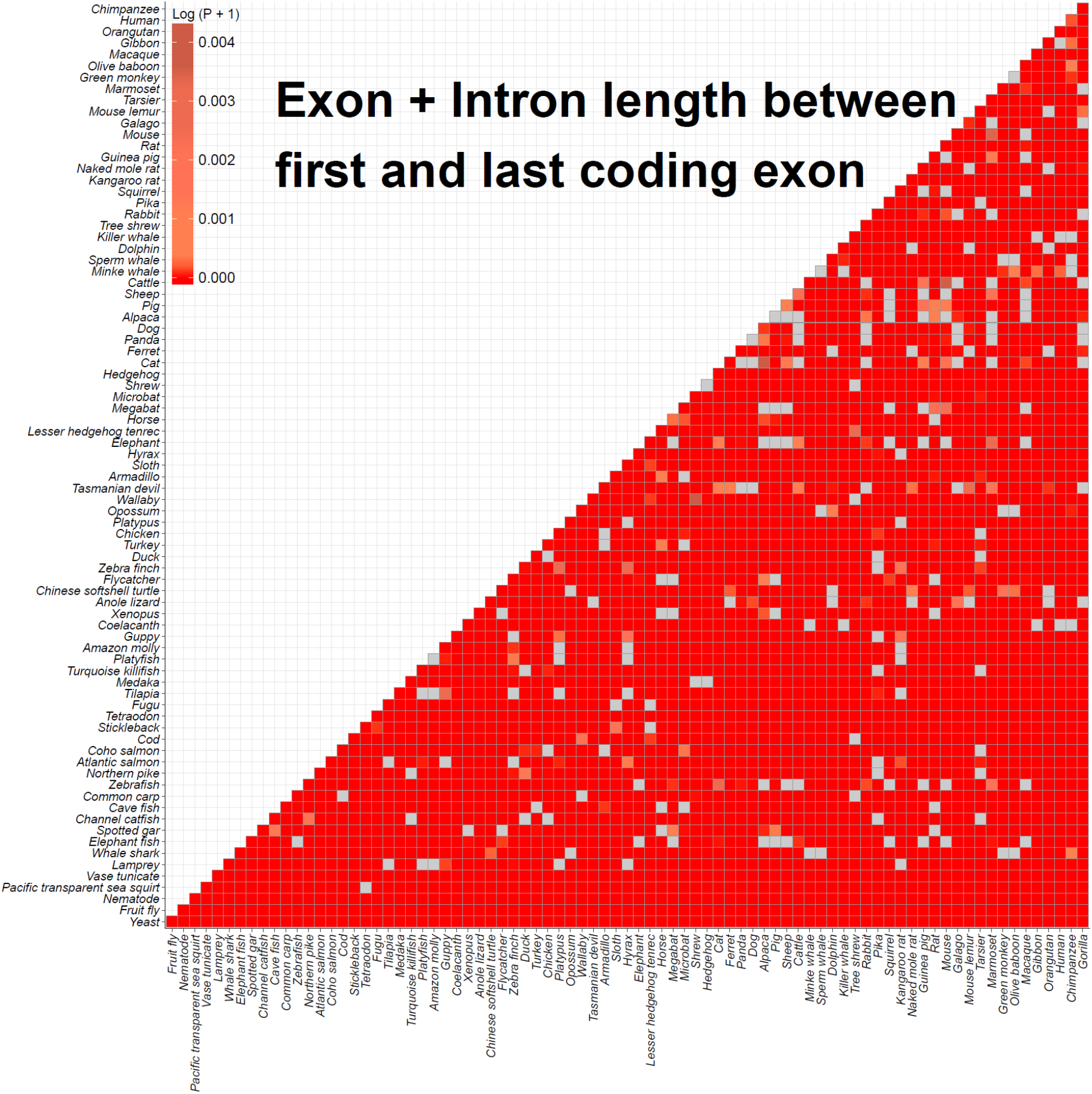
Comparison of sum length of exons and introns by Wilcoxon rank sum test among 82 species. Two sided Wilcoxon rank sum tests were computed with the sum length of exons and introns between first and last coding exon among 82 species. All *p*-values were adjusted using the Benjamini-Hochberg procedure and log-transformed. Deep red indicates significant adjusted *p*-values. Gray boxes indicate *p*-values higher than 0.01.

**Figure S4J.**
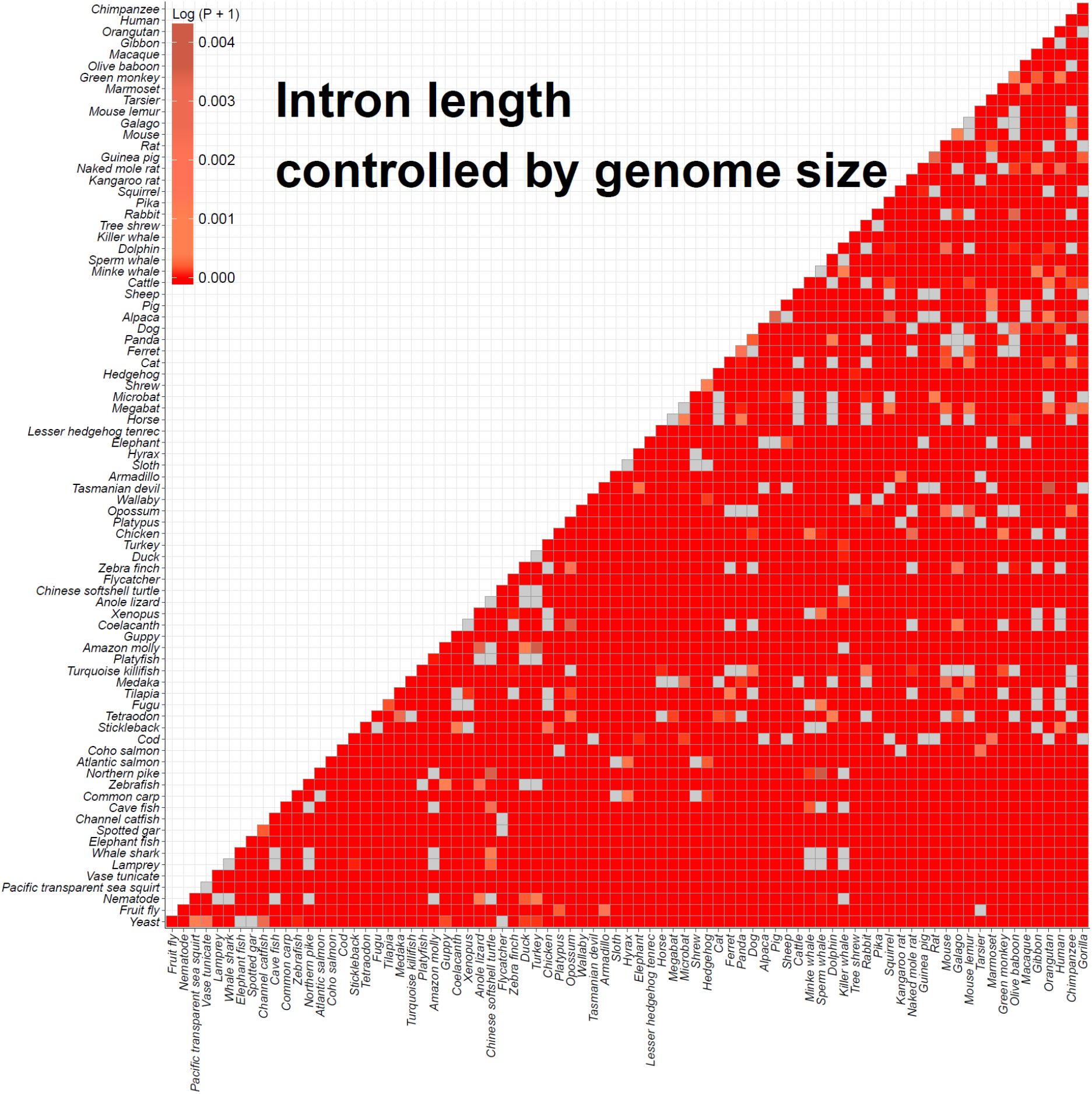
Comparison of controlled introns length by Wilcoxon rank sum test among 82 species. Two sided Wilcoxon rank sum tests were computed with the total intron length between first and last coding exon divided by genome size among 82 species. All *p*-values were adjusted using the Benjamini-Hochberg procedure and log-transformed. Deep red indicates significant adjusted *p*-values. Gray boxes indicate *p*-values higher than 0.01.

**Figure S4K.**
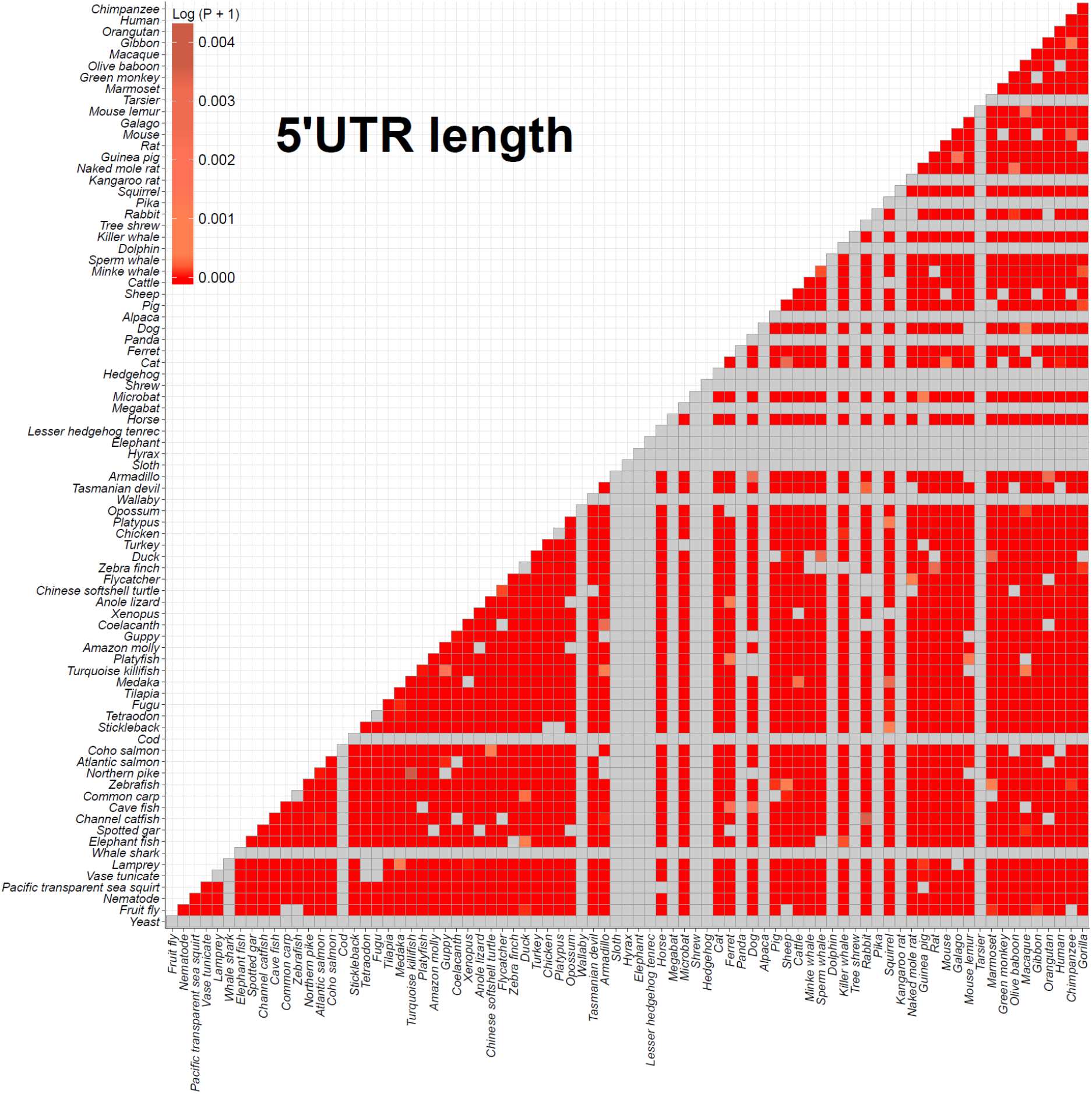
Comparison of 5’ UTR length by Wilcoxon rank sum test among 82 species. Two sided Wilcoxon rank sum tests were computed with the 5’ UTR length among 82 species. All *p*-values were adjusted using the Benjamini-Hochberg procedure and log-transformed. Deep red indicates significant adjusted *p*-values. Gray boxes indicate *p*-values higher than 0.01 or NA values (species which have no 5’ UTR information).

**Figure S4L.**
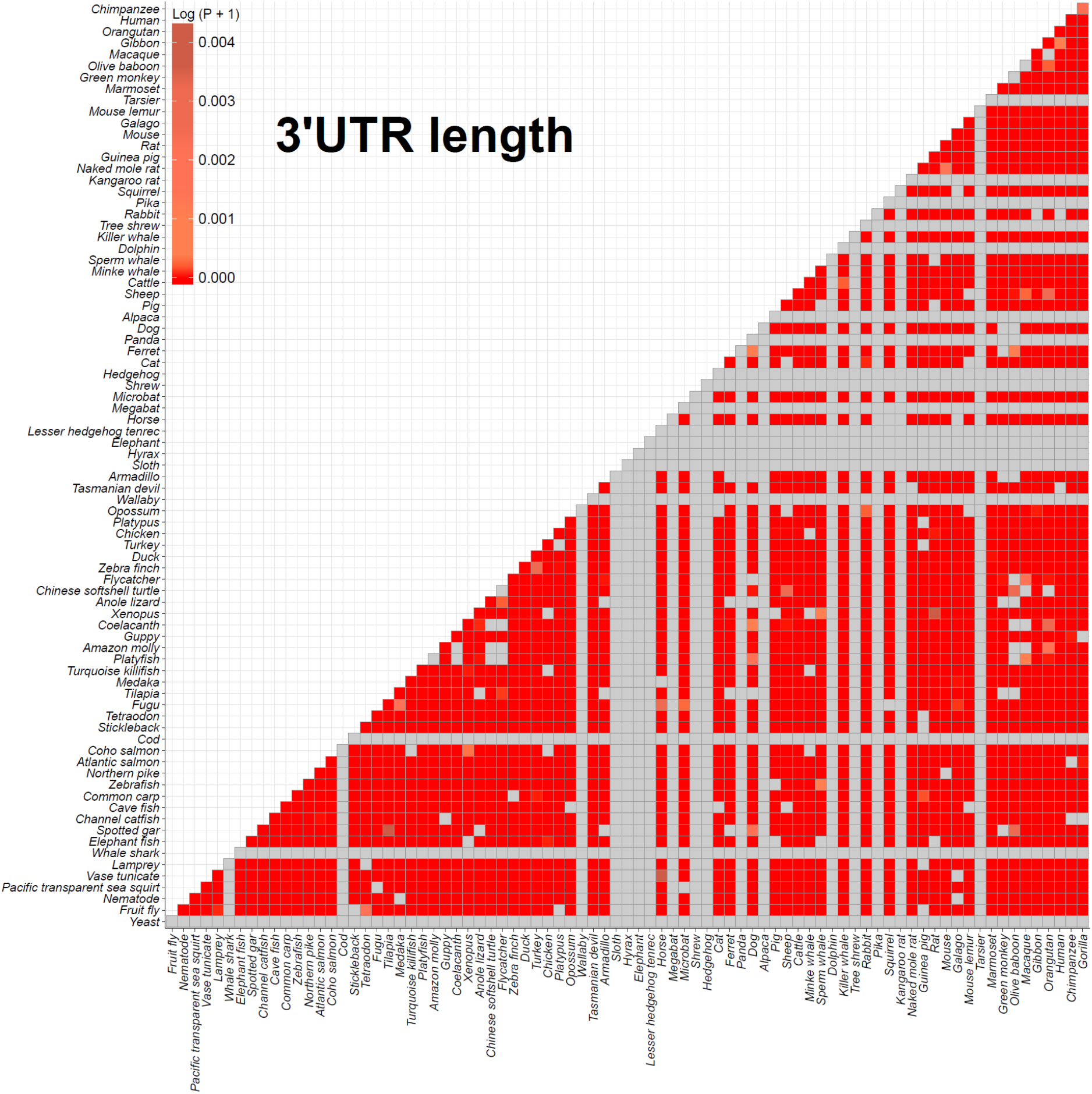
Comparison of 3’ UTR length by Wilcoxon rank sum test among 82 species. Two sided Wilcoxon rank sum tests were computed with the 3’ UTR length among 82 species. All *p*-values were adjusted using the Benjamini-Hochberg procedure and log-transformed. Deep red indicates significant adjusted *p*-values. Gray boxes indicate *p*-values higher than 0.01 or NA values (species which have no 3’ UTR information).

**Figure S4M.**
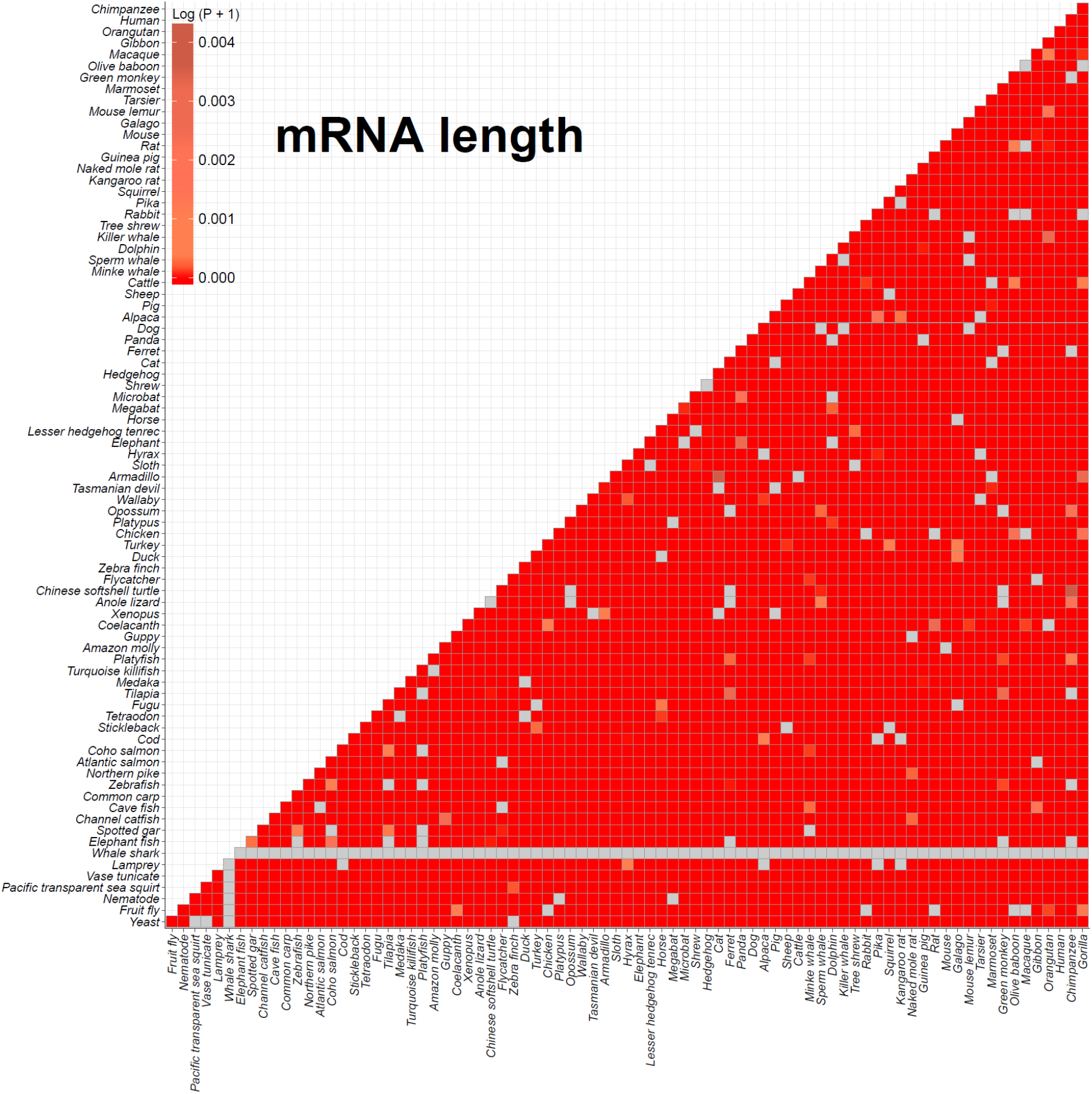
Comparison of mRNA length by Wilcoxon rank sum test among 82 species. Two sided Wilcoxon rank sum tests were computed with the mRNA length among 82 species. All *p*-values were adjusted using the Benjamini-Hochberg procedure and log-transformed. Deep red indicates significant adjusted *p*-values. Gray boxes indicate *p*-values higher than 0.01 or NA values (whale shark which have no mRNA information).

**Figure S4N.**
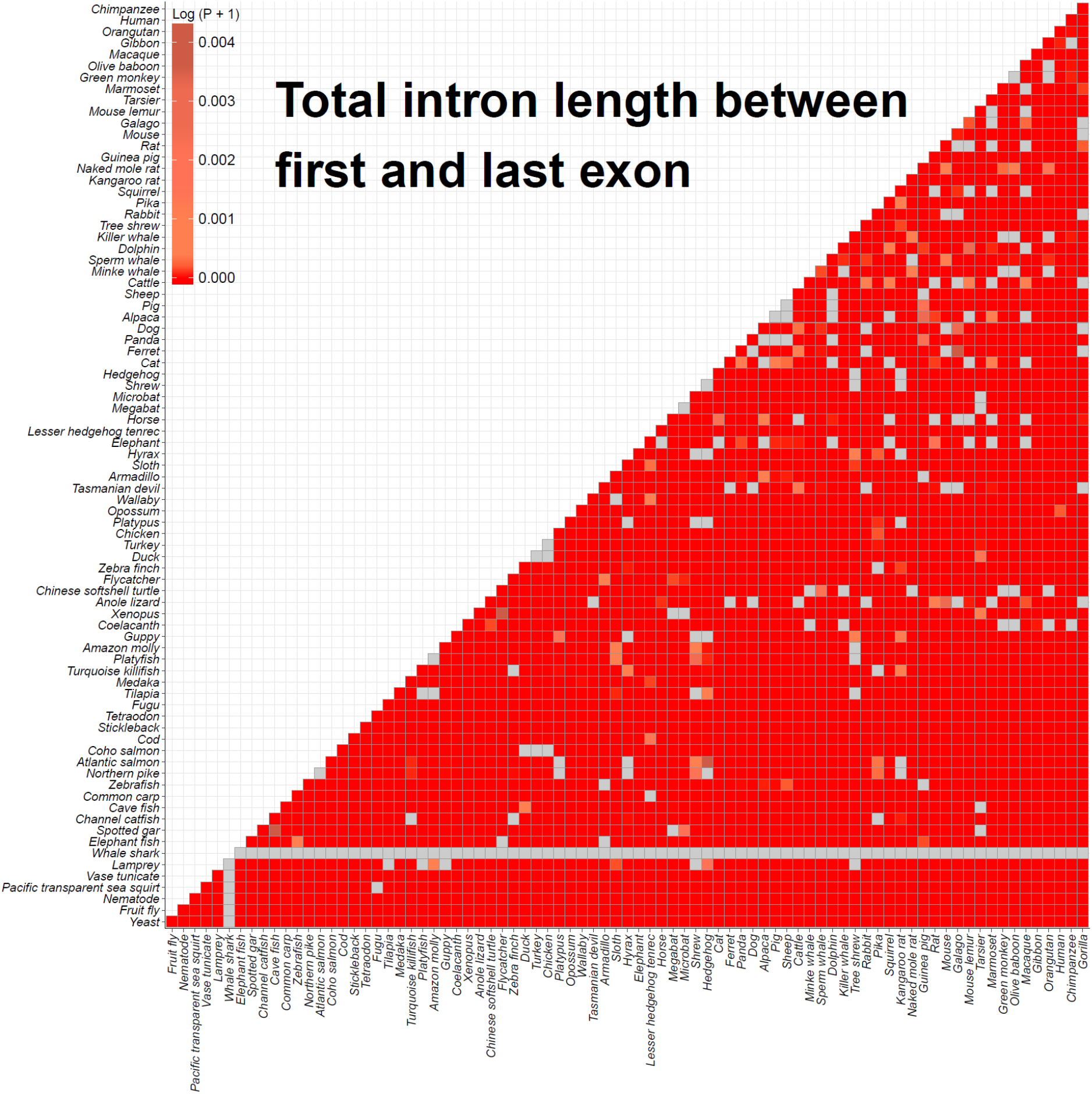
Comparison of total intron length between first and last exon by Wilcoxon rank sum test among 82 species. Two sided Wilcoxon rank sum tests were computed with the total length of introns between first and last exon among 82 species. All *p*-values were adjusted using the Benjamini-Hochberg procedure and log-transformed. Deep red indicates significant adjusted *p*-values. Gray boxes indicate *p*-values higher than 0.01 or NA values (e.g., whale shark, which have no mRNA information).

**Figure S4O.**
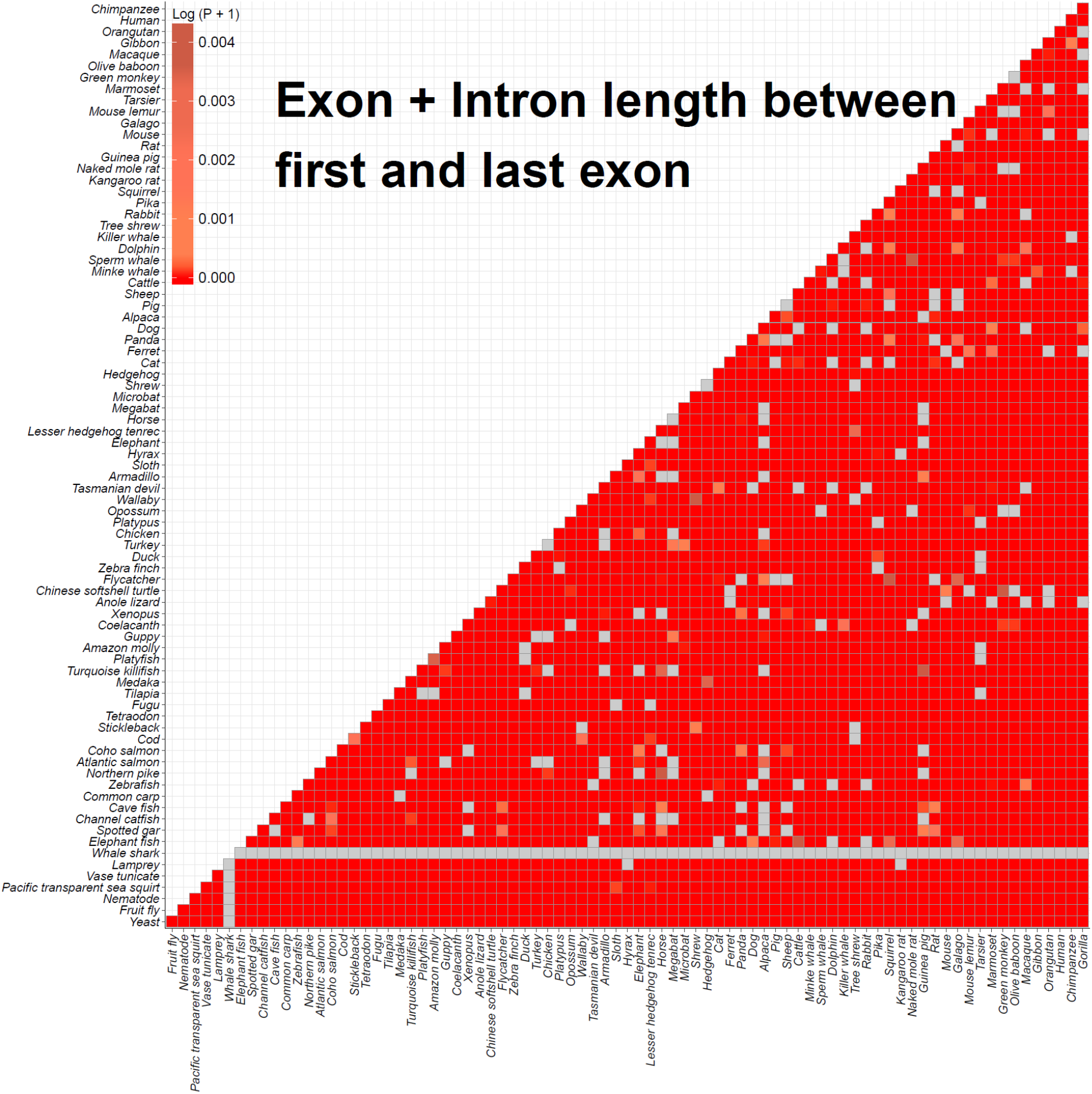
Comparison of sum length of exons and introns between first and last exon by Wilcoxon rank sum test among 82 species. Two sided Wilcoxon rank sum tests were computed with sum length of exons and introns between first and last exon among 82 species. All *p*-values were adjusted using the Benjamini-Hochberg procedure and log-transformed. Deep red indicates significant adjusted *p*-values. Gray boxes indicate *p*-values higher than 0.01 or NA values (whale shark which have no mRNA information).

**Figure S5.**
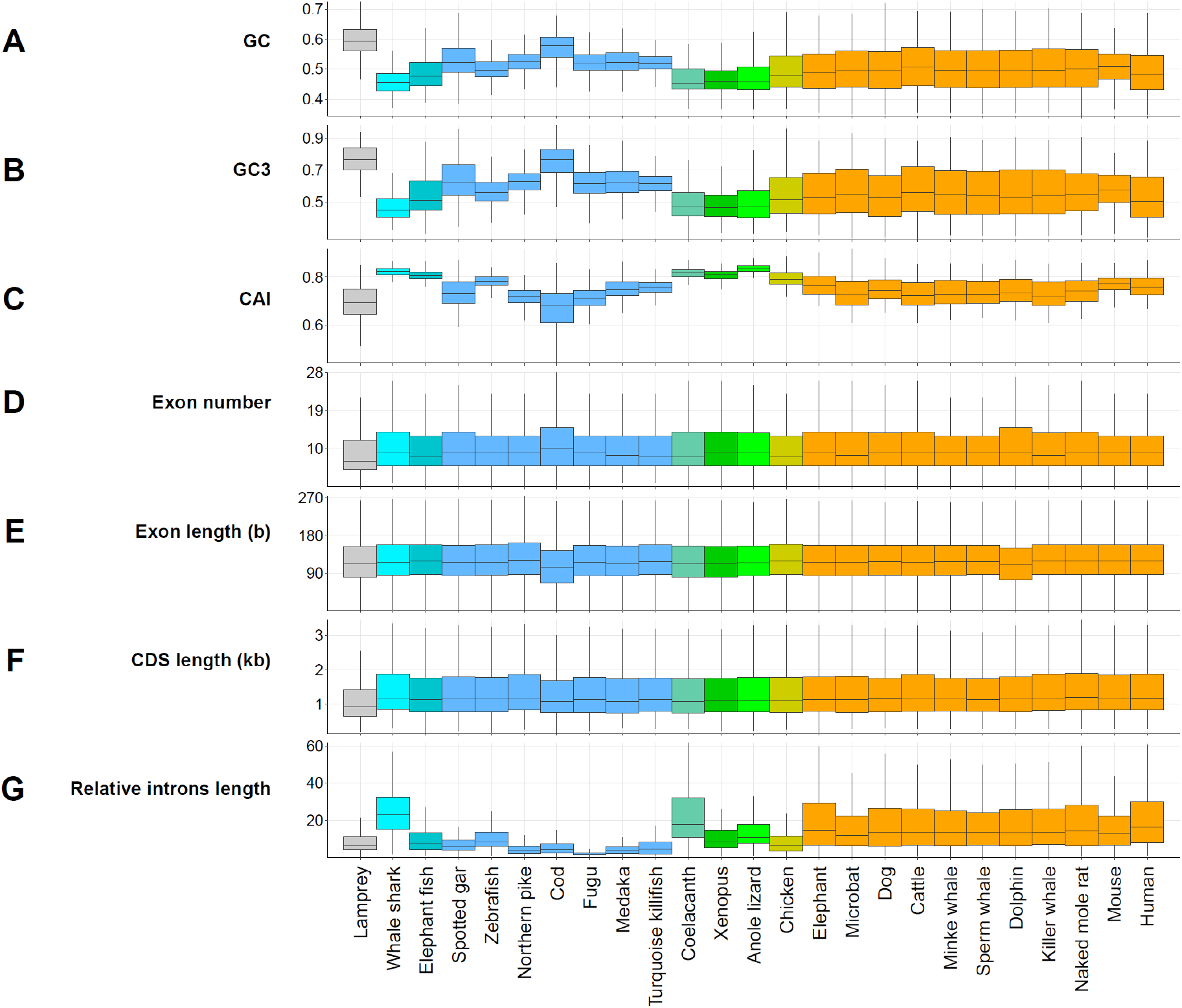
Comparison of genomic contexts in single-copy orthologous genes. 25 species were randomly selected from each class of 82 species. All comparisons (**A-G**) were performed using 275 single-copy gene families. The relative intron length (**G**) was calculated by dividing the total intron length between first coding exon and last coding exon by the CDS length. The nine colors indicate biological classifications (gray: Hyperoartia, cyan: whale shark, dark turquoise: elephant fish, light blue: Actinopterygii, aquamarine: Sarcopterygii, dark green: Amphibia, light green: Reptilia, dark yellow: Aves, orange: Mammalia).

**Figure S6.**
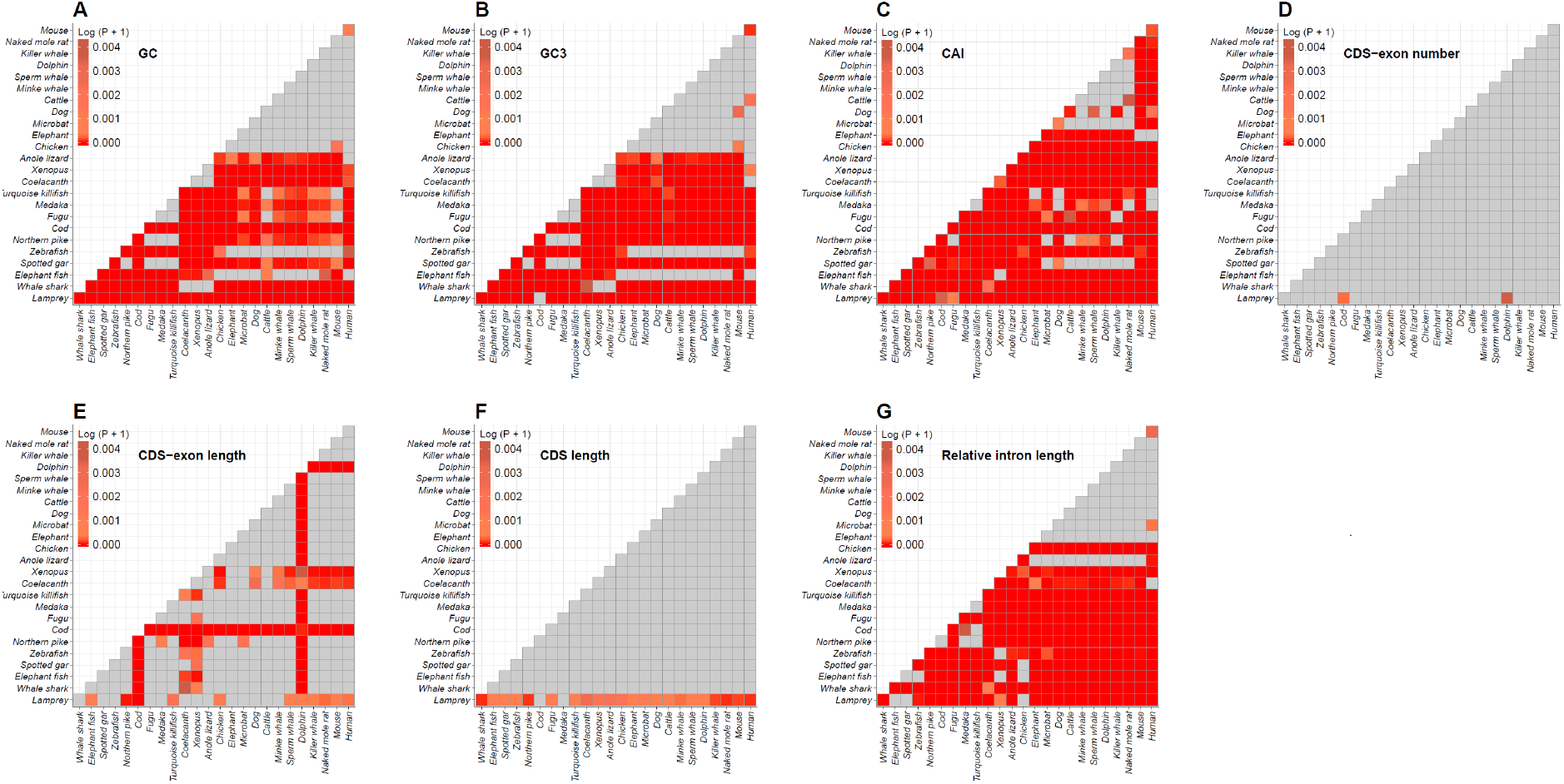
Comparison of seven genomic contexts by Wilcoxon rank sum test among 25 species. Two sided Wilcoxon rank sum tests were computed for each of the seven genomic properties in Figure S5 among 25 species. All *p*-values were adjusted using the Benjamini-Hochberg procedure and log-transformed. Deep red indicates significant adjusted *p*-values. Gray boxes indicate *p*-values higher than 0.01.

## 3. Maximum lifespan, body weight, basal metabolic rates association studies with gene size

### 3.1 Maximum lifespan data and maximum adult weight

Maximum lifespan, maximum adult weight and basal metabolic rates were downloaded from AnAge (http://genomics.senescence.info/species/), ADW (http://animaldiversity.org/), EOL (http://eol.org/), and aqW (https://www.theaquariumwiki.com/). The weight record of ten fishes were calculated by Froese, R., *et al.*’s methods, ‘length-weight relationship’ (Table S16).

**Table S16.** Maximum lifespan, weight, body temperature and basal metabolic rates of 82 species

| *Species* | *Class* | *Maximum*<br>*Lifespan* | *Reference* | *Maximum*<br>*adult weight*<br>*(g)* | *Reference* | *Average body temperature and*<br>*growth optimum temperature (°C)* | *Reference* | *Basal*<br>*metabolic*<br>*rate (W)* | *Reference* |
| --- | --- | --- | --- | --- | --- | --- | --- | --- | --- |
| *Gorilla gorilla* | Mammalia | 60.1 | AnAge | 275000 | <sup>16</sup> | 35.5 | AnAge | NA | NA |
| *Pan troglodytes* | Mammalia | 60 | ADW | 70000 | AnAge | 35.7 | AnAge | NA | NA |
| *Homo sapiens* | Mammalia | 122.5 | AnAge | 62035 | AnAge | 37 | AnAge | 82.78 | AnAge |
| *Pongo abelii* | Mammalia | 58 | ADW | 90000 | ADW | 37.6 | <sup>17</sup> | NA | NA |
| *Nomascus leucogenys* | Mammalia | 44.1 | AnAge | 7500 | EOL | 39 | <sup>18</sup> | NA | NA |
| *Macaca mulatta* | Mammalia | 40 | AnAge | 12000 | ADW | 37.3 | AnAge | NA | NA |
| *Papio anubis* | Mammalia | 45 | The Animal<br>Files | 37000 | The Animal Files | 37.3 | <sup>18,19</sup> | NA | NA |
| *Chlorocebus sabaeus* | Mammalia | 13 | ADW | 8000 | ADW | 37.35 | <sup>20</sup> | NA | NA |
| *Callithrix jacchus* | Mammalia | 22.8 | AnAge | 360 | ADW | 36 | AnAge | 0.848 | AnAge |
| *Tarsius syrichta* | Mammalia | 16 | AnAge | 165 | ADW | 33.8 | AnAge | 0.43 | AnAge |
| *Microcebus murinus* | Mammalia | 18.2 | AnAge | 71.1 | Primate Info Net | 36.1 | <sup>18,19</sup> | NA | NA |
| *Otolemur garnettii* | Mammalia | 20 | AnAge | 1345 | <sup>21</sup> | 36 | AnAge | 3.927 | AnAge |
| *Mus musculus* | Mammalia | 4 | AnAge | 30 | ADW | 36.9 | AnAge | 0.271 | AnAge |
| *Rattus norvegicus* | Mammalia | 4 | ADW | 500 | ADW | 37.1 | AnAge | 1.404 | AnAge |
| *Cavia porcellus* | Mammalia | 14 | ADW | 1100 | ADW | 39 | AnAge | 2.13 | AnAge |
| *Heterocephalus glaber* | Mammalia | 31 | AnAge | 80 | ADW | 32.1 | AnAge | 0.128 | AnAge |
| *Dipodomys ordii* | Mammalia | 9.9 | AnAge | 96 | ADW | 34.6 | AnAge | 0.339 | AnAge |
| *Ictidomys*<br>*tridecemlineatus* | Mammalia | 7.9 | AnAge | 220 | Vertebrate<br>Collection | 35.7 | AnAge | 0.983 | AnAge |
| *Ochotona princeps* | Mammalia | 7 | AnAge | 180 | Wildpro | 40.1 | AnAge | 0.932 | AnAge |
| *Oryctolagus cuniculus* | Mammalia | 9 | AnAge | 2500 | ADW | 39 | AnAge | 7.395 | AnAge |
| *Tupaia belangeri* | Mammalia | 12 | ADW | 270 | ADW | 38.8 | AnAge | NA | NA |
| *Orcinus orca* | Mammalia | 90 | AnAge | 10000000 | Seaworld | 36 | AnAge | NA | NA |
| *Tursiops truncatus* | Mammalia | 53 | ADW | 650000 | ACS | 36.9 | <sup>22</sup> | NA | NA |
| *Physeter catodon* | Mammalia | 77 | AnAge | 57000000 | MARINEBIO | 38 | WhaleForever | NA | NA |
| *Balaenoptera*<br>*acutorostrata scammoni* | Mammalia | 50 | AnAge | 13000000 | Arkive | 38 | WhaleForever | NA | NA |
| *Bos taurus* | Mammalia | 20 | AnAge | 1363000 | ADW | 38 | AnAge | 306.77 | AnAge |
| *Ovis aries* | Mammalia | 22.8 | AnAge | 200000 | ADW | 38.8 | AnAge | NA | NA |
| *Sus scrofa* | Mammalia | 27 | AnAge | 272000 | ADW | 39 | AnAge | 104.15 | AnAge |
| *Vicugna pacos* | Mammalia | 25.8 | AnAge | 84000 | Facts about Animals | 39.1 | AnAge | NA | NA |
| *Canis familiaris* | Mammalia | 29.5 | ADW | 70000 | ADW | 39 | Circadian Rhythm Lab. | NA | NA |
| *Ailuropoda melanoleuca* | Mammalia | 36.8 | AnAge | 125000 | ADW | 37 | Panda facts | NA | NA |
| *Mustela putorius furo* | Mammalia | 11.1 | AnAge | 2700 | ADW | 38.9 | Wildpro | NA | NA |
| *Felis catus* | Mammalia | 30 | AnAge | 5400 | AnAge | 38.1 | AnAge | NA | NA |
| *Erinaceus europaeus* | Mammalia | 11.7 | AnAge | 2000 | Wildpro | 34 | AnAge | 2.434 | AnAge |
| *Sorex araneus* | Mammalia | 3.2 | AnAge | 14 | ADW | 35 | AnAge | 0.348 | AnAge |
| *Myotis lucifugus* | Mammalia | 34 | AnAge | 14 | ADW | 32 | AnAge | 0.051 | AnAge |
| *Pteropus vampyrus* | Mammalia | 20.9 | AnAge | 1100 | ADW | 36.9 | AnAge | 4.486 | AnAge |
| *Equus caballus* | Mammalia | 57 | AnAge | 900000 | ADW | 38.3 | AnAge | NA | NA |
| *Echinops telfairi* | Mammalia | 19 | AnAge | 280 | ADW | 34.7 | AnAge | NA | NA |
| *Loxodonta africana* | Mammalia | 65 | AnAge | 6600000 | ELASMO | 36.2 | AnAge | NA | NA |
| *Procavia capensis* | Mammalia | 14.8 | AnAge | 4300 | ADW | 37 | AnAge | 4.954 | AnAge |
| *Choloepus hoffmanni* | Mammalia | 41 | AnAge | 12500 | Slothsanctuary | 34.4 | AnAge | 3.891 | AnAge |
| *Dasyurus novemcinctus* | Mammalia | 22.3 | AnAge | 7700 | ADW | 34.5 | AnAge | 4.655 | AnAge |
| *Sarcophilus harrisii* | Mammalia | 13 | AnAge | 12000 | ADW | 35.8 | AnAge | 8.664 | AnAge |
| *Macropus eugenii* | Mammalia | 15.1 | AnAge | 9100 | ADW | 36.5 | AnAge | 7.78 | AnAge |
| *Monodelphis domestica* | Mammalia | 5.1 | AnAge | 155 | ADW | 32.6 | AnAge | 0.335 | AnAge |
| *Ornithorhynchus anatinus* | Mammalia | 22.6 | AnAge | 2500 | AnAge | 34 | AnAge | 1.931 | AnAge |
| *Gallus gallus* | Aves | 30 | AnAge | 1450 | Arkive | 41 | <sup>23</sup> | 6.005 | AnAge |
| *Meleagris gallopavo* | Aves | 13 | AnAge | 11000 | ADW | 42.4 | <sup>24</sup> | NA | NA |
| *Anas platyrhynchos* | Aves | 29.1 | AnAge | 1580 | <sup>25</sup> | 40.2 | <sup>26</sup> | 4.068 | AnAge |
| *Taeniopygia guttata* | Aves | 12 | AnAge | 19 | FINCHINFO | 41 | AnAge | NA | NA |
| *Ficedula albicollis* | Aves | 9.8 | AnAge | 16 | Birds Natureguide | 41 | <sup>27</sup> | NA | NA |
| *Pelodiscus sinensis* | Reptilia | 20 | aqW | 2247 | EOL | 26.5 | INSECTIVORE | NA | NA |
| *Anolis carolinensis* | Reptilia | 7.2 | AnAge | 6 | ADW | 25.5 | The spruce | NA | NA |
| *Xenopus tropicalis* | Amphibia | 20 | Tropical-fish-keeping | 26 | <sup>28</sup> | 25.5 | Xenopus Express | NA | NA |
| *Latimeria chalumnae* | Sarcopterygii | 48 | AnAge | 95000 | Fishbase | 19 | VIMS | NA | NA |
| *Poecilia reticulata* | Actinopterygii | 5 | AnAge | 4.13 | Fishbase <sup>29</sup> | 22.5 | SERIOUSLY FISH | NA | NA |
| *Poecilia formosa* | Actinopterygii | 3 | <sup>30</sup> | 11.3 | Fishbase <sup>29</sup> | 25 | The aquarium guide | NA | NA |
| *Xiphophorus maculatus* | Actinopterygii | 5 | aqW | 2 | Fishbase <sup>29</sup> | 22 | Tropical Fish Site | NA | NA |
| *Nothobranchius furzeri* | Actinopterygii | 1.1 | AnAge | 1.3 | Fishbase <sup>29</sup> | 27.8 | WildNothos | NA | NA |
| *Oryzias latipes* | Actinopterygii | 5 | AnAge | 0.2 | Fishbase <sup>29</sup> | 19 | SERIOUSLY FISH | NA | NA |
| *Oreochromis niloticus* | Actinopterygii | 9 | AnAge | 4300 | Fishbase | 33.5 | FAO | NA | NA |
| *Takifugu rubripes* | Actinopterygii | 9 | IUCN | 5754 | Fishbase <sup>29</sup> | 25 | <sup>31</sup> | NA | NA |
| *Tetraodon nigroviridis* | Actinopterygii | 10 | aqW | 96 | Fishbase <sup>29</sup> | 26 | SERIOUSLY FISH | NA | NA |
| *Gasterosteus aculeatus* | Actinopterygii | 8 | AnAge | 16 | Fishbase <sup>29</sup> | 21 | <sup>32</sup> | NA | NA |
| *Gadus morhua* | Actinopterygii | 25 | AnAge | 96000 | Fishbase | 12 | <sup>33</sup> | NA | NA |
| *Oncorhynchus kisutch* | Actinopterygii | 5 | AnAge | 15200 | Fishbase | 12 | <sup>34</sup> | NA | NA |
| *Salmo salar* | Actinopterygii | 13 | AnAge | 46800 | Fishbase | 8 | USGS | NA | NA |
| *Esox lucius* | Actinopterygii | 30 | AnAge | 28400 | AnAge | 20.5 | <sup>35</sup> | NA | NA |
| *Danio rerio* | Actinopterygii | 5.5 | AnAge | 1 | Fishbase <sup>29</sup> | 28.5 | <sup>36</sup> | NA | NA |
| *Cyprinus carpio* | Actinopterygii | 47 | AnAge | 40100 | Fishbase | 22.5 | FAO | NA | NA |
| *Astyanax mexicanus* | Actinopterygii | 8 | aqW | 21 | Fishbase <sup>29</sup> | 24 | <sup>37</sup> | NA | NA |
| *Ictalurus punctatus* | Actinopterygii | 16 | Fishbase | 13733 | AnAge | 27 | FAO | NA | NA |
| *Lepisosteus oculatus* | Actinopterygii | 18 | AnAge | 4400 | Fishbase | 16 | SERIOUSLY FISH | NA | NA |
| *Callorhynchus milii* | Chondrichthyes | 20 | IUCN | 4000 | ADW | 14 | WHRHSmarin ebiology | NA | NA |
| *Rhincodon typus* | Chondrichthyes | 80.4 | <sup>38</sup> | 42000000 | IUCN | 25 | Arkive | NA | NA |
| *Petromyzon marinus* | Hyperoartia | 9 | AnAge | 2500 | AnAge | 20 | <sup>39</sup> | NA | NA |
| *Ciona intestinalis* | Ascidiacea | 1 | ADW | 0 | NA | NA | NA | NA | NA |
| *Ciona savignyi* | Ascidiacea | 1 | ADW | 0 | NA | NA | NA | NA | NA |
| *Caenorhabditis elegans* | Chromadorea | 0.16 | AnAge | 0 | NA | NA | NA | NA | NA |
| *Drosophila melanogaster* | Insecta | 0.3 | AnAge | 0 | NA | NA | NA | NA | NA |
| *Saccharomyces cerevisiae* | Saccharomycetes | 0.04 | AnAge | 0 | NA | NA | NA | NA | NA |

### 3.2 Basal metabolic rates calculation

We calculated basal metabolic rates (BMRs) of 82 species using Gillooly’s equation^40^ based on maximum adult weight and average body temperature (or growth optimum temperature for cold-blooded animal) (Table S16). The calculated BMRs were compared with published BMRs from the AnAge database (http://genomics.senescence.info/) using the Spearman’s rank correlation coefficient (Figure S7). The Calculated Gillooly’s BMRs^40^ were significantly correlated with BMRs downloaded from the AnAge database.

**Figure S7.**
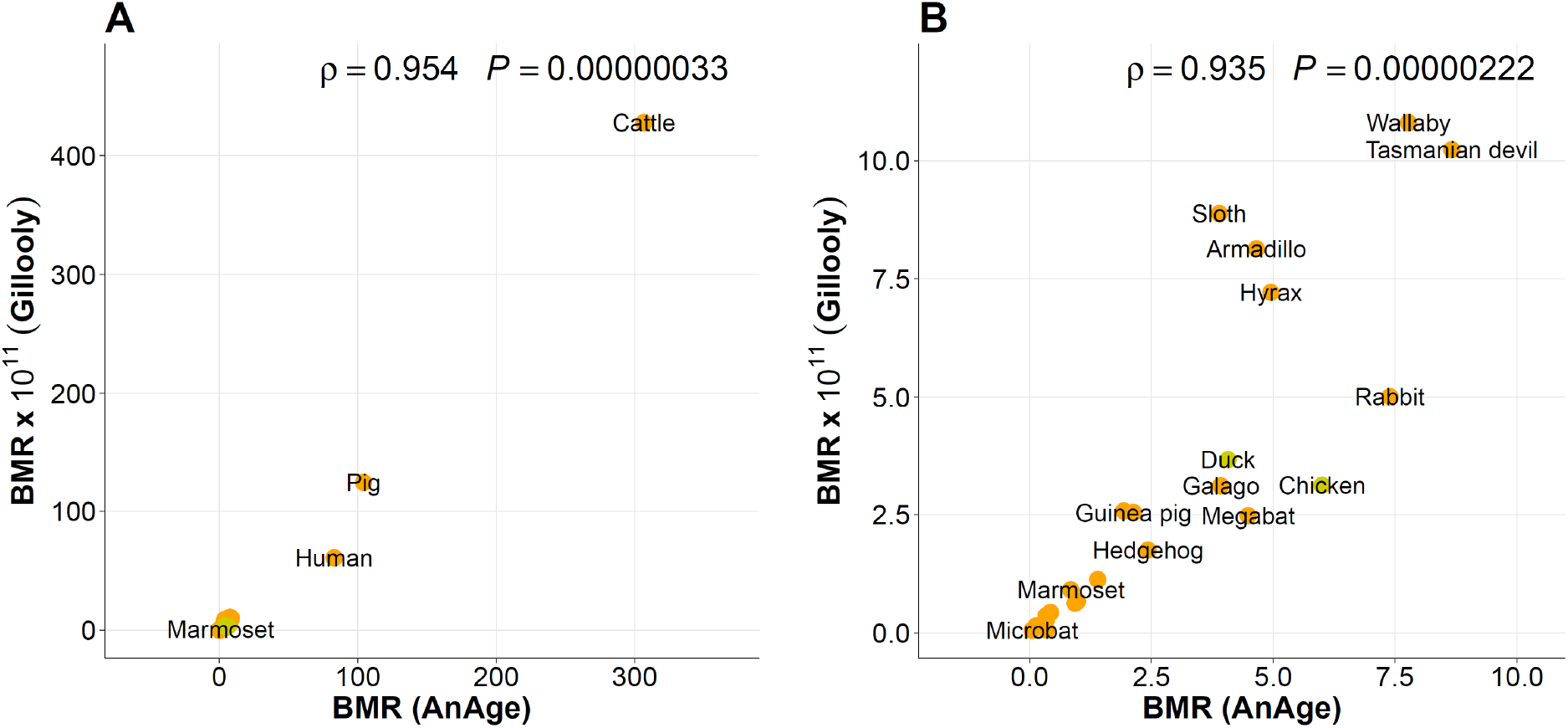
Correlation between AnAge’s BMR and calculated BMR. We downloaded the BMRs of 27 species from AnAge (http://genomics.senescence.info/, Table S16) and also calculated BMRs using Gillooly’s equation^40^. (**A**) The correlation test with 27 species. (**B**) The correlation test without cattle, pig, and human, which have extremely high BMR.

**Figure S8.**
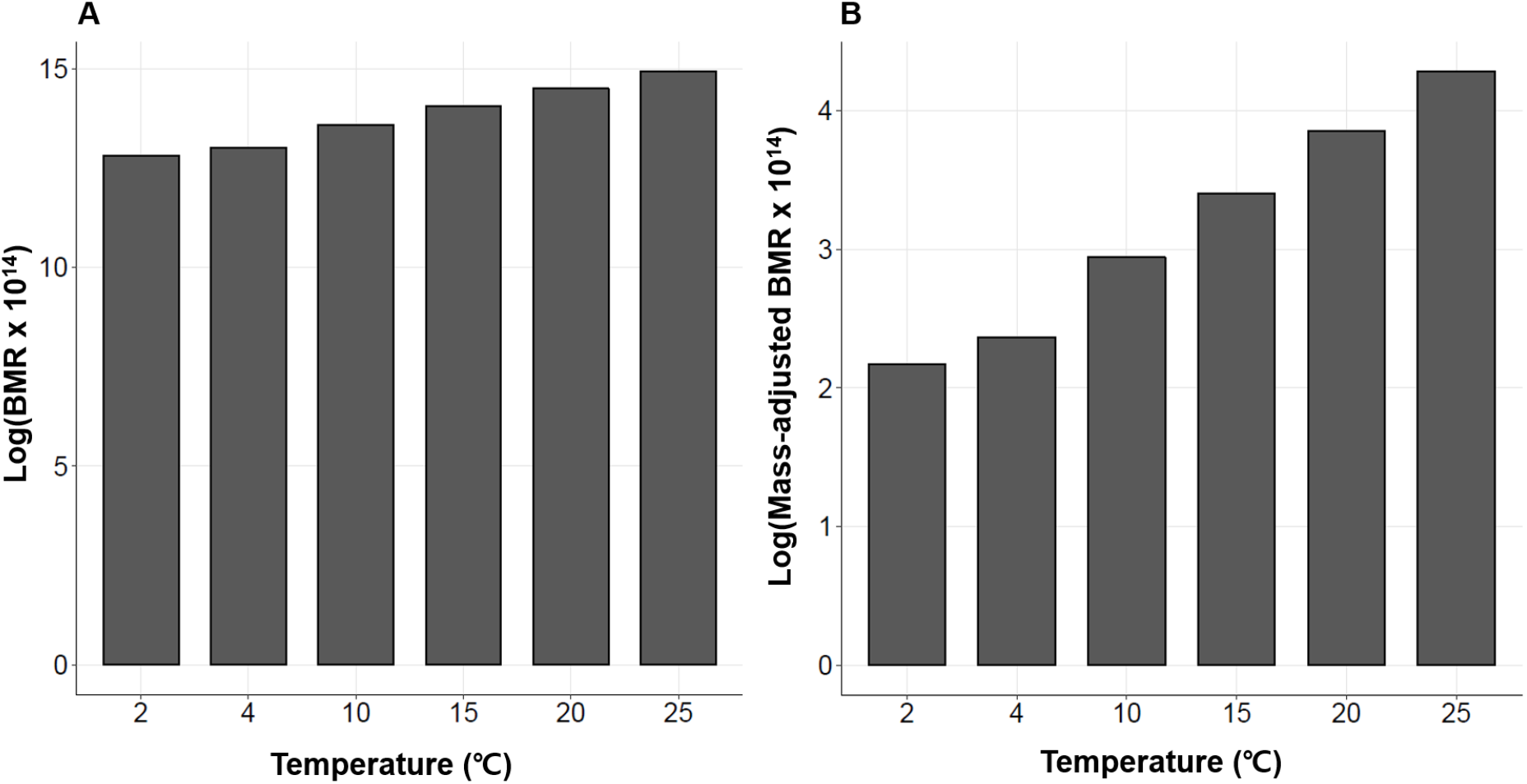
Changes of BMR and mass-adjusted BMR by temperature. Using the Gillooly’s equation^40^, we calculated the BMR (**A**) and the mass-adjusted BMR (**B**) using six temperatures selected within the range of temperatures at which whale shark (which dives to deep cold waters) is known to live. Both BMR and mass-adjusted BMR were multiplied by 10^14^ and log-transformed.

**Table S17.** Spearman’s rho rank correlations between 22 properties in each *Eukaryota, Mammalia* and *Actinopterygii*

| Property A | Property B | *Eukaryota* |  | *Mammalia* |  | *Actinopterygii* |  |
| --- | --- | --- | --- | --- | --- | --- | --- |
|  |  | Rho | *p* | Rho | *p* | Rho | *p* |
| Weight | Relative intron length | 0.517 | 6.7E-07 | 0.385 | 7.5E-03 | 0.020 | 9.4E-01 |
| Weight | mRNA length | 0.094 | 4.0E-01 | 0.223 | 1.3E-01 | 0.042 | 8.7E-01 |
| Weight | Intron length between first and last exons | 0.482 | 5.2E-06 | 0.343 | 1.8E-02 | -0.092 | 7.2E-01 |
| Weight | Intron length between first and last coding exons | 0.503 | 1.4E-06 | 0.351 | 1.6E-02 | -0.084 | 7.4E-01 |
| Weight | GC3 | 0.081 | 4.7E-01 | 0.067 | 6.5E-01 | 0.242 | 3.3E-01 |
| Weight | GC contents | 0.137 | 2.2E-01 | 0.079 | 6.0E-01 | 0.250 | 3.2E-01 |
| Weight | Exon+Intron length between first and last exon | 0.463 | 1.4E-05 | 0.342 | 1.9E-02 | -0.080 | 7.5E-01 |
| Weight | Exon+Intron length between first and last coding exon | 0.510 | 1.0E-06 | 0.388 | 7.0E-03 | -0.060 | 8.1E-01 |
| Weight | Exon number | 0.079 | 4.8E-01 | 0.183 | 2.2E-01 | -0.071 | 7.8E-01 |
| Weight | Exon length | -0.106 | 3.4E-01 | 0.174 | 2.4E-01 | 0.102 | 6.9E-01 |
| Weight | Controlled intron length | -0.164 | 1.4E-01 | 0.309 | 3.5E-02 | -0.344 | 1.6E-01 |
| Weight | CDS length | 0.074 | 5.1E-01 | 0.284 | 5.3E-02 | -0.082 | 7.5E-01 |
| Weight | CAI | 0.109 | 3.3E-01 | 0.004 | 9.8E-01 | -0.353 | 1.5E-01 |
| Weight | 5'UTR length | 0.060 | 6.4E-01 | -0.141 | 4.4E-01 | -0.025 | 9.3E-01 |
| Weight | 3'UTR length | 0.216 | 8.7E-02 | 0.029 | 8.7E-01 | 0.147 | 5.7E-01 |
| Temperature | Weight | 0.210 | 6.7E-02 | 0.392 | 6.5E-03 | -0.409 | 9.2E-02 |
| Temperature | Relative intron length | 0.397 | 3.5E-04 | 0.161 | 2.8E-01 | -0.064 | 8.0E-01 |
| Temperature | mRNA length | -0.196 | 9.0E-02 | 0.181 | 2.2E-01 | 0.092 | 7.2E-01 |
| Temperature | Intron length between first and last exons | 0.399 | 3.5E-04 | 0.190 | 2.0E-01 | 0.033 | 9.0E-01 |
| Temperature | Intron length between first and last coding exons | 0.389 | 4.8E-04 | 0.187 | 2.1E-01 | 0.019 | 9.4E-01 |
| Temperature | Genome size | 0.381 | 6.4E-04 | -0.028 | 8.5E-01 | -0.206 | 4.1E-01 |
| Temperature | GC3 | -0.149 | 2.0E-01 | 0.169 | 2.6E-01 | -0.495 | 3.7E-02 |
| Temperature | GC contents | -0.069 | 5.5E-01 | 0.189 | 2.0E-01 | -0.554 | 1.7E-02 |
| Temperature | Exon+Intron length between first and last exon | 0.289 | 1.1E-02 | 0.185 | 2.1E-01 | 0.054 | 8.3E-01 |
| Temperature | Exon+Intron length between first and last coding exon | 0.315 | 5.3E-03 | 0.187 | 2.1E-01 | 0.027 | 9.2E-01 |
| Temperature | Exon number | -0.103 | 3.7E-01 | 0.172 | 2.5E-01 | 0.218 | 3.9E-01 |
| Temperature | Exon length | -0.083 | 4.7E-01 | 0.106 | 4.8E-01 | 0.087 | 7.3E-01 |
| Temperature | Controlled intron length | -0.155 | 1.8E-01 | 0.173 | 2.5E-01 | 0.327 | 1.9E-01 |
| Temperature | CDS length | -0.155 | 1.8E-01 | 0.187 | 2.1E-01 | 0.423 | 8.0E-02 |
| Temperature | CAI | 0.249 | 2.9E-02 | -0.152 | 3.1E-01 | 0.607 | 7.6E-03 |
| Temperature | 5'UTR length | -0.192 | 1.4E-01 | 0.077 | 6.8E-01 | 0.064 | 8.1E-01 |
| Temperature | 3'UTR length | 0.022 | 8.7E-01 | 0.205 | 2.6E-01 | -0.038 | 8.9E-01 |
| Relative intron length | mRNA length | 0.194 | 8.2E-02 | 0.737 | 3.4E-09 | 0.523 | 2.8E-02 |
| Relative intron length | Intron length between first and last exons | 0.944 | 0.0E+0 | 0.914 | 0.0E+0 | 0.975 | 8.4E-06 |
| Relative intron length | GC3 | -0.207 | 6.2E-02 | -0.280 | 5.7E-02 | -0.447 | 6.5E-02 |
| Relative intron length | GC contents | -0.101 | 3.7E-01 | -0.178 | 2.3E-01 | -0.391 | 1.1E-01 |
| Relative intron length | Exon+Intron length between first and last exon | 0.874 | 0.0E+0 | 0.882 | 0.0E+0 | 0.959 | 5.6E-06 |
| Relative intron length | Exon+Intron length between first and last coding exon | 0.917 | 0.0E+0 | 0.891 | 0.0E+0 | 0.971 | 7.8E-06 |
| Relative intron length | Exon number | -0.093 | 4.1E-01 | 0.197 | 1.9E-01 | -0.379 | 1.2E-01 |
| Relative intron length | Exon length | -0.032 | 7.8E-01 | 0.579 | 2.0E-05 | 0.556 | 1.7E-02 |
| Relative intron length | Controlled intron length | -0.074 | 5.1E-01 | 0.713 | 9.6E-08 | 0.321 | 1.9E-01 |
| Relative intron length | CDS length | 0.076 | 5.0E-01 | 0.664 | 3.6E-07 | 0.259 | 3.0E-01 |
| Relative intron length | CAI | 0.497 | 2.8E-06 | 0.451 | 1.6E-03 | 0.302 | 2.2E-01 |
| Relative intron length | 5'UTR length | 0.214 | 9.0E-02 | 0.368 | 3.8E-02 | 0.478 | 5.4E-02 |
| Relative intron length | 3'UTR length | 0.407 | 8.4E-04 | 0.327 | 6.9E-02 | 0.495 | 4.5E-02 |
| mRNA length | GC3 | 0.091 | 4.2E-01 | 0.011 | 9.4E-01 | -0.292 | 2.4E-01 |
| mRNA length | GC contents | 0.085 | 4.5E-01 | 0.128 | 3.9E-01 | -0.255 | 3.1E-01 |
| mRNA length | Exon number | 0.371 | 6.6E-04 | 0.294 | 4.5E-02 | -0.122 | 6.3E-01 |
| mRNA length | Exon length | 0.618 | 8.2E-10 | 0.830 | 5.2E-13 | 0.783 | 1.2E-04 |
| mRNA length | CDS length | 0.836 | 2.7E-22 | 0.815 | 3.1E-12 | 0.724 | 6.8E-04 |
| mRNA length | CAI | -0.021 | 8.5E-01 | 0.079 | 6.0E-01 | 0.230 | 3.6E-01 |
| mRNA length | 5'UTR length | 0.705 | 7.9E-11 | 0.451 | 9.6E-03 | 0.583 | 1.6E-02 |
| mRNA length | 3'UTR length | 0.615 | 6.6E-08 | 0.404 | 2.2E-02 | 0.581 | 1.6E-02 |
| Maximum lifespan | Weight | 0.791 | 1.0E-18 | 0.720 | 1.2E-08 | 0.801 | 6.4E-05 |
| Maximum lifespan | Temperature | 0.246 | 3.1E-02 | 0.049 | 7.4E-01 | -0.239 | 3.4E-01 |
| Maximum lifespan | Relative intron length | 0.581 | 1.1E-08 | 0.399 | 5.5E-03 | -0.076 | 7.7E-01 |
| Maximum lifespan | mRNA length | 0.054 | 6.3E-01 | 0.278 | 5.9E-02 | -0.181 | 4.7E-01 |
| Maximum lifespan | Mass adjusted BMR | -0.553 | 2.2E-07 | -0.748 | 1.6E-09 | -0.718 | 1.2E-03 |
| Maximum lifespan | Intron length between first and last exons | 0.523 | 5.3E-07 | 0.340 | 1.9E-02 | -0.150 | 5.5E-01 |
| Maximum lifespan | Intron length between first and last coding exons | 0.554 | 6.7E-08 | 0.357 | 1.4E-02 | -0.154 | 5.4E-01 |
| Maximum lifespan | Genome size | 0.407 | 1.5E-04 | -0.151 | 3.1E-01 | 0.097 | 7.0E-01 |
| Maximum lifespan | GC3 | -0.023 | 8.4E-01 | 0.059 | 7.0E-01 | -0.017 | 9.5E-01 |
| Maximum lifespan | GC contents | 0.056 | 6.2E-01 | 0.111 | 4.6E-01 | -0.031 | 9.0E-01 |
| Maximum lifespan | Exon+Intron length between first and last exon | 0.494 | 2.8E-06 | 0.380 | 8.4E-03 | -0.151 | 5.5E-01 |
| Maximum lifespan | Exon+Intron length between first and last coding exon | 0.557 | 5.6E-08 | 0.421 | 3.3E-03 | -0.118 | 6.4E-01 |
| Maximum lifespan | Exon number | 0.064 | 5.7E-01 | 0.243 | 9.9E-02 | -0.071 | 7.8E-01 |
| Maximum lifespan | Exon length | -0.182 | 1.0E-01 | 0.160 | 2.8E-01 | -0.121 | 6.3E-01 |
| Maximum lifespan | Controlled maximum lifespan | 0.349 | 1.3E-03 | 0.254 | 8.5E-02 | 0.162 | 5.2E-01 |
| Maximum lifespan | Controlled intron length | -0.118 | 2.9E-01 | 0.362 | 1.3E-02 | -0.161 | 5.2E-01 |
| Maximum lifespan | CDS length | 0.035 | 7.5E-01 | 0.329 | 2.4E-02 | -0.196 | 4.4E-01 |
| Maximum lifespan | CAI | 0.253 | 2.2E-02 | 0.043 | 7.8E-01 | -0.112 | 6.6E-01 |
| Maximum lifespan | Basal metabolic rate | 0.756 | 2.7E-15 | 0.709 | 2.5E-08 | 0.813 | 7.3E-05 |
| Maximum lifespan | 5'UTR length | -0.018 | 8.9E-01 | -0.143 | 4.4E-01 | -0.133 | 6.1E-01 |
| Maximum lifespan | 3'UTR length | 0.145 | 2.5E-01 | -0.098 | 5.9E-01 | 0.058 | 8.3E-01 |
| Mass adjusted BMR | Weight | -0.863 | 1.3E-23 | -0.981 | 9.3E-34 | -0.957 | 1.8E-09 |
| Mass adjusted BMR | Temperature | 0.275 | 1.6E-02 | -0.246 | 9.6E-02 | 0.672 | 3.2E-03 |
| Mass adjusted BMR | Relative intron length | -0.142 | 2.2E-01 | -0.378 | 9.2E-03 | 0.032 | 9.1E-01 |
| Mass adjusted BMR | mRNA length | -0.090 | 4.4E-01 | -0.209 | 1.6E-01 | 0.125 | 6.3E-01 |
| Mass adjusted BMR | Intron length between first and last exons | -0.124 | 2.9E-01 | -0.333 | 2.3E-02 | 0.091 | 7.3E-01 |
| Mass adjusted BMR | Intron length between first and last coding exons | -0.149 | 2.0E-01 | -0.341 | 2.0E-02 | 0.081 | 7.6E-01 |
| Mass adjusted BMR | Genome size | -0.005 | 9.7E-01 | 0.064 | 6.7E-01 | -0.289 | 2.6E-01 |
| Mass adjusted BMR | GC3 | -0.093 | 4.3E-01 | -0.058 | 7.0E-01 | -0.304 | 2.4E-01 |
| Mass adjusted BMR | GC contents | -0.101 | 3.9E-01 | -0.072 | 6.3E-01 | -0.331 | 1.9E-01 |
| Mass adjusted BMR | Exon+Intron length between first and last exon | -0.161 | 1.7E-01 | -0.328 | 2.5E-02 | 0.100 | 7.0E-01 |
| Mass adjusted BMR | Exon+Intron length between first and last coding exon | -0.201 | 8.2E-02 | -0.376 | 9.6E-03 | 0.066 | 8.0E-01 |
| Mass adjusted BMR | Exon number | 0.012 | 9.2E-01 | -0.159 | 2.9E-01 | 0.184 | 4.8E-01 |
| Mass adjusted BMR | Exon length | -0.094 | 4.2E-01 | -0.156 | 3.0E-01 | -0.006 | 9.8E-01 |
| Mass adjusted BMR | Controlled intron length | -0.063 | 5.9E-01 | -0.290 | 4.9E-02 | 0.478 | 5.4E-02 |
| Mass adjusted BMR | CDS length | -0.074 | 5.2E-01 | -0.264 | 7.3E-02 | 0.275 | 2.9E-01 |
| Mass adjusted BMR | CAI | 0.091 | 4.3E-01 | -0.006 | 9.7E-01 | 0.463 | 6.3E-02 |
| Mass adjusted BMR | 5'UTR length | 0.062 | 6.4E-01 | 0.130 | 4.8E-01 | 0.109 | 6.9E-01 |
| Mass adjusted BMR | 3'UTR length | -0.011 | 9.3E-01 | -0.027 | 8.9E-01 | -0.062 | 8.2E-01 |
| Intron length between first and last exons | mRNA length | 0.424 | 7.8E-05 | 0.896 | 1.9E-17 | 0.554 | 1.9E-02 |
| Intron length between first and last exons | GC3 | -0.165 | 1.4E-01 | -0.104 | 4.9E-01 | -0.546 | 2.1E-02 |
| Intron length between first and last exons | GC contents | -0.062 | 5.8E-01 | 0.005 | 9.7E-01 | -0.494 | 3.9E-02 |
| Intron length between first and last exons | Exon+Intron length between first and last exon | 0.970 | 0.0E+0<br>0 | 0.966 | 0.0E+0<br>0 | 0.988 | 9.8E-06 |
| Intron length between first and last exons | Exon+Intron length between first and last coding exon | 0.979 | 0.0E+0<br>0 | 0.944 | 0.0E+0<br>0 | 0.994 | 1.0E-05 |
| Intron length between first and last exons | Exon number | 0.089 | 4.3E-01 | 0.339 | 2.0E-02 | -0.363 | 1.4E-01 |
| Intron length between first and last exons | Exon length | 0.155 | 1.7E-01 | 0.765 | 3.9E-10 | 0.580 | 1.2E-02 |
| Intron length between first and last exons | CDS length | 0.337 | 2.1E-03 | 0.821 | 1.7E-12 | 0.336 | 1.7E-01 |
| Intron length between first and last exons | CAI | 0.440 | 4.8E-05 | 0.261 | 7.7E-02 | 0.401 | 1.0E-01 |
| Intron length between first and last exons | 5'UTR length | 0.269 | 3.2E-02 | 0.407 | 2.1E-02 | 0.493 | 4.7E-02 |
| Intron length between first and last exons | 3'UTR length | 0.465 | 1.1E-04 | 0.367 | 3.9E-02 | 0.490 | 4.8E-02 |
| Intron length between first and last coding exons | Relative intron length | 0.960 | 0.0E+0<br>0 | 0.930 | 0.0E+0<br>0 | 0.979 | 8.9E-06 |
| Intron length between first and last coding exons | mRNA length | 0.372 | 6.3E-04 | 0.862 | 7.7E-15 | 0.544 | 2.1E-02 |
| Intron length between first and last coding exons | Intron length between first and last exons | 0.995 | 0.0E+0<br>0 | 0.990 | 0.0E+0<br>0 | 0.998 | 1.1E-05 |
| Intron length between first and last coding exons | GC3 | -0.188 | 9.2E-02 | -0.130 | 3.8E-01 | -0.513 | 3.1E-02 |
| Intron length between first and last coding exons | GC contents | -0.084 | 4.6E-01 | -0.029 | 8.5E-01 | -0.461 | 5.6E-02 |
| Intron length between first and last coding exons | Exon+Intron length between first and last exon | 0.954 | 0.0E+0<br>0 | 0.949 | 0.0E+0<br>0 | 0.986 | 9.6E-06 |
| Intron length between first and last coding exons | Exon+Intron length between first and last coding exon | 0.976 | 0.0E+0<br>0 | 0.945 | 0.0E+0<br>0 | 0.996 | 1.0E-05 |
| Intron length between first and last coding exons | Exon number | 0.039 | 7.3E-01 | 0.341 | 1.9E-02 | -0.334 | 1.8E-01 |
| Intron length between first and last coding exons | Exon length | 0.127 | 2.6E-01 | 0.752 | 1.1E-09 | 0.572 | 1.3E-02 |
| Intron length between first and last coding exons | Controlled intron length | 0.076 | 5.0E-01 | 0.833 | 0.0E+0<br>0 | 0.397 | 1.0E-01 |
| Intron length between first and last coding exons | CDS length | 0.256 | 2.0E-02 | 0.821 | 1.7E-12 | 0.338 | 1.7E-01 |
| Intron length between first and last coding exons | CAI | 0.463 | 1.5E-05 | 0.289 | 4.9E-02 | 0.366 | 1.4E-01 |
| Intron length between first and last coding exons | 5'UTR length | 0.240 | 5.6E-02 | 0.353 | 4.8E-02 | 0.493 | 4.7E-02 |
| Intron length between first and last coding exons | 3'UTR length | 0.435 | 3.2E-04 | 0.309 | 8.6E-02 | 0.490 | 4.8E-02 |
| Genome size | Weight | 0.419 | 8.9E-05 | -0.078 | 6.0E-01 | 0.193 | 4.4E-01 |
| Genome size | Relative intron length | 0.707 | 0.0E+0<br>0 | 0.132 | 3.7E-01 | 0.697 | 1.8E-03 |
| Genome size | mRNA length | -0.123 | 2.7E-01 | -0.122 | 4.1E-01 | 0.216 | 3.9E-01 |
| Genome size | Intron length between first and last exons | 0.572 | 4.5E-08 | -0.007 | 9.6E-01 | 0.678 | 2.6E-03 |
| Genome size | Intron length between first and last coding exons | 0.595 | 6.5E-09 | 0.020 | 8.9E-01 | 0.662 | 3.6E-03 |
| Genome size | GC3 | -0.125 | 2.6E-01 | -0.473 | 9.0E-04 | -0.486 | 4.3E-02 |
| Genome size | GC contents | -0.042 | 7.1E-01 | -0.434 | 2.5E-03 | -0.418 | 8.6E-02 |
| Genome size | Exon+Intron length between first and last exon | 0.450 | 3.0E-05 | -0.083 | 5.8E-01 | 0.643 | 5.0E-03 |
| Genome size | Exon+Intron length between first and last coding exon | 0.489 | 4.3E-06 | -0.106 | 4.8E-01 | 0.631 | 6.1E-03 |
| Genome size | Exon number | -0.267 | 1.5E-02 | -0.339 | 2.0E-02 | -0.804 | 5.8E-05 |
| Genome size | Exon length | -0.324 | 3.0E-03 | -0.295 | 4.4E-02 | 0.401 | 9.9E-02 |
| Genome size | Controlled intron length | -0.610 | 1.3E-09 | -0.456 | 1.4E-03 | -0.327 | 1.9E-01 |
| Genome size | CDS length | -0.284 | 9.7E-03 | -0.338 | 2.0E-02 | -0.158 | 5.3E-01 |
| Genome size | CAI | 0.327 | 2.8E-03 | 0.356 | 1.4E-02 | 0.333 | 1.8E-01 |
| Genome size | 5'UTR length | 0.238 | 5.8E-02 | 0.431 | 1.4E-02 | 0.238 | 3.6E-01 |
| Genome size | 3'UTR length | 0.365 | 3.0E-03 | 0.407 | 2.2E-02 | 0.216 | 4.0E-01 |
| GC3 | Exon number | 0.447 | 2.5E-05 | 0.530 | 1.3E-04 | 0.421 | 8.2E-02 |
| GC3 | Exon length | -0.121 | 2.8E-01 | -0.002 | 9.9E-01 | -0.248 | 3.2E-01 |
| GC3 | CDS length | 0.216 | 5.1E-02 | 0.216 | 1.5E-01 | -0.237 | 3.4E-01 |
| GC3 | CAI | -0.841 | 0.0E+0<br>0 | -0.895 | 0.0E+0<br>0 | -0.936 | 0.0E+0<br>0 |
| GC3 | 5'UTR length | 0.012 | 9.2E-01 | -0.309 | 8.5E-02 | -0.314 | 2.2E-01 |
| GC3 | 3'UTR length | -0.093 | 4.7E-01 | -0.250 | 1.7E-01 | -0.265 | 3.0E-01 |
| GC contents | GC3 | 0.975 | 0.0E+0<br>0 | 0.969 | 0.0E+0<br>0 | 0.992 | 1.0E-05 |
| GC contents | Exon number | 0.444 | 2.9E-05 | 0.598 | 8.9E-06 | 0.375 | 1.3E-01 |
| GC contents | Exon length | -0.133 | 2.3E-01 | 0.046 | 7.6E-01 | -0.216 | 3.9E-01 |
| GC contents | CDS length | 0.193 | 8.2E-02 | 0.297 | 4.3E-02 | -0.264 | 2.9E-01 |
| GC contents | CAI | -0.799 | 0.0E+0<br>0 | -0.855 | 0.0E+0<br>0 | -0.948 | 3.2E-06 |
| GC contents | 5'UTR length | 0.013 | 9.2E-01 | -0.195 | 2.9E-01 | -0.277 | 2.8E-01 |
| GC contents | 3'UTR length | -0.050 | 7.0E-01 | -0.131 | 4.7E-01 | -0.223 | 3.9E-01 |
| Exon+Intron length between first and last exon | mRNA length | 0.579 | 1.5E-08 | 0.930 | 4.0E-21 | 0.600 | 9.9E-03 |
| Exon+Intron length between first and last exon | GC3 | -0.125 | 2.6E-01 | -0.036 | 8.1E-01 | -0.554 | 1.9E-02 |
| Exon+Intron length between first and last exon | GC contents | -0.034 | 7.7E-01 | 0.078 | 6.0E-01 | -0.505 | 3.5E-02 |
| Exon+Intron length between first and last exon | Exon number | 0.188 | 9.3E-02 | 0.368 | 1.1E-02 | -0.354 | 1.5E-01 |
| Exon+Intron length between first and last exon | Exon length | 0.255 | 2.2E-02 | 0.776 | 1.5E-10 | 0.564 | 1.5E-02 |
| Exon+Intron length between first and last exon | CDS length | 0.490 | 3.4E-06 | 0.862 | 7.6E-15 | 0.352 | 1.5E-01 |
| Exon+Intron length between first and last exon | CAI | 0.383 | 4.5E-04 | 0.199 | 1.8E-01 | 0.408 | 9.4E-02 |
| Exon+Intron length between first and last exon | 5'UTR length | 0.331 | 7.5E-03 | 0.383 | 3.1E-02 | 0.478 | 5.4E-02 |
| Exon+Intron length between first and last exon | 3'UTR length | 0.501 | 2.4E-05 | 0.338 | 5.9E-02 | 0.480 | 5.3E-02 |
| Exon+Intron length between first and last coding exon | mRNA length | 0.452 | 2.3E-05 | 0.835 | 2.9E-13 | 0.558 | 1.8E-02 |
| Exon+Intron length between first and last coding exon | GC3 | -0.145 | 1.9E-01 | -0.005 | 9.8E-01 | -0.529 | 2.6E-02 |
| Exon+Intron length between first and last coding exon | GC contents | -0.042 | 7.1E-01 | 0.102 | 4.9E-01 | -0.480 | 4.6E-02 |
| Exon+Intron length between first and last coding exon | Exon+Intron length between first and last exon | 0.981 | 0.0E+0<br>0 | 0.964 | 0.0E+0<br>0 | 0.990 | 9.9E-06 |
| Exon+Intron length between first and last coding exon | Exon number | 0.148 | 1.9E-01 | 0.497 | 3.8E-04 | -0.307 | 2.2E-01 |
| Exon+Intron length between first and last coding exon | Exon length | 0.165 | 1.4E-01 | 0.697 | 5.4E-08 | 0.564 | 1.5E-02 |
| Exon+Intron length between first and last coding exon | CDS length | 0.370 | 6.3E-04 | 0.887 | 1.1E-16 | 0.366 | 1.4E-01 |
| Exon+Intron length between first and last coding exon | CAI | 0.419 | 1.1E-04 | 0.215 | 1.5E-01 | 0.375 | 1.3E-01 |
| Exon+Intron length between first and last coding exon | 5'UTR length | 0.246 | 5.0E-02 | 0.285 | 1.1E-01 | 0.488 | 4.9E-02 |
| Exon+Intron length between first and last coding exon | 3'UTR length | 0.441 | 2.7E-04 | 0.256 | 1.6E-01 | 0.498 | 4.4E-02 |
| Exon number | Exon length | -0.074 | 5.1E-01 | 0.220 | 1.4E-01 | -0.265 | 2.9E-01 |
| Exon number | CDS length | 0.643 | 7.6E-11 | 0.604 | 7.1E-06 | 0.303 | 2.2E-01 |
| Exon number | 5'UTR length | 0.215 | 8.8E-02 | -0.038 | 8.4E-01 | 0.172 | 5.1E-01 |
| Exon number | 3'UTR length | 0.219 | 8.2E-02 | 0.009 | 9.6E-01 | 0.193 | 4.6E-01 |
| Exon length | CDS length | 0.518 | 6.1E-07 | 0.801 | 1.4E-11 | 0.701 | 1.2E-03 |
| Exon length | 5'UTR length | 0.170 | 1.8E-01 | 0.310 | 8.4E-02 | 0.334 | 1.9E-01 |
| Exon length | 3'UTR length | 0.088 | 4.9E-01 | 0.260 | 1.5E-01 | 0.312 | 2.2E-01 |
| Controlled maximum lifespan | Weight | -0.152 | 1.7E-01 | -0.397 | 5.7E-03 | -0.191 | 4.5E-01 |
| Controlled maximum lifespan | Temperature | 0.036 | 7.5E-01 | -0.411 | 4.1E-03 | 0.223 | 3.7E-01 |
| Controlled maximum lifespan | Relative intron length | 0.178 | 1.1E-01 | -0.077 | 6.1E-01 | 0.056 | 8.3E-01 |
| Controlled maximum lifespan | mRNA length | 0.077 | 5.0E-01 | -0.060 | 6.9E-01 | 0.021 | 9.4E-01 |
| Controlled maximum lifespan | Mass adjusted BMR | 0.467 | 2.1E-05 | 0.351 | 1.6E-02 | 0.484 | 4.9E-02 |
| Controlled maximum lifespan | Intron length between first and last exons | 0.182 | 1.1E-01 | -0.124 | 4.0E-01 | 0.007 | 9.8E-01 |
| Controlled maximum lifespan | Intron length between first and last coding exons | 0.158 | 1.6E-01 | -0.124 | 4.1E-01 | -0.009 | 9.7E-01 |
| Controlled maximum lifespan | Genome size | 0.148 | 1.8E-01 | 0.025 | 8.7E-01 | -0.165 | 5.1E-01 |
| Controlled maximum lifespan | GC3 | -0.064 | 5.7E-01 | -0.096 | 5.2E-01 | -0.210 | 4.0E-01 |
| Controlled maximum lifespan | GC contents | -0.029 | 8.0E-01 | -0.022 | 8.8E-01 | -0.212 | 4.0E-01 |
| Controlled maximum lifespan | Exon+Intron length between first and last exon | 0.183 | 1.0E-01 | -0.059 | 6.9E-01 | 0.023 | 9.3E-01 |
| Controlled maximum lifespan | Exon+Intron length between first and last coding exon | 0.162 | 1.5E-01 | -0.053 | 7.3E-01 | 0.011 | 9.6E-01 |
| Controlled maximum lifespan | Exon number | 0.187 | 9.2E-02 | 0.036 | 8.1E-01 | 0.086 | 7.4E-01 |
| Controlled maximum lifespan | Exon length | -0.296 | 6.9E-03 | -0.188 | 2.1E-01 | -0.168 | 5.1E-01 |
| Controlled maximum lifespan | Controlled intron length | -0.028 | 8.1E-01 | -0.081 | 5.9E-01 | 0.402 | 9.9E-02 |
| Controlled maximum lifespan | CDS length | 0.076 | 5.0E-01 | -0.058 | 7.0E-01 | -0.066 | 7.9E-01 |
| Controlled maximum lifespan | CAI | 0.240 | 3.0E-02 | 0.107 | 4.7E-01 | 0.240 | 3.4E-01 |
| Controlled maximum lifespan | Basal metabolic rate | -0.395 | 4.2E-04 | -0.410 | 4.5E-03 | -0.338 | 1.8E-01 |
| Controlled maximum lifespan | 5'UTR length | 0.159 | 2.1E-01 | 0.075 | 6.9E-01 | 0.039 | 8.8E-01 |
| Controlled maximum lifespan | 3'UTR length | 0.170 | 1.8E-01 | -0.069 | 7.1E-01 | 0.093 | 7.2E-01 |
| Controlled intron length | mRNA length | 0.459 | 1.6E-05 | 0.791 | 3.7E-11 | 0.414 | 8.9E-02 |
| Controlled intron length | Intron length between first and last exons | 0.083 | 4.6E-01 | 0.842 | 0.0E+0<br>0 | 0.393 | 1.1E-01 |
| Controlled intron length | GC3 | -0.235 | 3.4E-02 | 0.139 | 3.5E-01 | -0.302 | 2.2E-01 |
| Controlled intron length | GC contents | -0.257 | 2.0E-02 | 0.201 | 1.8E-01 | -0.321 | 1.9E-01 |
| Controlled intron length | Exon+Intron length between first and last exon | 0.198 | 7.6E-02 | 0.853 | 0.0E+0<br>0 | 0.430 | 7.6E-02 |
| Controlled intron length | Exon+Intron length between first and last coding exon | 0.173 | 1.2E-01 | 0.869 | 0.0E+0<br>0 | 0.439 | 7.0E-02 |
| Controlled intron length | Exon number | 0.296 | 6.9E-03 | 0.483 | 5.9E-04 | 0.499 | 3.5E-02 |
| Controlled intron length | Exon length | 0.513 | 8.1E-07 | 0.808 | 6.4E-12 | 0.186 | 4.6E-01 |
| Controlled intron length | CDS length | 0.512 | 8.9E-07 | 0.915 | 2.7E-19 | 0.578 | 1.2E-02 |
| Controlled intron length | CAI | 0.136 | 2.2E-01 | 0.061 | 6.8E-01 | 0.313 | 2.1E-01 |
| Controlled intron length | 5'UTR length | 0.035 | 7.8E-01 | 0.074 | 6.9E-01 | 0.426 | 8.9E-02 |
| Controlled intron length | 3'UTR length | 0.084 | 5.1E-01 | 0.053 | 7.7E-01 | 0.453 | 6.9E-02 |
| CDS length | 5'UTR length | 0.407 | 8.5E-04 | 0.107 | 5.6E-01 | 0.511 | 3.6E-02 |
| CDS length | 3'UTR length | 0.373 | 2.4E-03 | 0.140 | 4.5E-01 | 0.451 | 6.9E-02 |
| CAI | Exon number | -0.301 | 6.1E-03 | -0.336 | 2.1E-02 | -0.293 | 2.4E-01 |
| CAI | Exon length | -0.019 | 8.7E-01 | 0.063 | 6.7E-01 | 0.166 | 5.1E-01 |
| CAI | CDS length | -0.118 | 2.9E-01 | -0.010 | 9.5E-01 | 0.235 | 3.5E-01 |
| CAI | 5'UTR length | 0.009 | 9.4E-01 | 0.279 | 1.2E-01 | 0.235 | 3.6E-01 |
| CAI | 3'UTR length | 0.171 | 1.8E-01 | 0.249 | 1.7E-01 | 0.152 | 5.6E-01 |
| Basal metabolic rate | Weight | 0.958 | 6.8E-42 | 0.997 | 6.4E-52 | 0.920 | 1.7E-07 |
| Basal metabolic rate | Temperature | 0.401 | 3.4E-04 | 0.444 | 1.8E-03 | -0.221 | 3.9E-01 |
| Basal metabolic rate | Relative intron length | 0.489 | 9.6E-06 | 0.386 | 7.7E-03 | -0.184 | 4.8E-01 |
| Basal metabolic rate | mRNA length | -0.092 | 4.3E-01 | 0.230 | 1.2E-01 | -0.098 | 7.1E-01 |
| Basal metabolic rate | Mass adjusted BMR | -0.709 | 0.0E+0<br>0 | -0.970 | 0.0E+0<br>0 | -0.811 | 1.0E-04 |
| Basal metabolic rate | Intron length between first and last exons | 0.428 | 1.5E-04 | 0.347 | 1.7E-02 | -0.191 | 4.6E-01 |
| Basal metabolic rate | Intron length between first and last coding exons | 0.462 | 3.3E-05 | 0.355 | 1.5E-02 | -0.186 | 4.7E-01 |
| Basal metabolic rate | Genome size | 0.356 | 1.7E-03 | -0.082 | 5.9E-01 | 0.100 | 7.0E-01 |
| Basal metabolic rate | GC3 | -0.097 | 4.0E-01 | 0.079 | 6.0E-01 | 0.005 | 9.9E-01 |
| Basal metabolic rate | GC contents | -0.024 | 8.4E-01 | 0.090 | 5.5E-01 | -0.010 | 9.7E-01 |
| Basal metabolic rate | Exon+Intron length between first and last exon | 0.378 | 8.9E-04 | 0.346 | 1.8E-02 | -0.154 | 5.5E-01 |
| Basal metabolic rate | Exon+Intron length between first and last coding exon | 0.452 | 5.0E-05 | 0.394 | 6.5E-03 | -0.137 | 6.0E-01 |
| Basal metabolic rate | Exon number | -0.121 | 3.0E-01 | 0.190 | 2.0E-01 | -0.031 | 9.1E-01 |
| Basal metabolic rate | Exon length | 0.013 | 9.1E-01 | 0.177 | 2.3E-01 | -0.058 | 8.3E-01 |
| Basal metabolic rate | Controlled intron length | -0.108 | 3.5E-01 | 0.316 | 3.1E-02 | -0.279 | 2.8E-01 |
| Basal metabolic rate | CDS length | -0.075 | 5.2E-01 | 0.294 | 4.5E-02 | -0.037 | 8.9E-01 |
| Basal metabolic rate | CAI | 0.164 | 1.6E-01 | -0.006 | 9.7E-01 | -0.103 | 6.9E-01 |
| Basal metabolic rate | 5'UTR length | -0.170 | 2.0E-01 | -0.127 | 4.9E-01 | -0.129 | 6.3E-01 |
| Basal metabolic rate | 3'UTR length | 0.044 | 7.4E-01 | 0.043 | 8.2E-01 | 0.015 | 9.6E-01 |
| 5'UTR length | 3'UTR length | 0.887 | 1.6E-22 | 0.913 | 3.4E-13 | 0.956 | 0.0E+0<br>0 |

**Figure S9.**
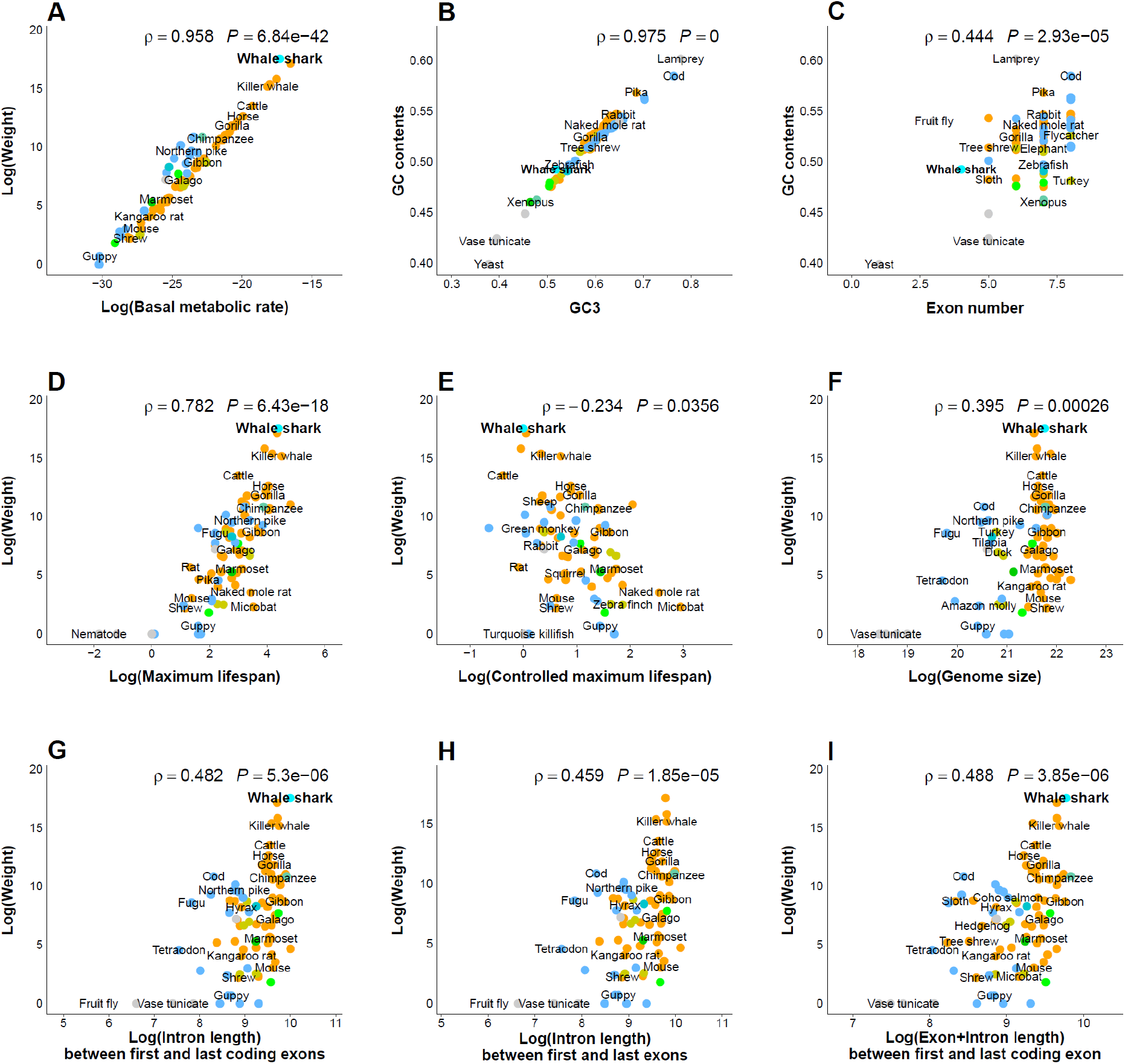
Scaling relationships between genomic and physiologic properties across 82 species. Extended data from Figure 2. The properties on both x-axis and y-axis were used to calculate Spearman’s rank correlation coefficient for each plot. All *p*-values and rho values are shown in top of each plot. The general species names are centered over their dots. Overlapping species names in the same layer were not plotted. The nine colors of dots indicate biological classification (gray: Hyperoartia, Ascidiacea, Chromadorea, Insecta and Saccharomycetes, turquoise: Chondrichthyes (cyan: whale shark), light blue: Actinopterygii, aquamarine: Sarcopterygii, dark green: Amphibia, light green: Reptilia, dark yellow: Aves, orange: Mammalia).

### 3.3 Intron gain or loss

From the single-copy orthologous gene sets, CDS of each orthologous family were aligned using MUSCLE (version 3.8.31)^41^. Exon-exon boundaries as intron positions were marked on the CDS alignments. The intron positions within a permissible length (six bp), which are considered as alternative splice sites, were aligned. The aligned intron position was converted to a binary character matrix as an input table of the Malin program^42^, which was used to calculate the “intron gain or loss” using Dollo parsimony (Figure S10).

**Figure S10.**
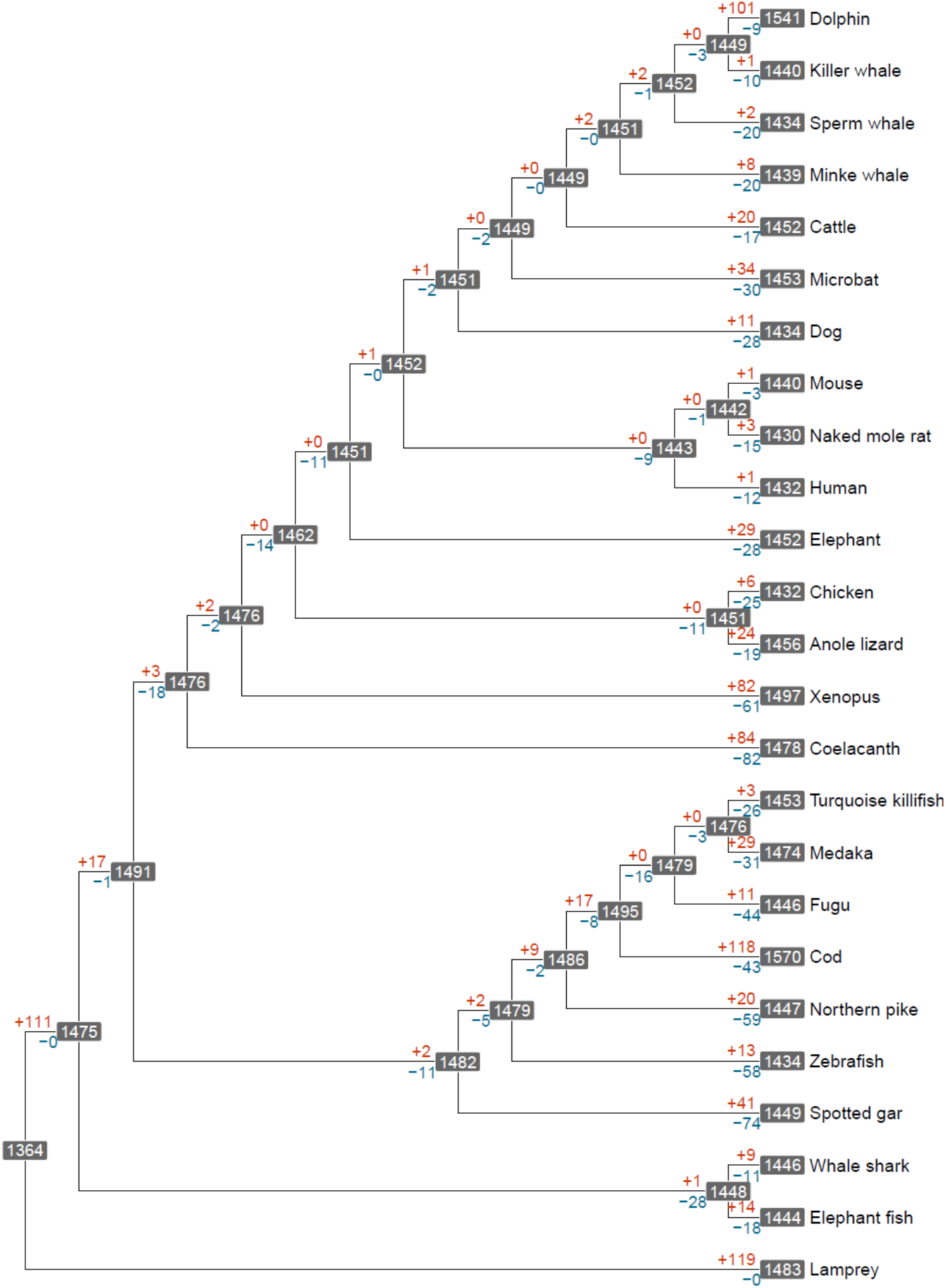
Intron gain or loss in the single-copy orthologous gene group. The phylogenetic tree was derived from Figure 3D. The intron gains and losses were computed by Dollo parsimony using Malin^42^. The numbers in the gray boxes indicate the number of introns. Red and blue numbers indicate the number of gained and lost introns from the most common ancestor, respectively.

### 3.4 Prediction of repetitive elements within introns

To compare intronic repetitive elements in each species, we constructed consensus models of putative interspersed repeats by RepeatModeler (version 1.0.10)^7^. Using RepeatMasker (version 4.0.5)^6^, we then predicted repeat elements in the introns of 81 species (yeast is excluded from our 82 species set) with the ‘-no_is-cutoff 255-frag 20000’ options. The predicted repetitive elements containing domains that overlapped with other repeats (higher-scoring match), which were denoted by asterisk in the RepeatMasker result file, were filtered out.

**Figure S11.**
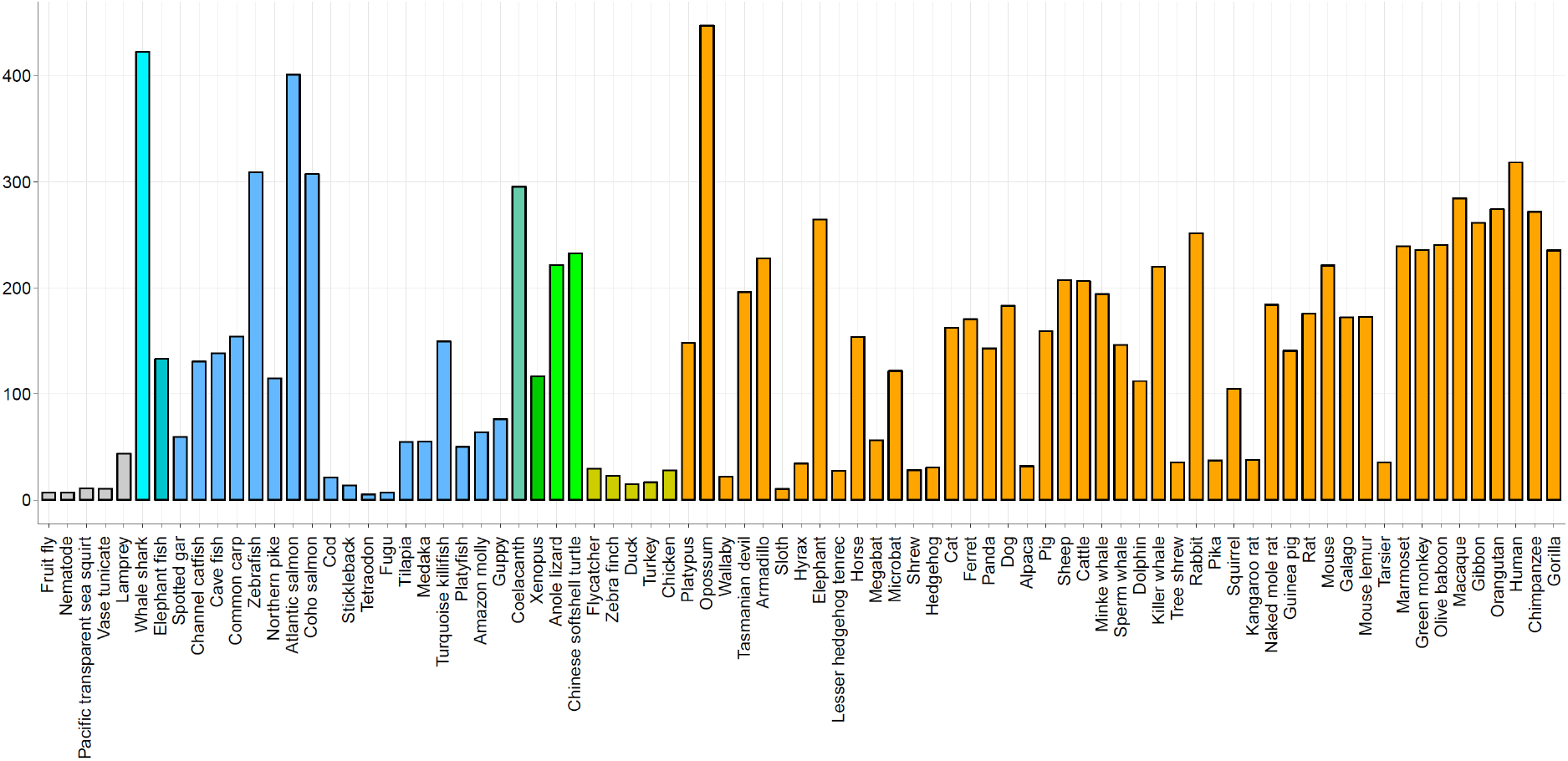
Total length of repetitive elements in the introns of 81 species. The total length was summed across ten repetitive elements: SINEs, LINEs, LTR elements, DNA elements, unclassified elements, interspersed repeats, small RNA, satellites, simple repeats, and low complexity region. Colors indicate the species class as in Figure 1. Yeast is excluded from 82 species.

**Figure S12.**
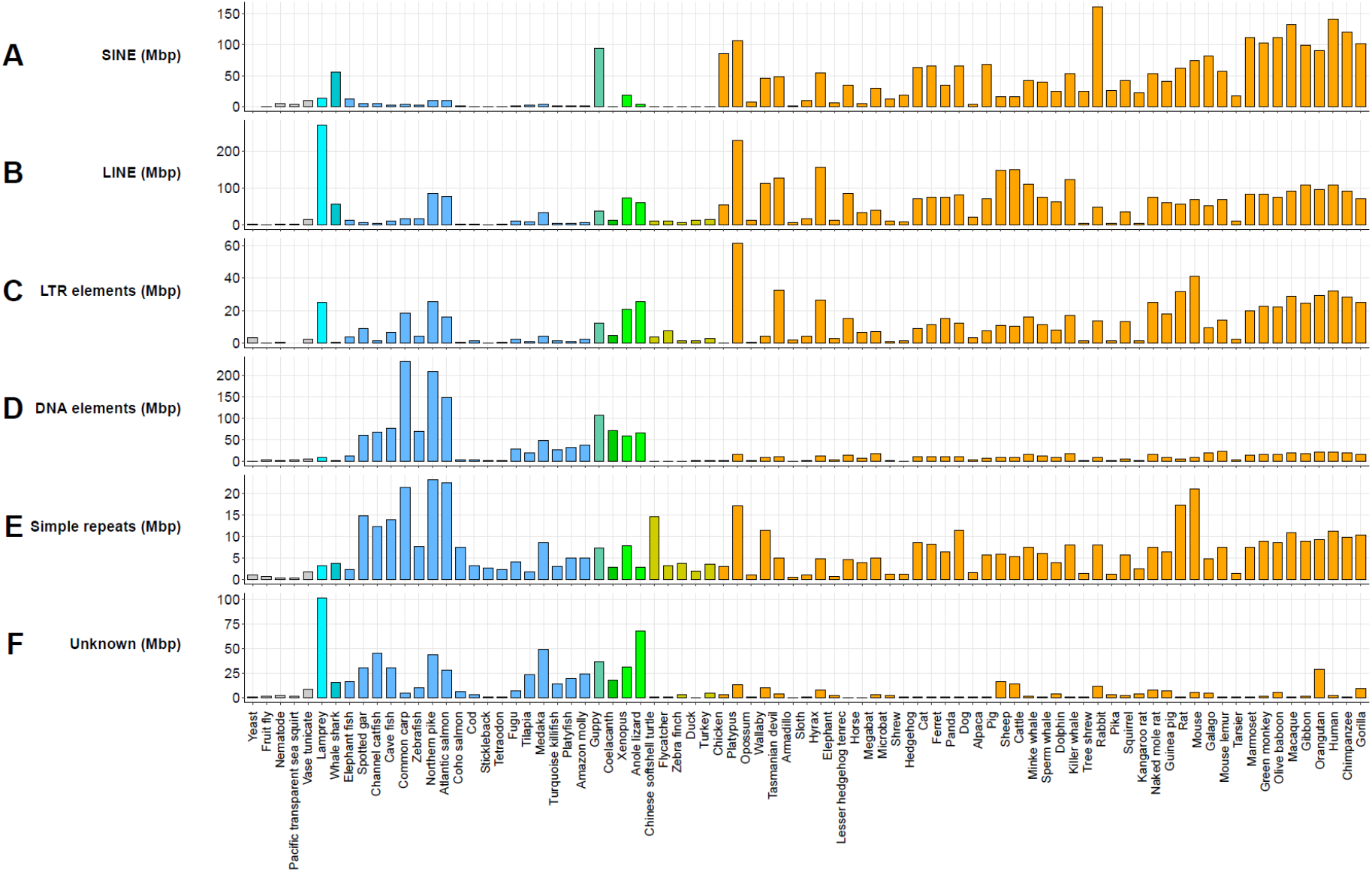
Total length of six repetitive elements in the introns of 81 species. The bold text to the left of the plots indicates the type of repeat element. Colors indicate the species class as in Figure 1. Yeast is excluded from 82 species.

**Figure S13.**
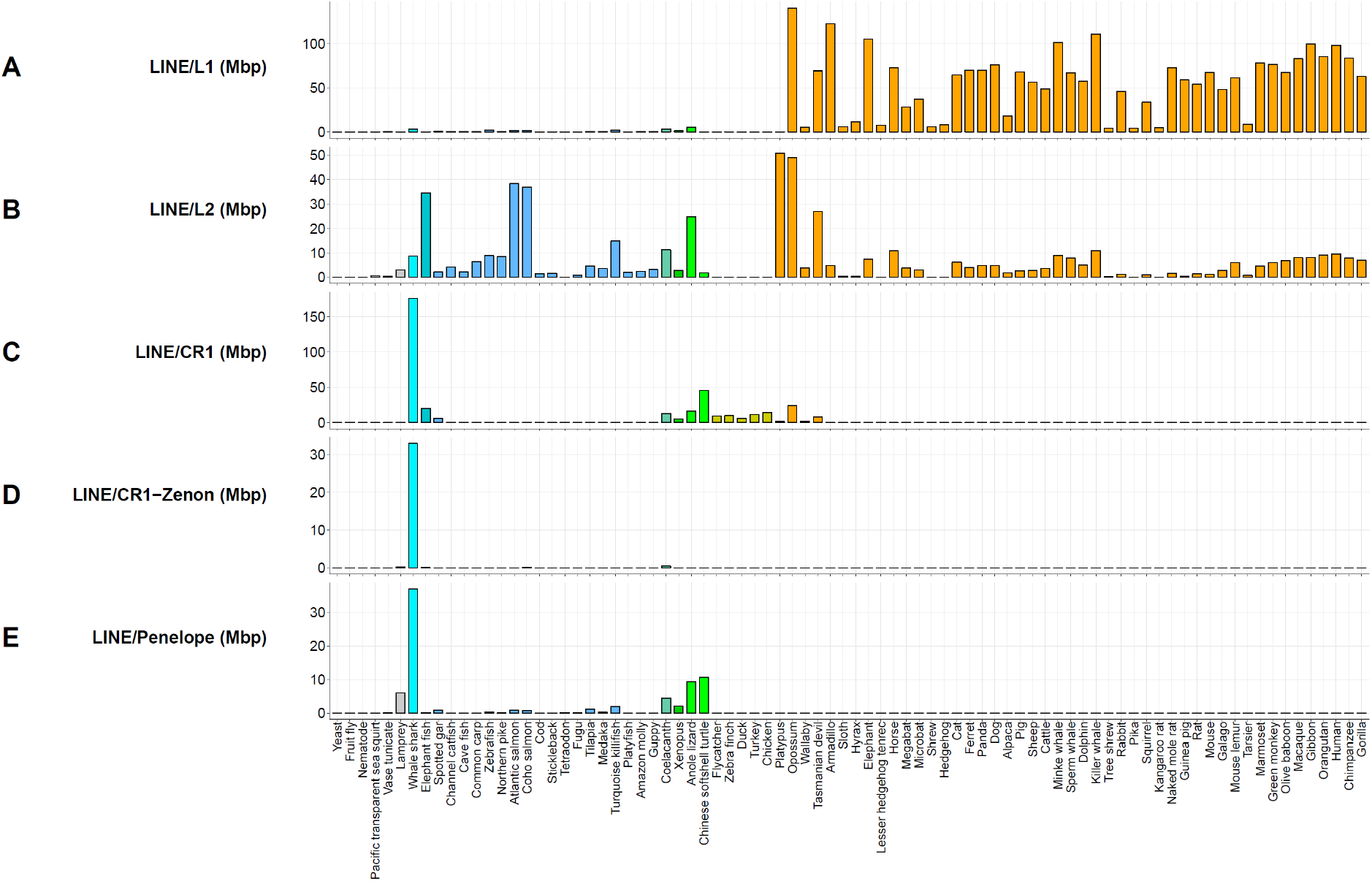
Total length of five LINEs in the introns of 81 species. The bold text to the left of the plots indicates the type of repeat elements. Colors indicate the species class as in Figure 1. Yeast is excluded from 82 species.

**Figure S14.**
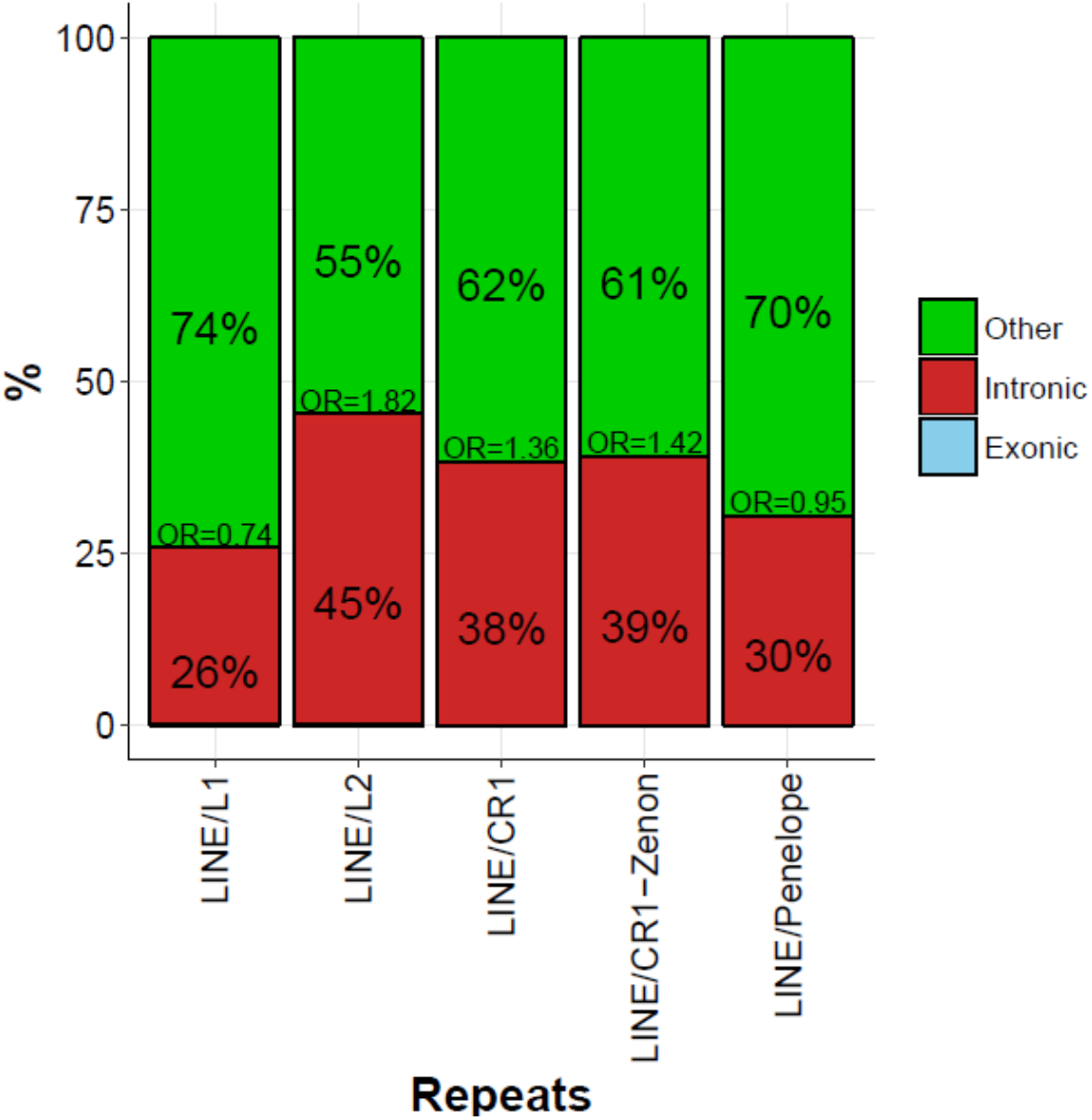
Distribution of LINEs in the whale shark genome. Using the non-redundant predicted gene model, we analyzed the proportion of LINE elements across the intronic, exonic, and intergenic regions. The bar plots show the percentage of five repetitive elements in each of the three regions. The actual percentages are shown in middle of the bars. Odds ratio analyses of the five LINE elements across exonic, intronic, and intergenic regions within the whale shark showed a slightly higher representation of these elements within introns, though these findings lacked statistical significance (*p*-value calculated by chi-square test > 0.05, Figure S14). The odd ratios are listed in the middle of the bars, and are the ratio of the proportion of repetitive elements in the intronic region to the proportion of the intronic region in the whale shark genome.

**Figure S15.**
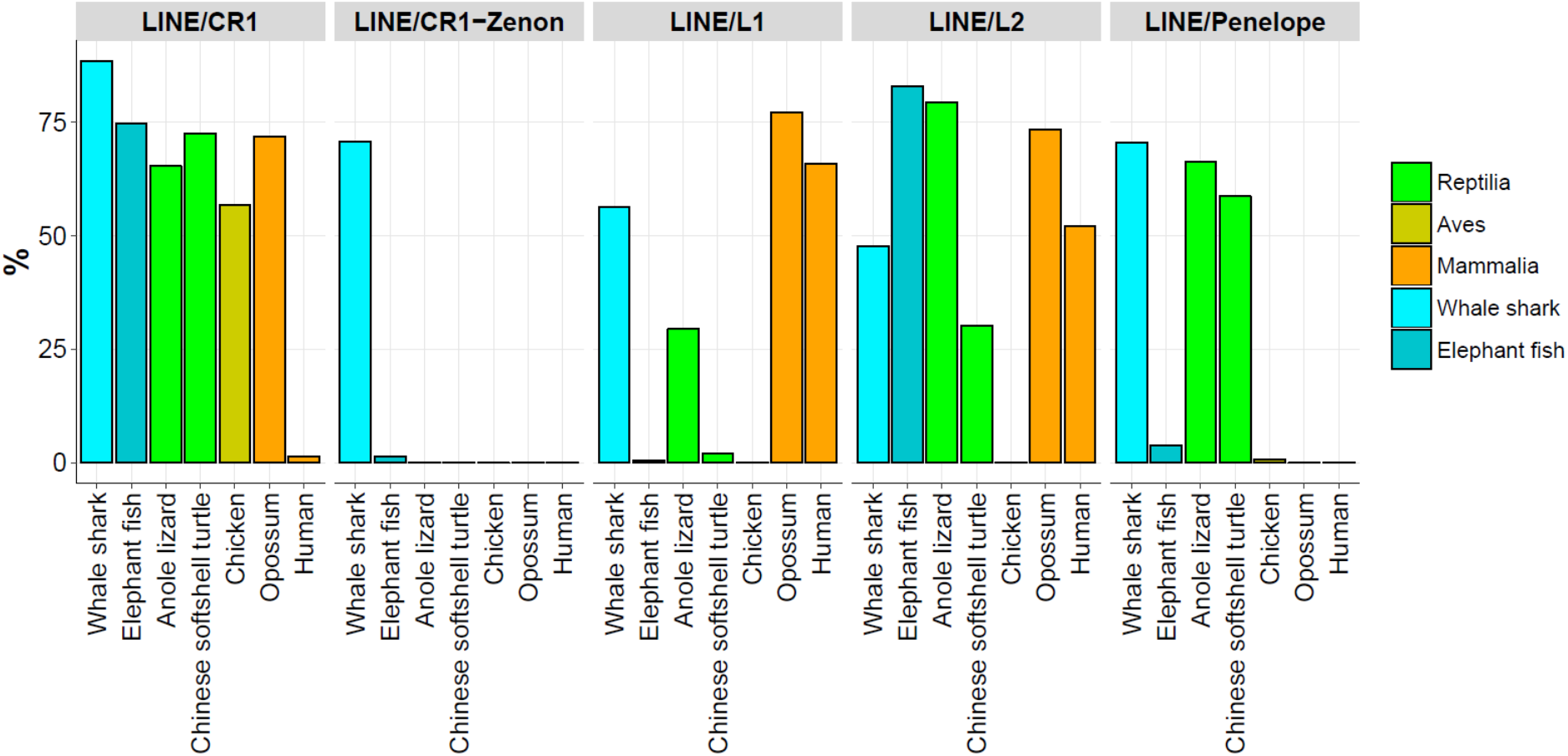
Proportion of genes containing LINEs in their introns. Each bar plot shows the proportion of genes which have repetitive elements (element types are shown in gray boxes) in the genome of each species (x-axis).

### 3.5 Synonymous codon usage comparison

We measured relative synonymous codon usage (RSCU) using Sharp *et. al.*’s method^43^ in each of the 82 species. A principal component analysis (PCA) on the RSCU was performed using the R packages (version 3.3.0)^44^ ggplot2^45^ and ggfortify^46^ (Figure S16).

**Figure S16A.**
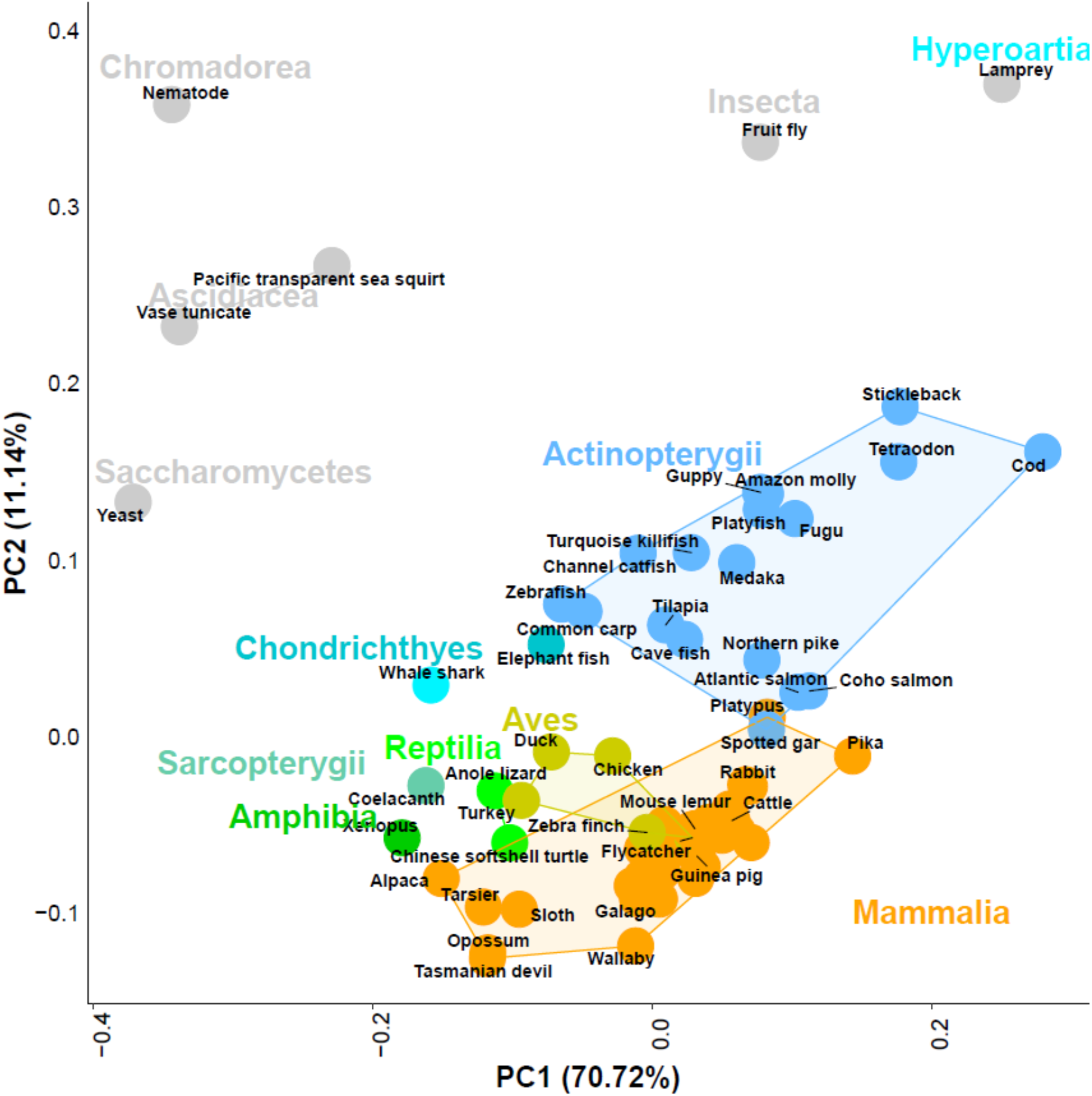
Principal component analysis of relative synonymous codon usage of 82 species. The common species name is shown in black text over a colored circle. Each color shows the class of 82 species (gray: Hyperoartia, Ascidiacea, Chromadorea, Insecta and Saccharomycetes, turquoise: Chondrichthyes (cyan: whale shark), light blue: Actinopterygii, aquamarine: Sarcopterygii, dark green: Amphibia, light green: Reptilia, dark yellow: Aves, orange: Mammalia).

**Figure S16B.**
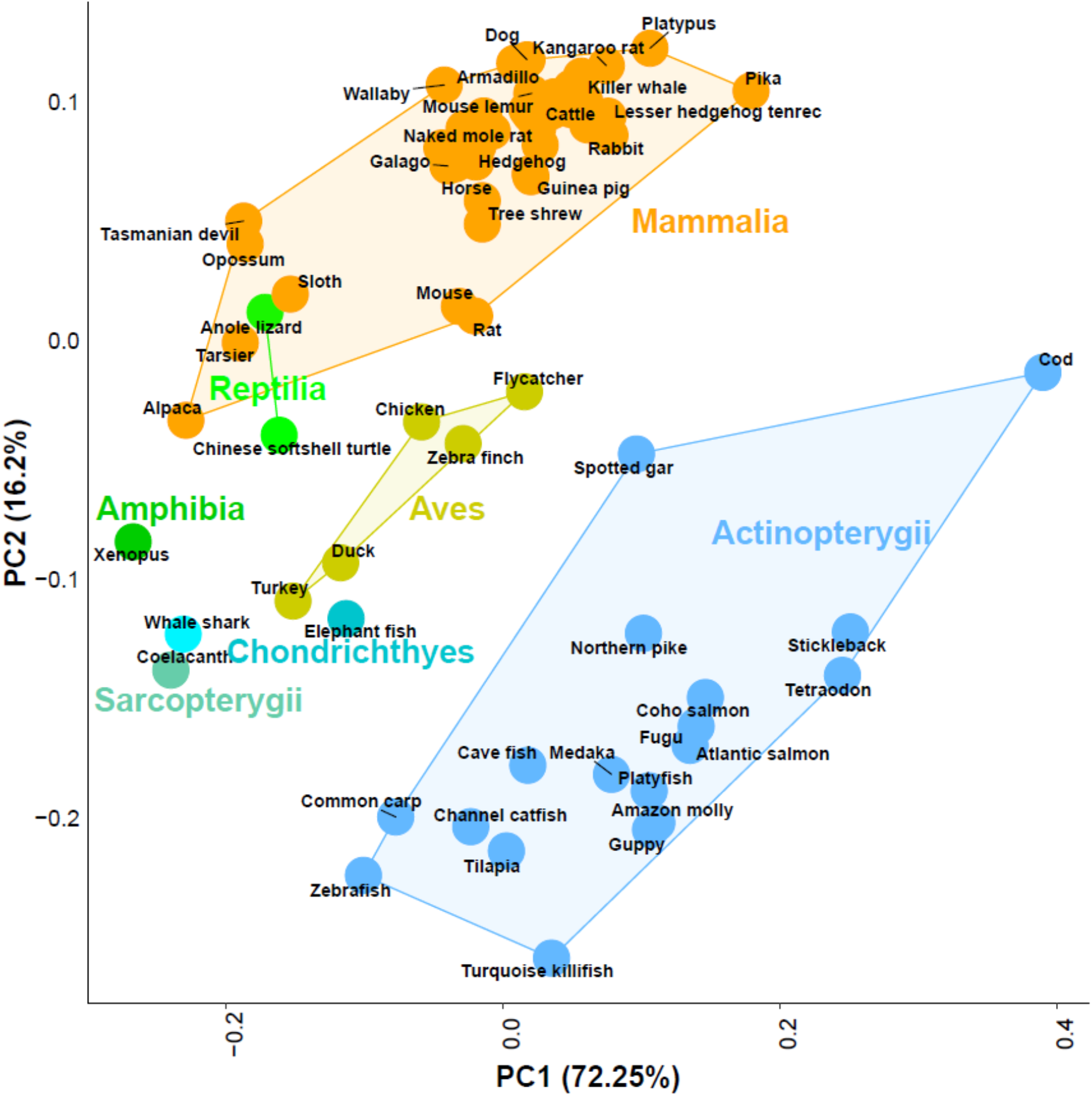
Principal component analysis of relative synonymous codon usage of 76 species. Six species having distant codon usage pattern were excluded from comparison of 82 species to investigate the evolutionary history of codon usage from whale shark to human. Common species name is shown in black text over a colored circle. Each color of circles shows the class of 76 species (turquoise: Chondrichthyes (cyan: whale shark), light blue: Actinopterygii, aquamarine: Sarcopterygii, dark green: Amphibia, light green: Reptilia, dark yellow: Aves, orange: Mammalia).

## 4. Evolutionary studies of whale shark

### 4.1 Phylogeny construction

We constructed a phylogenetic tree of 25 species including whale shark (*Anolis carolinensis, Balaenoptera acutorostrata scammoni, Bos taurus, Callorhinchus milii, Canis familiaris, Danio rerio, Esox lucius, Gadus morhua, Gallus gallus, Heterocephalus glaber, Homo sapiens, Latimeria chalumnae, Lepisosteus oculatus, Loxodonta africana, Mus musculus, Myotis lucifugus, Nothobranchius furzeri, Orcinus orca, Oryzias latipes, Petromyzon marinus, Physeter catodon, Rhincodon typus, Takifugu rubripes, Tursiops truncatus, Xenopus tropicalis*). We first extracted 275 single-copy gene families of 25 species from the orthologous gene family table of 82 species. We filtered out clusters with average GC3 below 0.45 to prevent bias^47^, leaving 255 clusters. We performed multiple sequence alignment (MSA) of each remaining single copy gene family using MUSCLE 3.8.31^41^ and concatenated the MSA results without gap regions. The phylogenetic tree was constructed using RAxML 8.2^48^ with maximum likelihood (1,000 bootstrapping), using the PROTCATLG amino acid substitution model (Figure 3D).

### 4.2 Divergence time estimation

We estimated that the common ancestor of the whale shark and elephant fish diverged roughly 268 million years ago (MYA) (Figure 3D). Divergence times were estimated using the MCMCtree program in PAML package 4.8^49^ with the independent rates model (clock=2). The date of the node between *O. orca-L. chalumnae* was constrained to 401-425 MYA and *O. latipes-R. typus* was constrained to 450-497 MYA based on the TimeTree database^50^.

### 4.3 Whale shark evolutionary rate

We compared the molecular evolutionary rate of the whale shark and other 23 species with sea lamprey as an outgroup. We found that the whale shark had the shortest distance to the outgroup (sea lamprey) indicating slowest evolutionary rate (Table S18). We also performed a relative rate test using MEGA7^51^, and found that the whale shark protein coding genes are evolving more slowly than any 81 species (Table S19). We also performed the Two-Cluster test with LINTRE^52^. The distances between nodes in the phylogenetic tree (Figure 3D and Figure S17) used as the pairwise distances for the Two-Cluster test. The distances from the sea lamprey as an outgroup were calculated using the ‘ape’ R-package^53^. The two-cluster test also supported that the whale shark has a slower evolutionary rate than the elephant fish (Table S20).

**Table S18.** Pairwise distance to the outgroup for 24 species

| Species | Distance to *Petromyzon marinus* |
| --- | --- |
| *Rhincodon typus* | **0.61505568** |
| *Callorhinchus milii* | 0.62512995 |
| *Latimeria chalumnae* | 0.63722456 |
| *Lepisosteus oculatus* | 0.6546978 |
| *Gallus gallus* | 0.66767913 |
| *Anolis carolinensis* | 0.68388984 |
| *Loxodonta africana* | 0.68996989 |
| *Canis familiaris* | 0.69043867 |
| *Homo sapiens* | 0.69124695 |
| *Balaenoptera acutorostrata scammoni* | 0.69823335 |
| *Orcinus orca* | 0.69905548 |
| *Tursiops truncatus* | 0.70085231 |
| *Physeter catodon* | 0.70232111 |
| *Bos taurus* | 0.70553639 |
| *Myotis lucifugus* | 0.70839722 |
| *Heterocephalus glaber* | 0.71537803 |
| *Esox lucius* | 0.71747889 |
| *Xenopus tropicalis* | 0.71869153 |
| *Mus musculus* | 0.72308379 |
| *Danio rerio* | 0.72473367 |
| *Nothobranchius furzeri* | 0.77434503 |
| *Gadus morhua* | 0.77479036 |
| *Takifugu rubripes* | 0.77698009 |
| *Oryzias latipes* | 0.78572188 |

The pairwise distances were calculated using the R-package ‘ape’^53^ with an outgroup (sea lamprey).

**Table S19.**
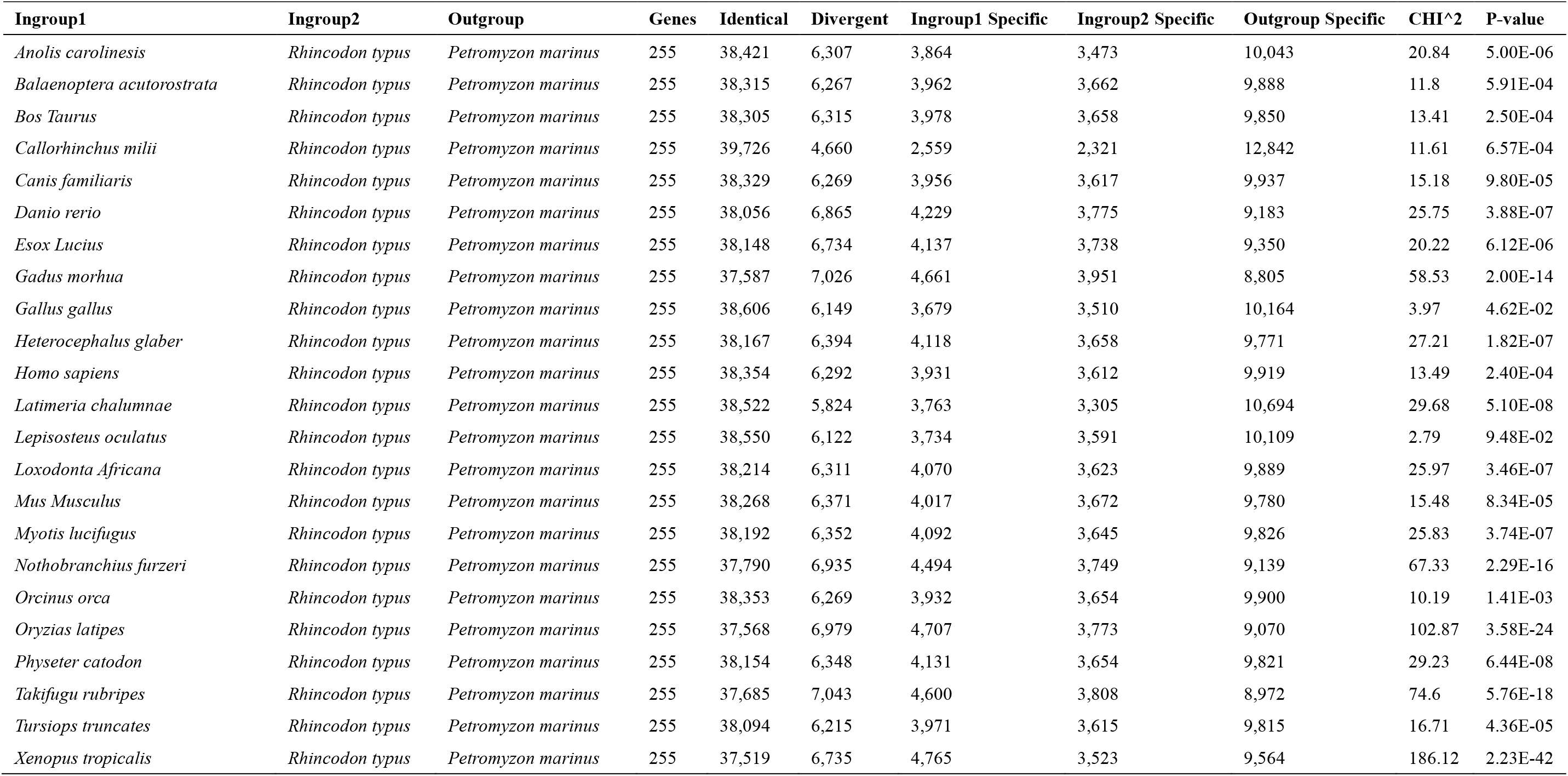
Results of relative rate test of whale shark versus other vertebrates. The ‘Identical’ and ‘Divergent’ columns indicate the number of sites where the amino acid is same or different in all three groups, respectively.

**Figure S17.**
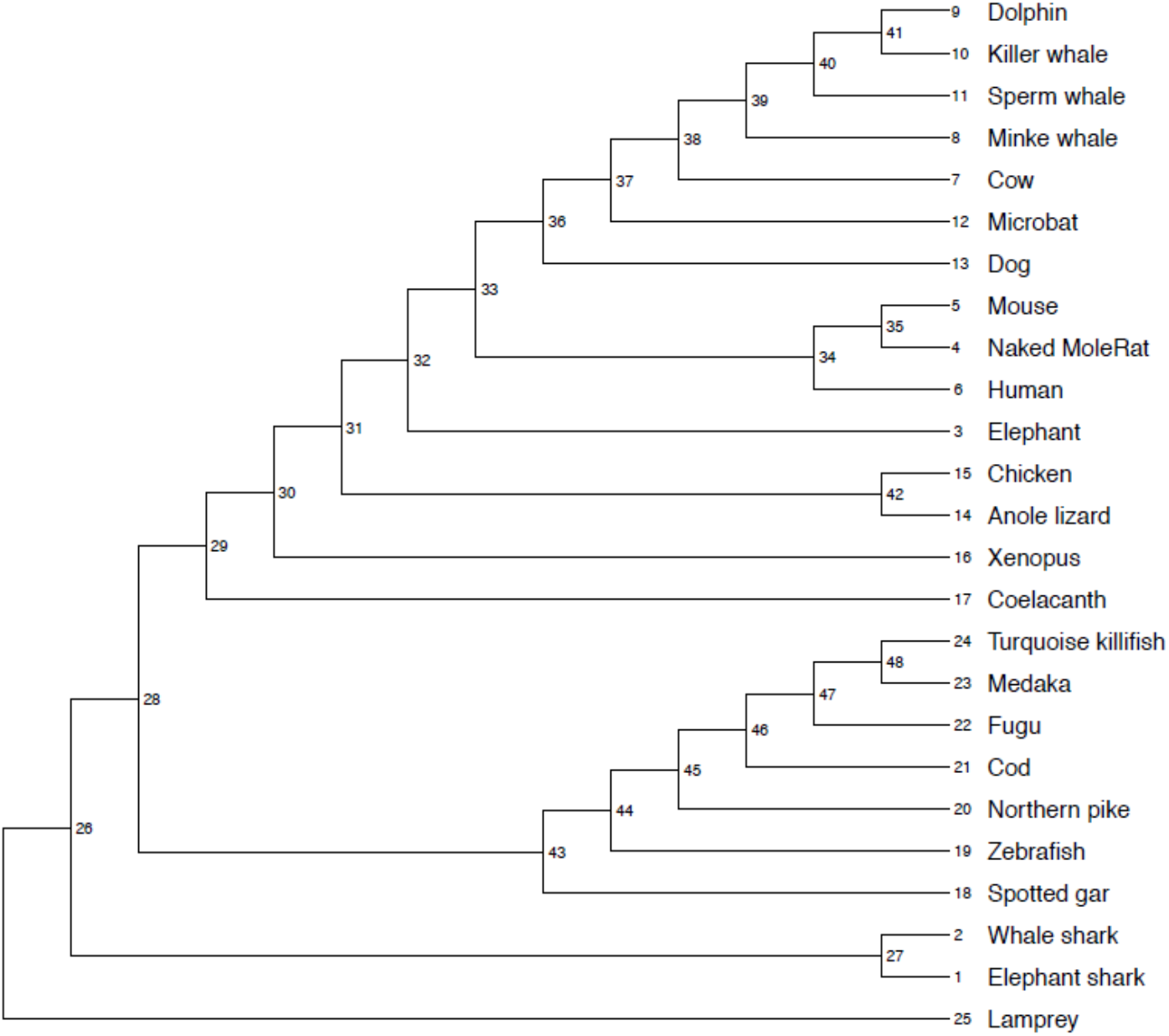
The phylogenetic tree used in the two-cluster test. Numbers indicate the nodes, the left, and the right in Table S20

**Table S20.** The results of two cluster test of whale shark versus other vertebrates

| Node | Left |  | Right | delta | s.e. | Z | CP | height | s.e. | bA | bB | bC |
| --- | --- | --- | --- | --- | --- | --- | --- | --- | --- | --- | --- | --- |
| 27 | 2 | < | 1 | 0.066988 | 0.001331 | 50.327722 | 99.96% | 0.104797 | 0.001132 | 0.071303 | 0.138291 | 0.128126 |
| 42 | 14 | < | 15 | 0.007774 | 0.000899 | 8.645688 | 99.96% | 0.041636 | 0.000634 | 0.037749 | 0.045523 | 0.1763 |
| 35 | 4 | > | 5 | 0.113236 | 0.00173 | 65.466742 | 99.96% | 0.148789 | 0.001454 | 0.205407 | 0.092171 | 0.106489 |
| 34 | 35 | < | 6 | 0.047336 | 0.001573 | 30.091571 | 99.96% | 0.161306 | 0.001385 | 0.137638 | 0.184974 | 0.11458 |
| 41 | 10 | < | 9 | 0.037975 | 0.0013 | 29.221827 | 99.96% | 0.100237 | 0.001098 | 0.08125 | 0.119225 | 0.13499 |
| 40 | 41 | > | 11 | 0.029928 | 0.000826 | 36.233157 | 99.96% | 0.063647 | 0.000669 | 0.078611 | 0.048683 | 0.161523 |
| 39 | 40 | < | 8 | 0.090038 | 0.0016 | 56.27053 | 99.96% | 0.171928 | 0.001515 | 0.126909 | 0.216947 | 0.097834 |
| 38 | 39 | < | 7 | 0.049123 | 0.001291 | 38.043271 | 99.96% | 0.140867 | 0.001195 | 0.116305 | 0.165428 | 0.129223 |
| 37 | 38 | < | 12 | 0.040654 | 0.001526 | 26.636184 | 99.96% | 0.142865 | 0.001167 | 0.122538 | 0.163191 | 0.131217 |
| 36 | 37 | < | 13 | 0.025798 | 0.001283 | 20.105648 | 99.96% | 0.129603 | 0.001044 | 0.116704 | 0.142502 | 0.1439 |
| 33 | 36 | < | 34 | 0.005831 | 0.000777 | 7.501552 | 99.96% | 0.131407 | 0.000932 | 0.128492 | 0.134322 | 0.136093 |
| 32 | 33 | > | 3 | 0.077327 | 0.000853 | 90.663726 | 99.96% | 0.112226 | 0.000887 | 0.150889 | 0.073562 | 0.118436 |
| 31 | 32 | > | 42 | 0.048181 | 0.000708 | 68.032487 | 99.96% | 0.106587 | 0.000812 | 0.130678 | 0.082497 | 0.139804 |
| 30 | 31 | > | 16 | 0.03557 | 0.000815 | 43.664202 | 99.96% | 0.096445 | 0.000766 | 0.11423 | 0.07866 | 0.15522 |
| 29 | 30 | < | 17 | 0.04978 | 0.001324 | 37.594456 | 99.96% | 0.147465 | 0.001199 | 0.122575 | 0.172355 | 0.141532 |
| 48 | 24 | < | 23 | 0.112731 | 0.001606 | 70.206746 | 99.96% | 0.169157 | 0.001603 | 0.112792 | 0.225523 | 0.081835 |
| 47 | 48 | < | 22 | 0.054553 | 0.001641 | 33.250362 | 99.96% | 0.156946 | 0.001331 | 0.129669 | 0.184222 | 0.118464 |
| 46 | 47 | > | 21 | 0.065644 | 0.000942 | 69.650583 | 99.96% | 0.110532 | 0.000931 | 0.143354 | 0.07771 | 0.125119 |
| 45 | 46 | < | 20 | 0.255255 | 0.003121 | 81.792064 | 99.96% | 0.246672 | 0.001959 | 0.119045 | 0.3743 | 0.120952 |
| 44 | 45 | < | 19 | 0.014926 | 0.001527 | 9.775209 | 99.96% | 0.169814 | 0.001295 | 0.162351 | 0.177277 | 0.126141 |
| 43 | 44 | > | 18 | 0.11091 | 0.000979 | 113.273282 | 99.96% | 0.123299 | 0.000928 | 0.178754 | 0.067844 | 0.11469 |
| 28 | 43 | > | 29 | 0.038478 | 0.000685 | 56.169011 | 99.96% | 0.137945 | 0.000922 | 0.157184 | 0.118706 | 0.128968 |
| 26 | 28 | > | 27 | 0.043569 | 0.001146 | 38.020739 | 99.96% | 0.115526 | 0.00083 | 0.13731 | 0.093741 | 0.180338 |

Q=17759.984030

**Figure S18.**
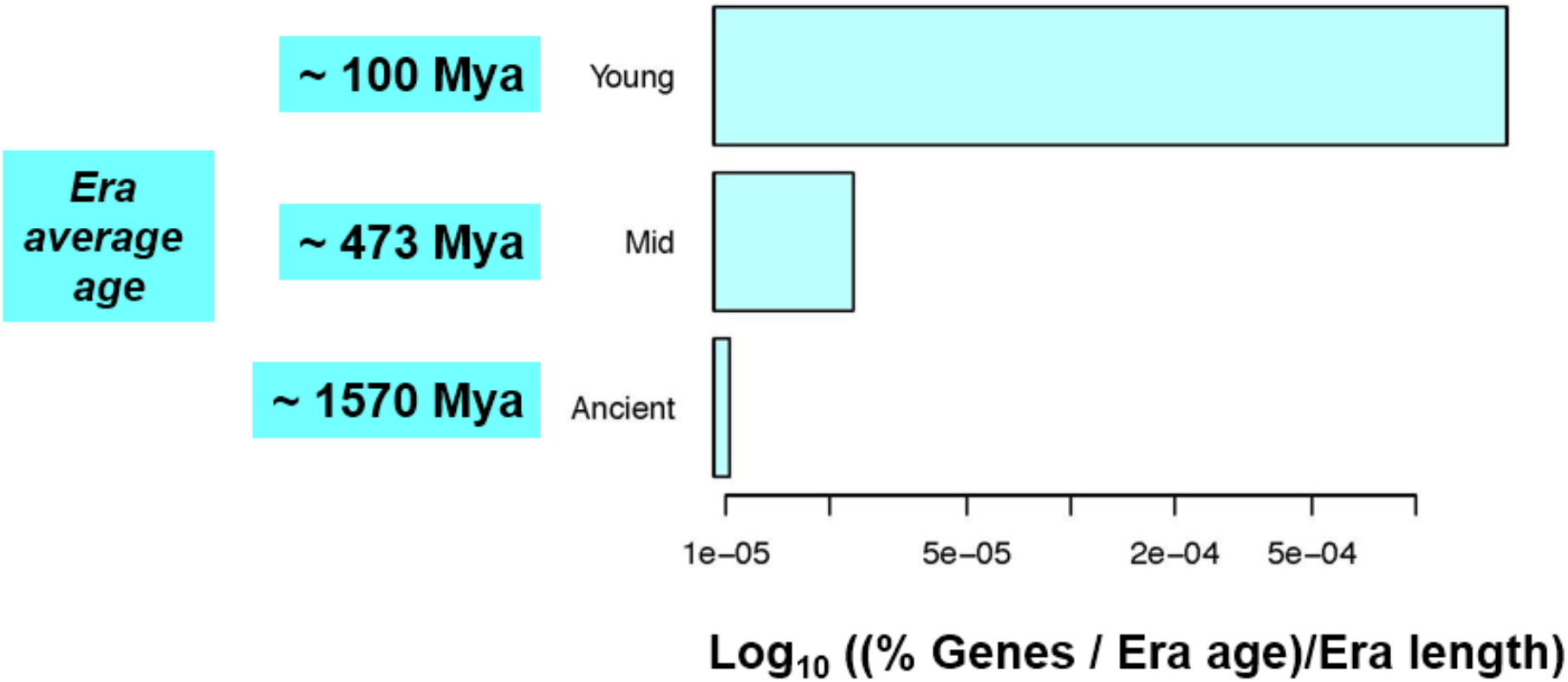
Supplementary figure linked to Figure 3C.

When accounting for the age and length (duration) of evolutionary eras (average era ages: ancient, ~1,570 Mya; mid, ~473 Mya; young, ~100 Mya), the number of genes in every era increases steadily as the genes are more recent, which suggests that gene turnover is highest in recent ages.

**Figure S19.**
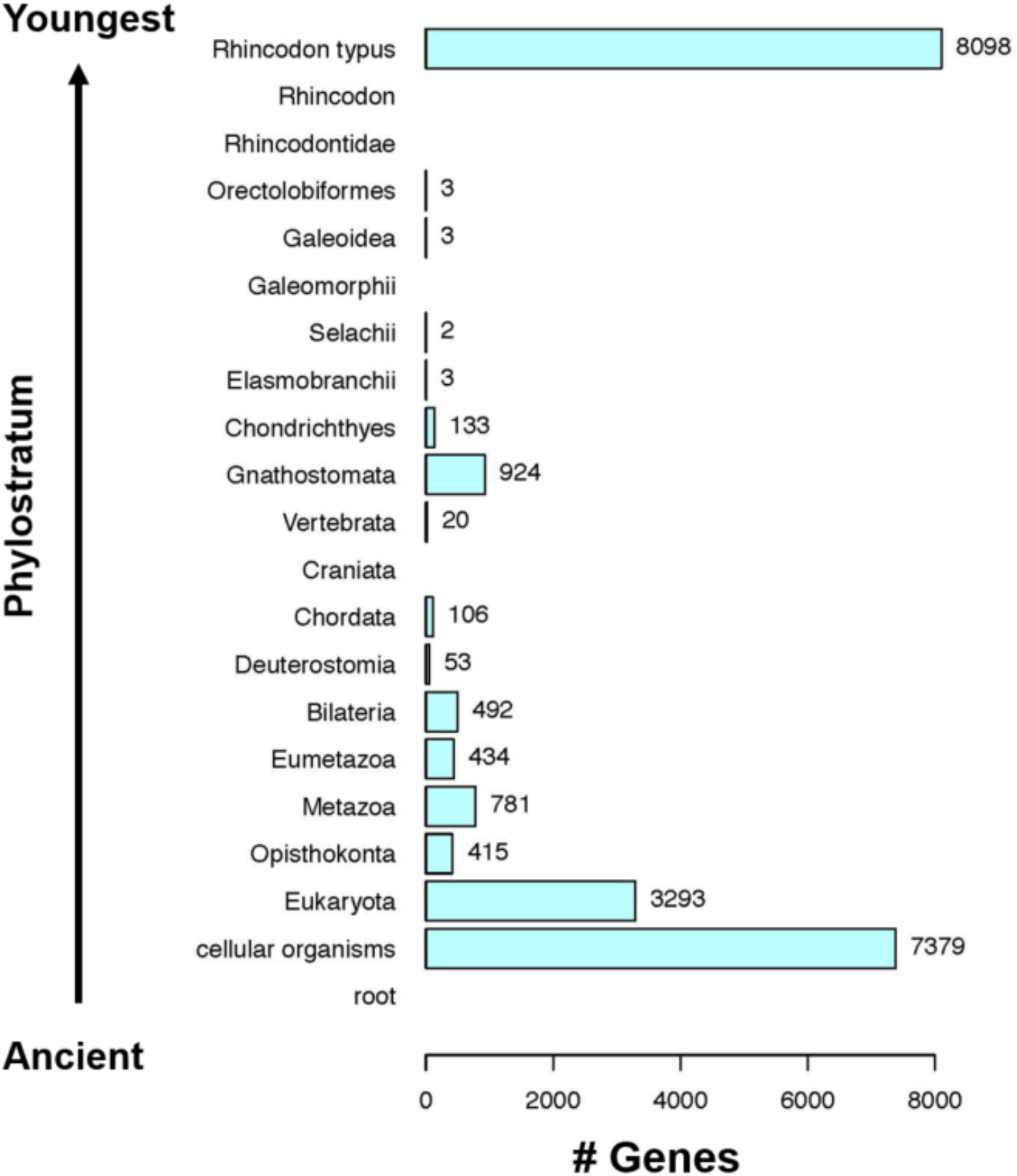
The number of genes in every phylostratum from most ancient to the youngest shows that most whale shark genes are ancient (7,379 genes in PS 1 and 3,293 in Eukaryota). The large number of genes that appear species-specific (8,098 genes) likely reflects the scarcity of sequenced genomes since the emergence of Chondrichthyes.

### 4.4 Gene family expansion and contraction analyses

Gene family expansion/contraction analyses were performed by using CAFÉ software^54^ (v3.1) with phylogenetic tree and p-value cut-off <0.05 demonstrating significantly changed the number of genes in the family (Figure 3D). 32 gene families were expanded and 233 gene families were contracted in the whale shark genome. We performed gene ontology (GO) enrichment test using ClueGO^55^. Expanded gene families were enriched in pattern specification involved in kidney development (GO:0061004) and nephron tubule formation (GO:0072079). Contracted gene families were enriched in nucleosome assembly (GO:0006334) and chromatin assembly (GO:0031497). We also found smaller number of histone 1 (H1), histone 2A (H2A) and histone 2Bs (H2Bs) in the whale shark than in other bony fishes and mammals (Figure S20).

**Figure S20.**
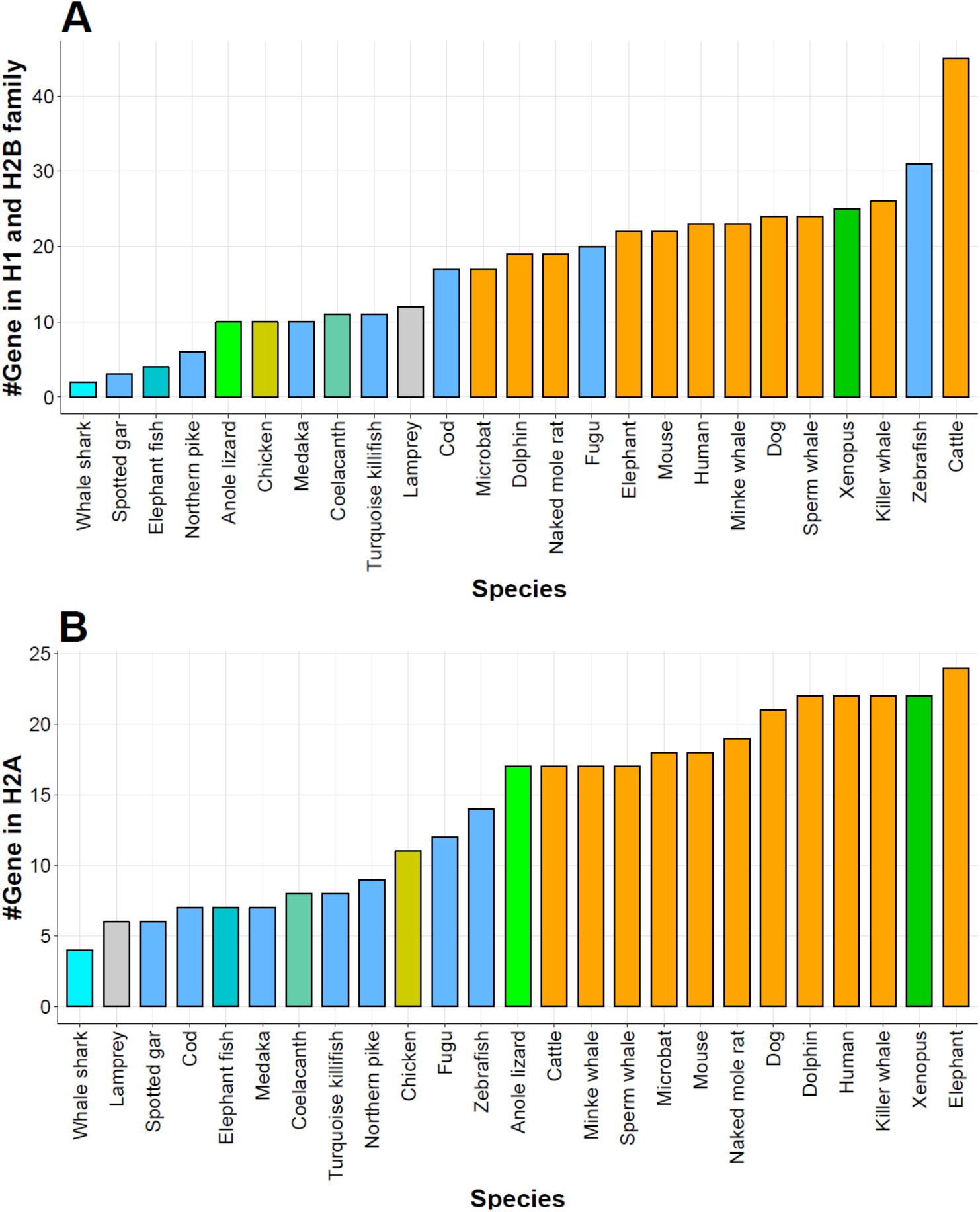
Contracted Histone gene families in whale shark. The OG000022 cluster contains the H1 and H2B classes. The OG000023 cluster contains the H2A class. The Y-axis indicates the number of histone genes in the cluster for each species. Each color shows the class of 82 species (gray: Hyperoartia, turquoise: Chondrichthyes (cyan: whale shark), light blue: Actinopterygii, aquamarine: Sarcopterygii, dark green: Amphibia, light green: Reptilia, dark yellow: Aves, orange: Mammalia).

**Table S21A.**
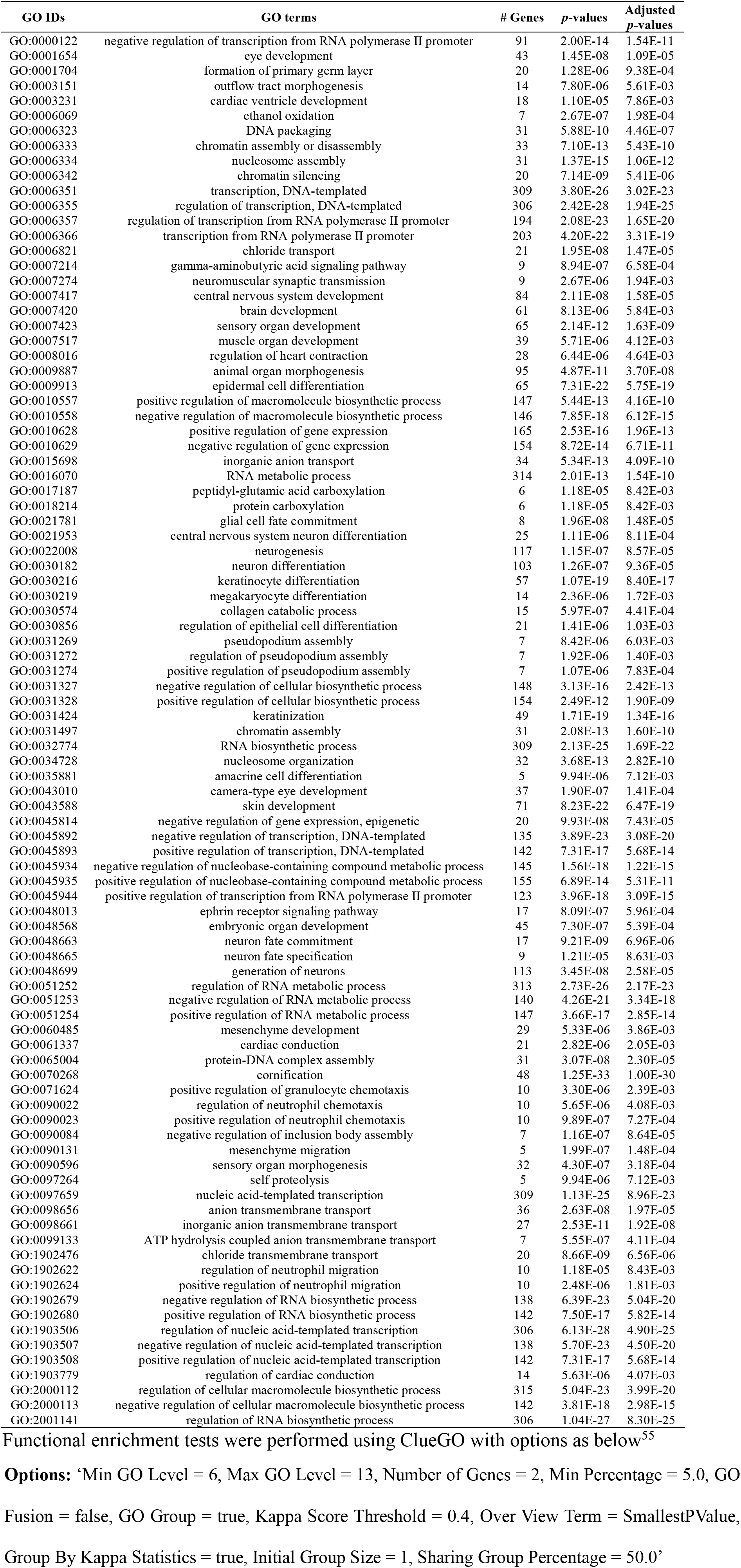
The GO terms enriched in contracted single-copy orthologous gene families in the whale shark from MRCA

| GO IDs | GO terms | # Genes | p-values | Adjusted p-values |
| --- | --- | --- | --- | --- |
| GO:0000122 | negative regulation of transcription from RNA polymerase II promoter | 91 | 2.00E-14 | 1.54E-11 |
| GO:0001654 | eye development | 43 | 1.45E-08 | 1.09E-05 |
| GO:0001704 | formation of primary germ layer | 20 | 1.28E-06 | 9.38E-04 |
| GO:0003151 | outflow tract morphogenesis | 14 | 7.80E-06 | 5.61E-03 |
| GO:0003231 | cardiac ventricle development | 18 | 1.10E-05 | 7.86E-03 |
| GO:0006069 | ethanol oxidation | 7 | 2.67E-07 | 1.98E-04 |
| GO:0006323 | DNA packaging | 31 | 5.88E-10 | 4.46E-07 |
| GO:0006333 | chromatin assembly or disassembly | 33 | 7.10E-13 | 5.43E-10 |
| GO:0006334 | nucleosome assembly | 31 | 1.37E-15 | 1.06E-12 |
| GO:0006342 | chromatin silencing | 20 | 7.14E-09 | 5.41E-06 |
| GO:0006351 | transcription, DNA-templated | 309 | 3.80E-26 | 3.02E-23 |
| GO:0006355 | regulation of transcription, DNA-templated | 306 | 2.42E-28 | 1.94E-25 |
| GO:0006357 | regulation of transcription from RNA polymerase II promoter | 194 | 2.08E-23 | 1.65E-20 |
| GO:0006366 | transcription from RNA polymerase II promoter | 203 | 4.20E-22 | 3.31E-19 |
| GO:0006821 | chloride transport | 21 | 1.95E-08 | 1.47E-05 |
| GO:0007214 | gamma-aminobutyric acid signaling pathway | 9 | 8.94E-07 | 6.58E-04 |
| GO:0007274 | neuromuscular synaptic transmission | 9 | 2.67E-06 | 1.94E-03 |
| GO:0007417 | central nervous system development | 84 | 2.11E-08 | 1.58E-05 |
| GO:0007420 | brain development | 61 | 8.13E-06 | 5.84E-03 |
| GO:0007423 | sensory organ development | 65 | 2.14E-12 | 1.63E-09 |
| GO:0007517 | muscle organ development | 39 | 5.71E-06 | 4.12E-03 |
| GO:0008016 | regulation of heart contraction | 28 | 6.44E-06 | 4.64E-03 |
| GO:0009887 | animal organ morphogenesis | 95 | 4.87E-11 | 3.70E-08 |
| GO:0009913 | epidermal cell differentiation | 65 | 7.31E-22 | 5.75E-19 |
| GO:0010557 | positive regulation of macromolecule biosynthetic process | 147 | 5.44E-13 | 4.16E-10 |
| GO:0010558 | negative regulation of macromolecule biosynthetic process | 146 | 7.85E-18 | 6.12E-15 |
| GO:0010628 | positive regulation of gene expression | 165 | 2.53E-16 | 1.96E-13 |
| GO:0010629 | negative regulation of gene expression | 154 | 8.72E-14 | 6.71E-11 |
| GO:0015698 | inorganic anion transport | 34 | 5.34E-13 | 4.09E-10 |
| GO:0016070 | RNA metabolic process | 314 | 2.01E-13 | 1.54E-10 |
| GO:0017187 | peptidyl-glutamic acid carboxylation | 6 | 1.18E-05 | 8.42E-03 |
| GO:0018214 | protein carboxylation | 6 | 1.18E-05 | 8.42E-03 |
| GO:0021781 | glial cell fate commitment | 8 | 1.96E-08 | 1.48E-05 |
| GO:0021953 | central nervous system neuron differentiation | 25 | 1.11E-06 | 8.11E-04 |
| GO:0022008 | neurogenesis | 117 | 1.15E-07 | 8.57E-05 |
| GO:0030182 | neuron differentiation | 103 | 1.26E-07 | 9.36E-05 |
| GO:0030216 | keratinocyte differentiation | 57 | 1.07E-19 | 8.40E-17 |
| GO:0030219 | megakaryocyte differentiation | 14 | 2.36E-06 | 1.72E-03 |
| GO:0030574 | collagen catabolic process | 15 | 5.97E-07 | 4.41E-04 |
| GO:0030856 | regulation of epithelial cell differentiation | 21 | 1.41E-06 | 1.03E-03 |
| GO:0031269 | pseudopodium assembly | 7 | 8.42E-06 | 6.03E-03 |
| GO:0031272 | regulation of pseudopodium assembly | 7 | 1.92E-06 | 1.40E-03 |
| GO:0031274 | positive regulation of pseudopodium assembly | 7 | 1.07E-06 | 7.83E-04 |
| GO:0031327 | negative regulation of cellular biosynthetic process | 148 | 3.13E-16 | 2.42E-13 |
| GO:0031328 | positive regulation of cellular biosynthetic process | 154 | 2.49E-12 | 1.90E-09 |
| GO:0031424 | keratinization | 49 | 1.71E-19 | 1.34E-16 |
| GO:0031497 | chromatin assembly | 31 | 2.08E-13 | 1.60E-10 |
| GO:0032774 | RNA biosynthetic process | 309 | 2.13E-25 | 1.69E-22 |
| GO:0034728 | nucleosome organization | 32 | 3.68E-13 | 2.82E-10 |
| GO:0035881 | amacrine cell differentiation | 5 | 9.94E-06 | 7.12E-03 |
| GO:0043010 | camera-type eye development | 37 | 1.90E-07 | 1.41E-04 |
| GO:0043588 | skin development | 71 | 8.23E-22 | 6.47E-19 |
| GO:0045814 | negative regulation of gene expression, epigenetic | 20 | 9.93E-08 | 7.43E-05 |
| GO:0045892 | negative regulation of transcription, DNA-templated | 135 | 3.89E-23 | 3.08E-20 |
| GO:0045893 | positive regulation of transcription, DNA-templated | 142 | 7.31E-17 | 5.68E-14 |
| GO:0045934 | negative regulation of nucleobase-containing compound metabolic process | 145 | 1.56E-18 | 1.22E-15 |
| GO:0045935 | positive regulation of nucleobase-containing compound metabolic process | 155 | 6.89E-14 | 5.31E-11 |
| GO:0045944 | positive regulation of transcription from RNA polymerase II promoter | 123 | 3.96E-18 | 3.09E-15 |
| GO:0048013 | ephrin receptor signaling pathway | 17 | 8.09E-07 | 5.96E-04 |
| GO:0048568 | embryonic organ development | 45 | 7.30E-07 | 5.39E-04 |
| GO:0048663 | neuron fate commitment | 17 | 9.21E-09 | 6.96E-06 |
| GO:0048665 | neuron fate specification | 9 | 1.21E-05 | 8.63E-03 |
| GO:0048699 | generation of neurons | 113 | 3.45E-08 | 2.58E-05 |
| GO:0051252 | regulation of RNA metabolic process | 313 | 2.73E-26 | 2.17E-23 |
| GO:0051253 | negative regulation of RNA metabolic process | 140 | 4.26E-21 | 3.34E-18 |
| GO:0051254 | positive regulation of RNA metabolic process | 147 | 3.66E-17 | 2.85E-14 |
| GO:0060485 | mesenchyme development | 29 | 5.33E-06 | 3.86E-03 |
| GO:0061337 | cardiac conduction | 21 | 2.82E-06 | 2.05E-03 |
| GO:0065004 | protein-DNA complex assembly | 31 | 3.07E-08 | 2.30E-05 |
| GO:0070268 | cornification | 48 | 1.25E-33 | 1.00E-30 |
| GO:0071624 | positive regulation of granulocyte chemotaxis | 10 | 3.30E-06 | 2.39E-03 |
| GO:0090022 | regulation of neutrophil chemotaxis | 10 | 5.65E-06 | 4.08E-03 |
| GO:0090023 | positive regulation of neutrophil chemotaxis | 10 | 9.89E-07 | 7.27E-04 |
| GO:0090084 | negative regulation of inclusion body assembly | 7 | 1.16E-07 | 8.64E-05 |
| GO:0090131 | mesenchyme migration | 5 | 1.99E-07 | 1.48E-04 |
| GO:0090596 | sensory organ morphogenesis | 32 | 4.30E-07 | 3.18E-04 |
| GO:0097264 | self proteolysis | 5 | 9.94E-06 | 7.12E-03 |
| GO:0097659 | nucleic acid-templated transcription | 309 | 1.13E-25 | 8.96E-23 |
| GO:0098656 | anion transmembrane transport | 36 | 2.63E-08 | 1.97E-05 |
| GO:0098661 | inorganic anion transmembrane transport | 27 | 2.53E-11 | 1.92E-08 |
| GO:0099133 | ATP hydrolysis coupled anion transmembrane transport | 7 | 5.55E-07 | 4.11E-04 |
| GO:1902476 | chloride transmembrane transport | 20 | 8.66E-09 | 6.56E-06 |
| GO:1902622 | regulation of neutrophil migration | 10 | 1.18E-05 | 8.43E-03 |
| GO:1902624 | positive regulation of neutrophil migration | 10 | 2.48E-06 | 1.81E-03 |
| GO:1902679 | negative regulation of RNA biosynthetic process | 138 | 6.39E-23 | 5.04E-20 |
| GO:1902680 | positive regulation of RNA biosynthetic process | 142 | 7.50E-17 | 5.82E-14 |
| GO:1903506 | regulation of nucleic acid-templated transcription | 306 | 6.13E-28 | 4.90E-25 |
| GO:1903507 | negative regulation of nucleic acid-templated transcription | 138 | 5.70E-23 | 4.50E-20 |
| GO:1903508 | positive regulation of nucleic acid-templated transcription | 142 | 7.31E-17 | 5.68E-14 |
| GO:1903779 | regulation of cardiac conduction | 14 | 5.63E-06 | 4.07E-03 |
| GO:2000112 | regulation of cellular macromolecule biosynthetic process | 315 | 5.04E-23 | 3.99E-20 |
| GO:2000113 | negative regulation of cellular macromolecule biosynthetic process | 142 | 3.81E-18 | 2.98E-15 |
| GO:2001141 | regulation of RNA biosynthetic process | 306 | 1.04E-27 | 8.30E-25 |

Functional enrichment tests were performed using ClueGO with options as below<sup>55</sup>
**Options:** ‘Min GO Level = 6, Max GO Level = 13, Number of Genes = 2, Min Percentage = 5.0, GO
Fusion = false, GO Group = true, Kappa Score Threshold = 0.4, Over View Term = SmallestPValue,
Group By Kappa Statistics = true, Initial Group Size = 1, Sharing Group Percentage = 50.0’

**Table S21B.**
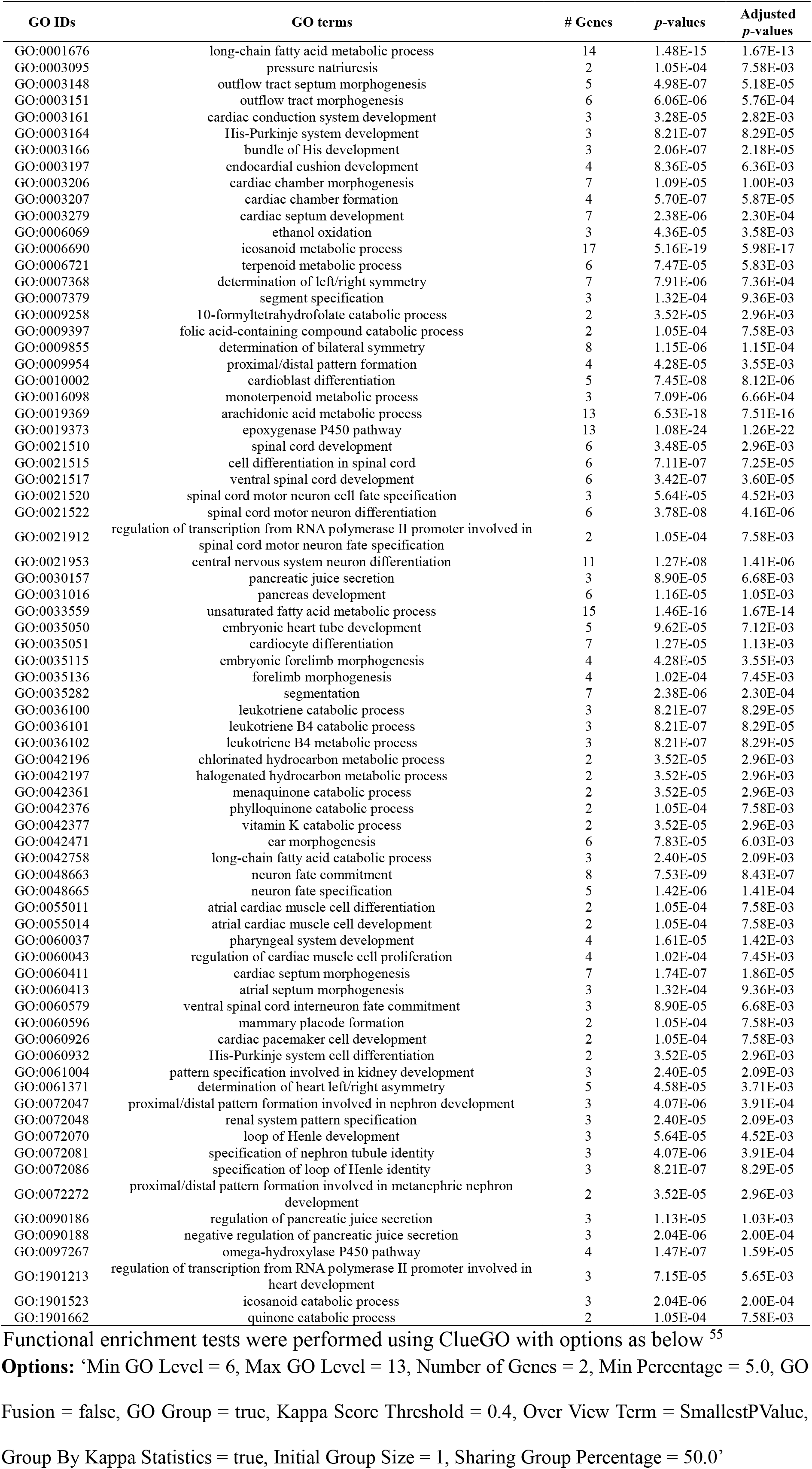
The GO terms enriched in expanded single-copy orthologous gene families in the whale shark from MRCA

### 4.5 Neural genes

We downloaded and corrected the neuronal genes with ten categories from GO and public databases as below.

1) Neuronal connectivity genes:

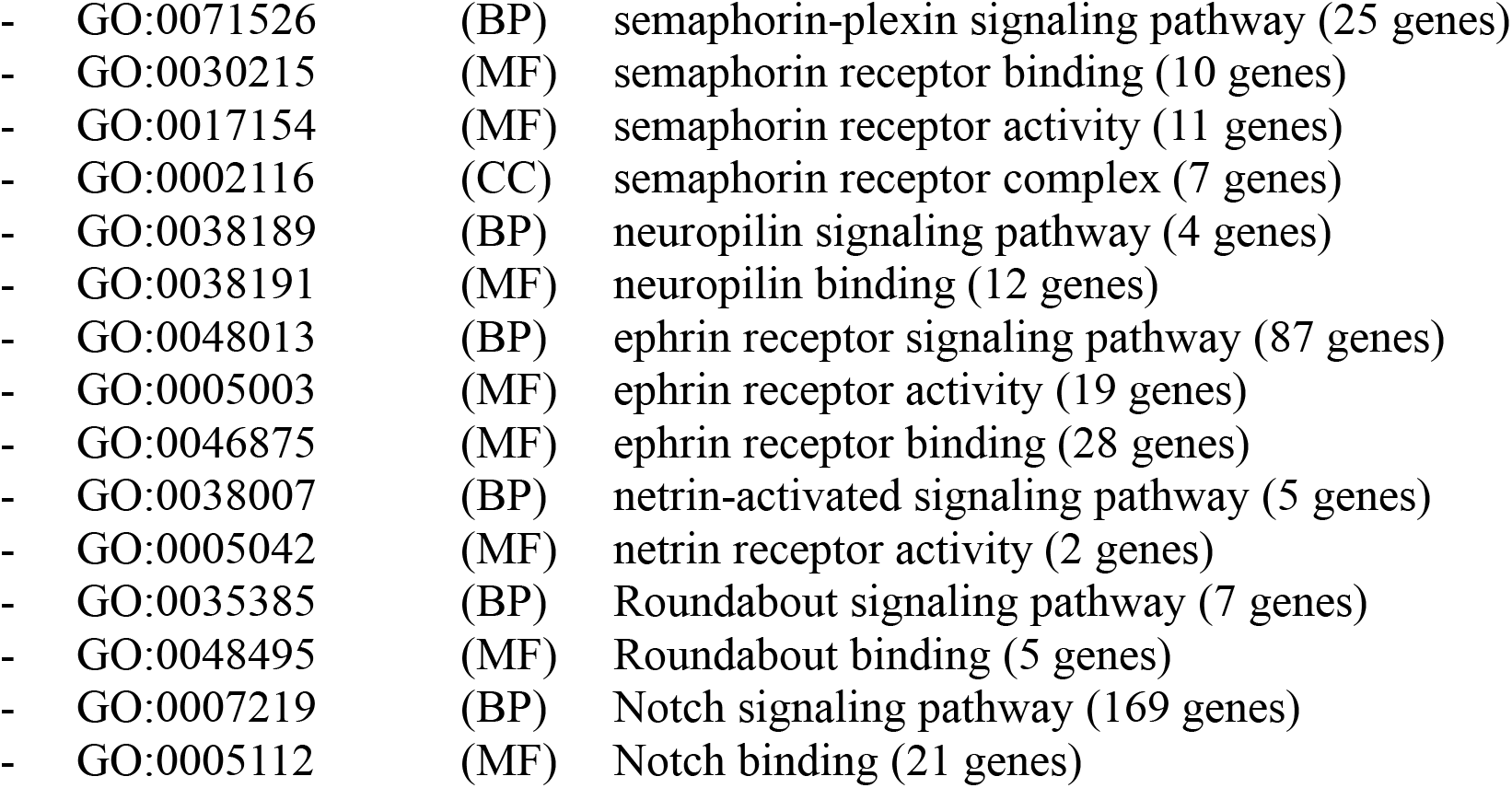
2) Cell adhesion:

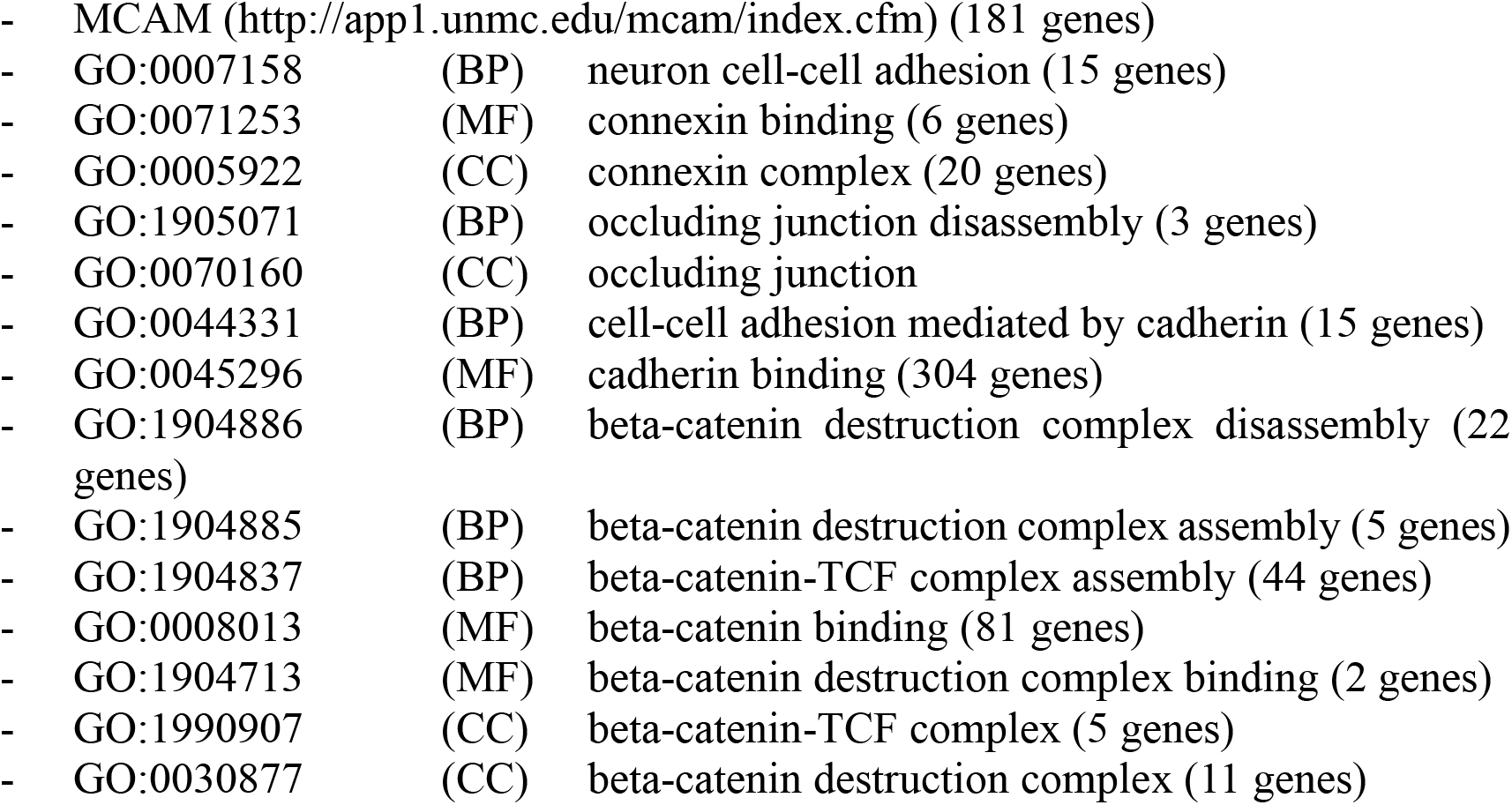
3) Olfactory receptors:

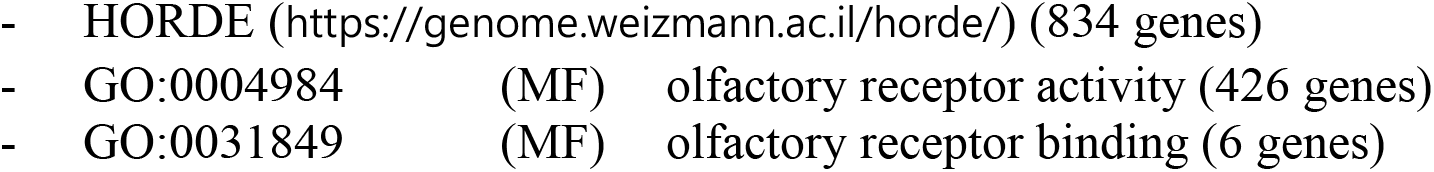
4) Ion channel:

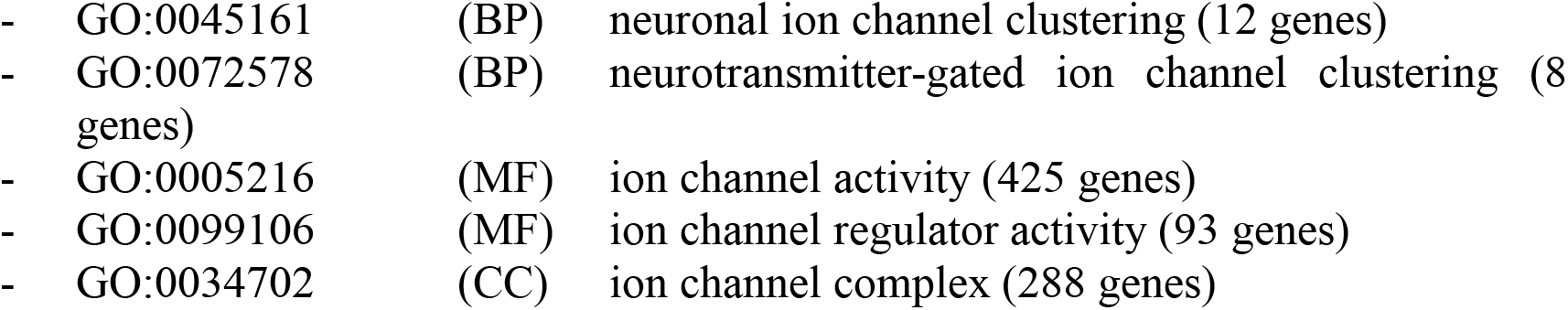
5) Unfolded protein response associated genes:

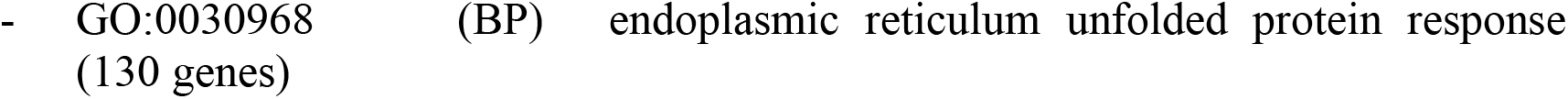
6) Neuronal activity and memory:

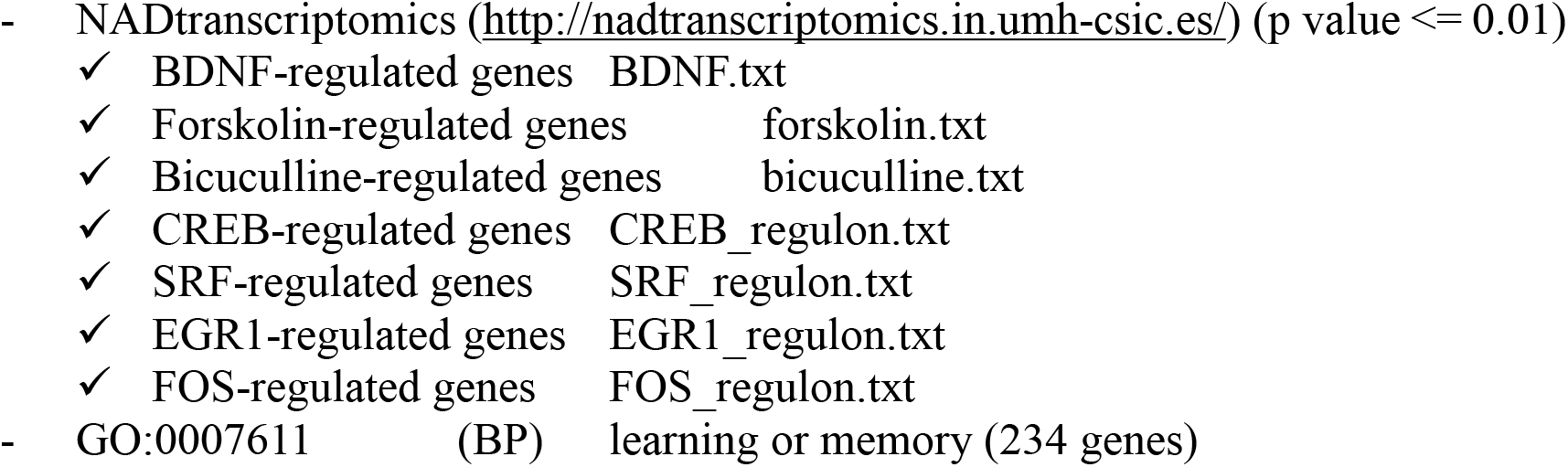
7) Neuropeptides:

- Neuropeptide database (http://www.neuropeptides.nl/tabel%20neuropeptides%20linked.htm) (96 genes)
- Two genes, CCAP and AstA (Allatostatin)
8) Homeobox genes:

- HGNC database (https://www.genenames.org/) (319 genes)
9) Synaptic genes:

- SynaptomeDB (http://metamoodics.org/SynaptomeDB/index.php) (1,886 genes)
10) Neurodegeneration:

- KEGG Human diseases (http://www.genome.jp/)
  ✓ Neurodegenerative diseases (236 genes)

**Figure S21.**
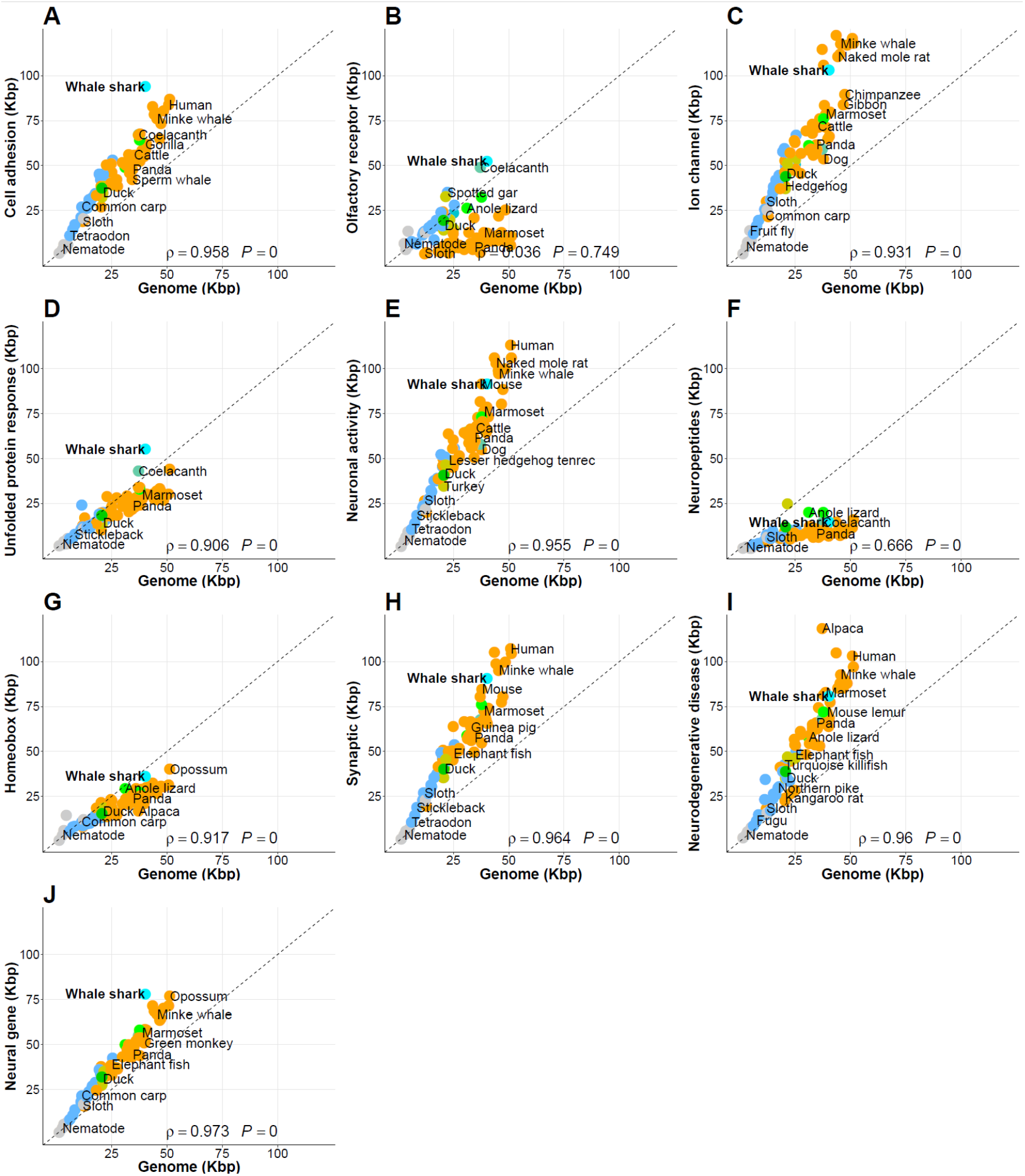
Supplementary figure linked to Figure 4 – All ten other scatter plots. Neuronal connectivity genes are longer in 81 species except yeast. The x- and y-axes correspond to average gene length (exon + intron) and the gene length of neuronal-related genes, respectively.

**Figure S22.**
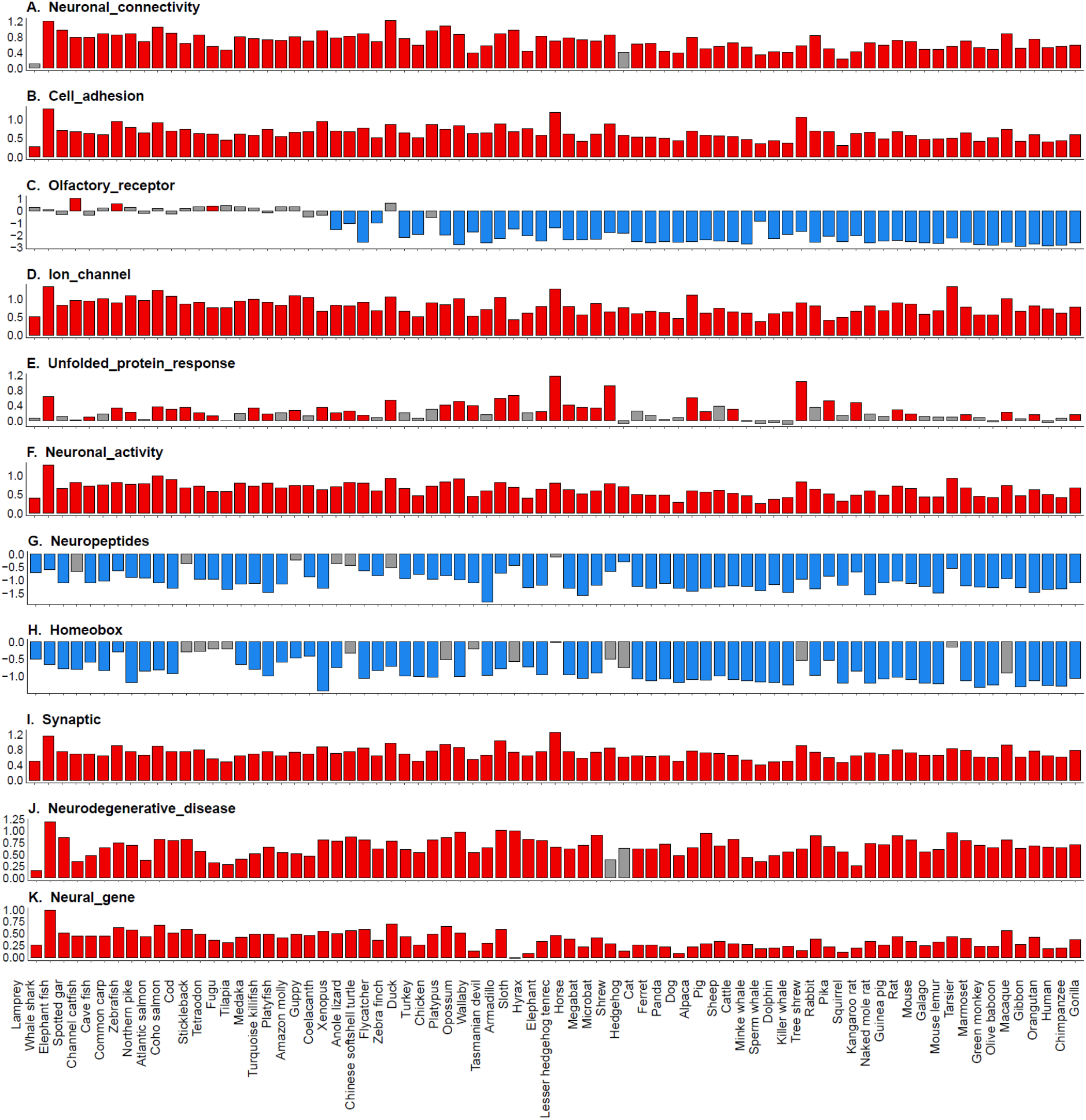
Relative median gene size of each neural subsets to median of gene size of genome. The Y-axis shows log transformed relative median value. The relative median values were calculated by dividing median of gene length (exon + intron) in each neuronal subset by median of gene length in genome. Red (or blue) bars indicate significantly higher (or lower) median gene length in the neuronal subset compared to the median genome-wide gene length by Wilcoxon-rank sum test.

### 4.6 Gene set enrichment analysis with gene size

Gene Set Enrichment Analysis (GSEA)^56^ was used to calculate statistically significant differences between short and long gene in 82 species using the clusterProfiler package^57^ with Gene Ontology. All genes were assigned to human gene symbols in order to use human-GO. Finally, we obtained the results of 77 species (Figure 4C and Additional File 1).

**Figure S23.**
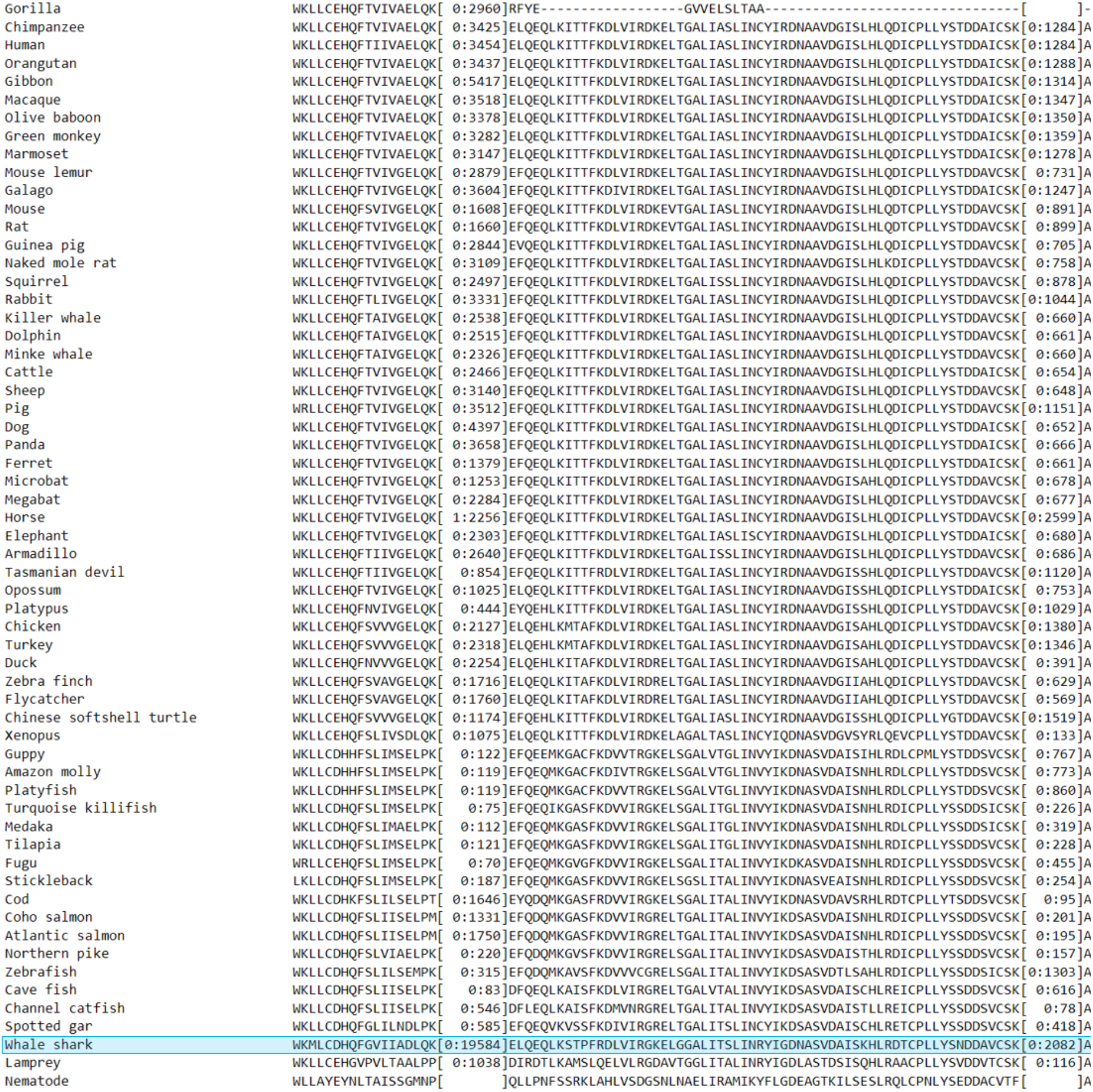
Portion of the sequence alignment of the NUP155 cluster of single copy orthologous genes. Intron position and length are shown in the square brackets. The cyan box shows the partially aligned sequence of the whale shark.

**Table S22.** GO enrichment of correlated single-copy orthologous gene families between gene length and the maximum lifespan, body weight, and BMR simultaneously

| GO ID | GO terms | # Genes | *p*-value | Adjusted *p*-value | Associated Genes Found |
| --- | --- | --- | --- | --- | --- |
| GO:0006405 | RNA export from nucleus | 7 | 2.E-04 | 6.E-03 | [ABCE1, ENY2, NUP155, RAE1, SARNP, SDAD1, XPO5] |
| GO:0006611 | protein export from nucleus | 9 | 3.E-05 | 8.E-04 | [ABCE1, CSE1L, ENY2, NUP155, RAE1, SARNP, SDAD1, STYX, XPO5] |
| GO:0007004 | telomere maintenance via telomerase | 3 | 2.E-02 | 2.E-02 | [MAPKAPK5, NAT10, XRCC5] |
| GO:0008209 | androgen metabolic process | 3 | 2.E-03 | 4.E-02 | [CYP19A1, HSD17B3, SGPL1] |
| GO:0008210 | estrogen metabolic process | 3 | 2.E-03 | 3.E-02 | [CYP19A1, SGPL1, STAR] |
| GO:0042537 | benzene-containing compound metabolic process | 3 | 1.E-03 | 2.E-02 | [AADAT, KMO, STAR] |
| GO:0043648 | dicarboxylic acid metabolic process | 5 | 3.E-03 | 5.E-02 | [AADAT, DLD, KMO, NMNAT2, STAR] |
| GO:0051168 | nuclear export | 9 | 6.E-05 | 2.E-03 | [ABCE1, CSE1L, ENY2, NUP155, RAE1, SARNP, SDAD1, STYX, XPO5] |
| GO:0051439 | regulation of ubiquitin-protein ligase activity involved in mitotic cell cycle | 3 | 2.E-02 | 5.E-02 | [ANAPC5, PSMA1, PSMD14] |
| GO:0060632 | regulation of microtubule-based movement | 3 | 1.E-03 | 3.E-02 | [CFAP20, CNIH4, TCTEX1D2] |
| GO:0061370 | testosterone biosynthetic process | 3 | 5.E-05 | 1.E-03 | [CYP19A1, HSD17B3, STAR] |
| GO:0065002 | intracellular protein transmembrane transport | 4 | 1.E-03 | 3.E-02 | [AGK, PEX3, PEX7, SRP54] |
| GO:0071166 | ribonucleoprotein complex localization | 6 | 9.E-04 | 2.E-02 | [ABCE1, ENY2, NUP155, RAE1, SARNP, SDAD1] |
| GO:0071426 | ribonucleoprotein complex export from nucleus | 6 | 8.E-04 | 2.E-02 | [ABCE1, ENY2, NUP155, RAE1, SARNP, SDAD1] |
| GO:0071806 | protein transmembrane transport | 4 | 2.E-03 | 4.E-02 | [AGK, PEX3, PEX7, SRP54] |

Functional enrichment tests were performed using ClueGO with options as below<sup>55</sup>
**Options:** ‘Min GO Level = 3, Max GO Level = 8, Number of Genes = 3, Min Percentage = 4.0, GO Fusion = false, GO Group = true, Kappa Score Threshold = 0.4, Over View Term = SmallestPValue, Group By Kappa Statistics = true, Initial Group Size = 1, Sharing Group Percentage = 50.0’

**Table S23.**
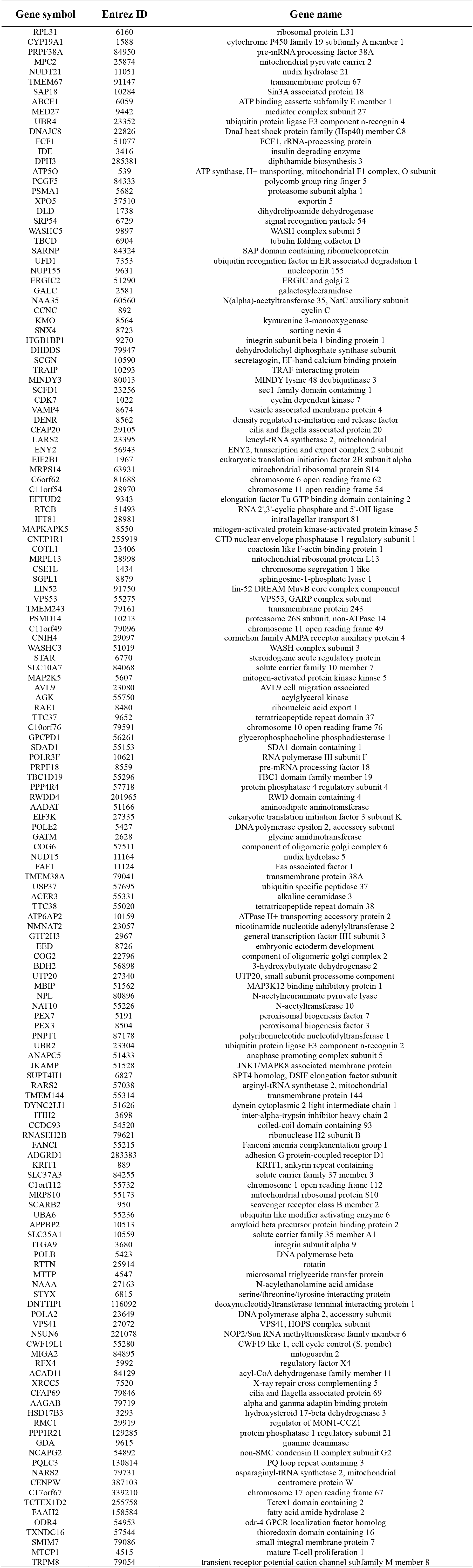
Representative human gene list in the single-copy orthologous gene families having correlated gene length with the maximum lifespan, the body weight, and the BMR simultaneously

**Table S24.**
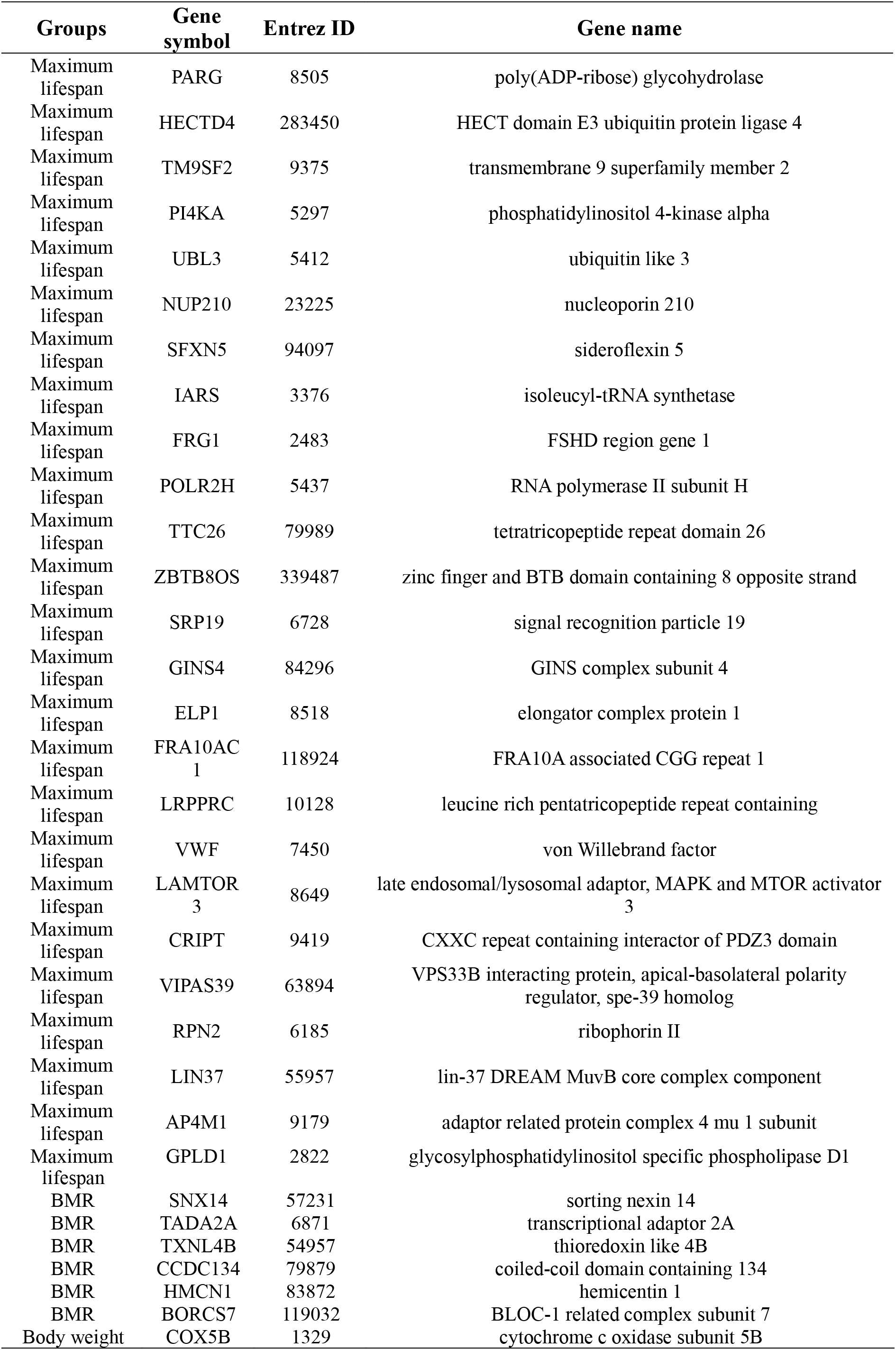
Representative human gene list of single-copy orthologous gene families with correlations between gene length and only maximum lifespan, only the body mass, or only the BMR, respectively

**Figure S24.**
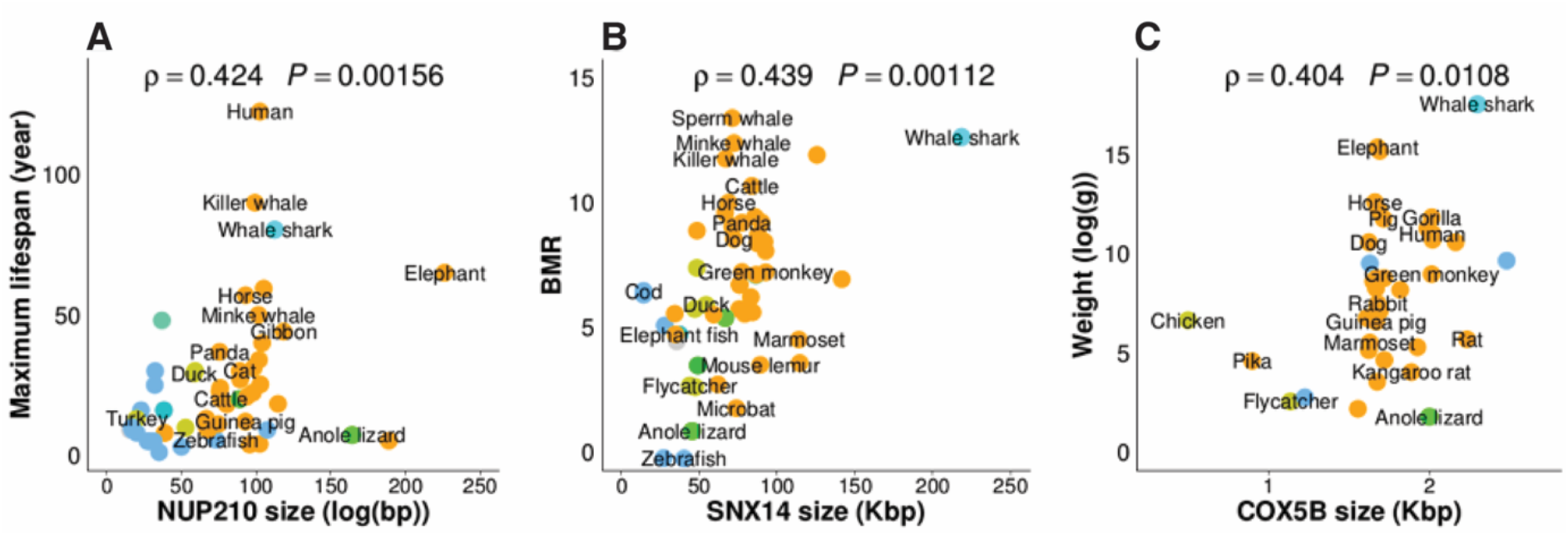
Single-copy orthologous gene families with correlations between gene length and maximum lifespan, weight, and BMR. Three instances of correlation between gene lengths of NUP210 (A), SNX14 (B), and COX5B (C) single-copy orthologous gene families and maximum lifespan, BMR, and body weight respectively. Dot colors represent the class as in Figure 1.

## 5. Scaling relationships

### 5.1 Scaling between genomic traits, physiological traits, and ecological parameters

We set out to determine whether the statistically significant correlations between genomic traits, physiological traits, and ecological parameters that we observed (Figure 2) in our array of species (centered on Chordates) reflect scaling relationships that may be formalized as power laws written as *Y = A*X^B^* ^40^. Consistent with previous work^40^, we found that the Basal Metabolic Rate (BMR) correlates with mass (B = 0.68, Figure S25). Furthermore, genome size, measured as golden path length, scales with gene size, measured as summed length of exons and intron per gene (B = 1.32, Figure S26), consistent with the observed lengthening of the whale shark genome by expanded CR1 repetitive elements (Figure 1A). Additionally, unlike in bacteria^58^ and crustaceans^59^, genome size in Chordates scales positively with temperature (B = 0.77, Figure S27).

**Figure S25.**
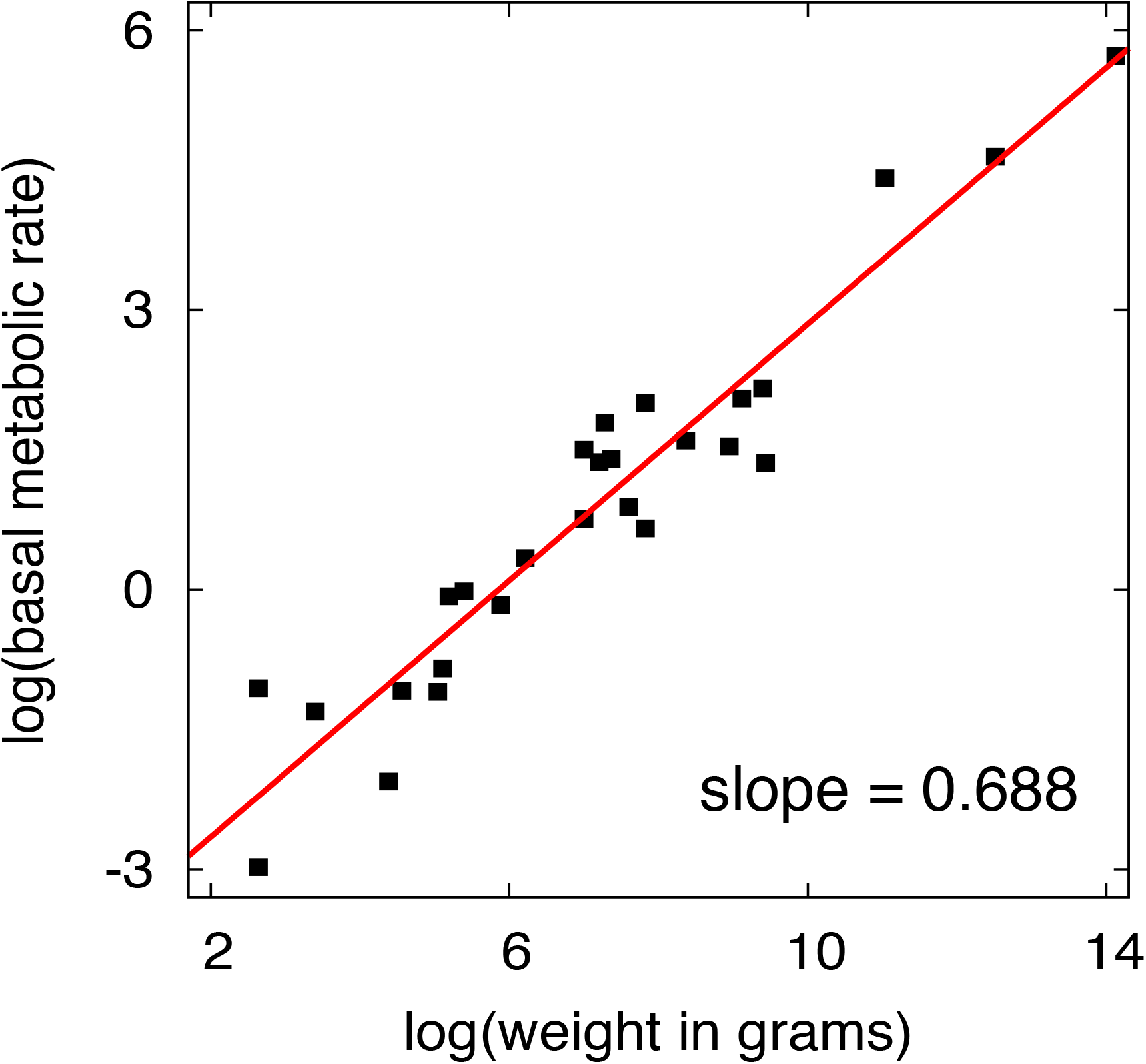
Scaling of basal metabolic rate to body size. Our regression analysis shows that, in 27 animal species, experimentally-determined BMR is positively scaled with body mass with an exponent B that is smaller than 1 (B = 0.688). Thus, BMRs are increased less than expected from body size.

**Figure S26.**
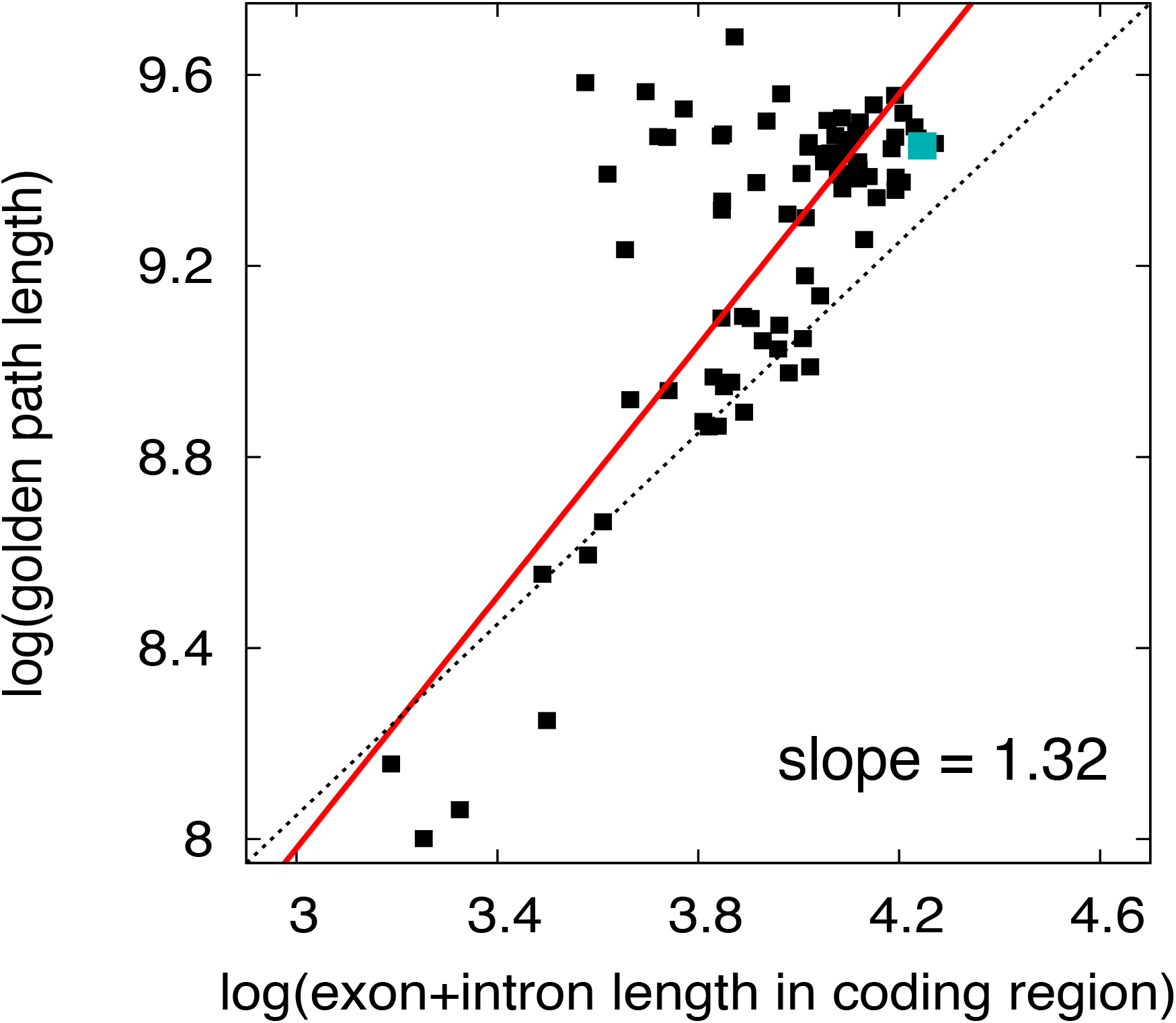
Scaling of genome size to gene size. Our regression analysis shows that, in 81 animal species, genome size (measured as golden path length) is positively scaled with gene size (measured for every gene as the sum of exons and introns) with an exponent B that is bigger than 1 (B = 1.32). Thus, genome size is significantly longer than expected from gene size alone.

**Figure S27.**
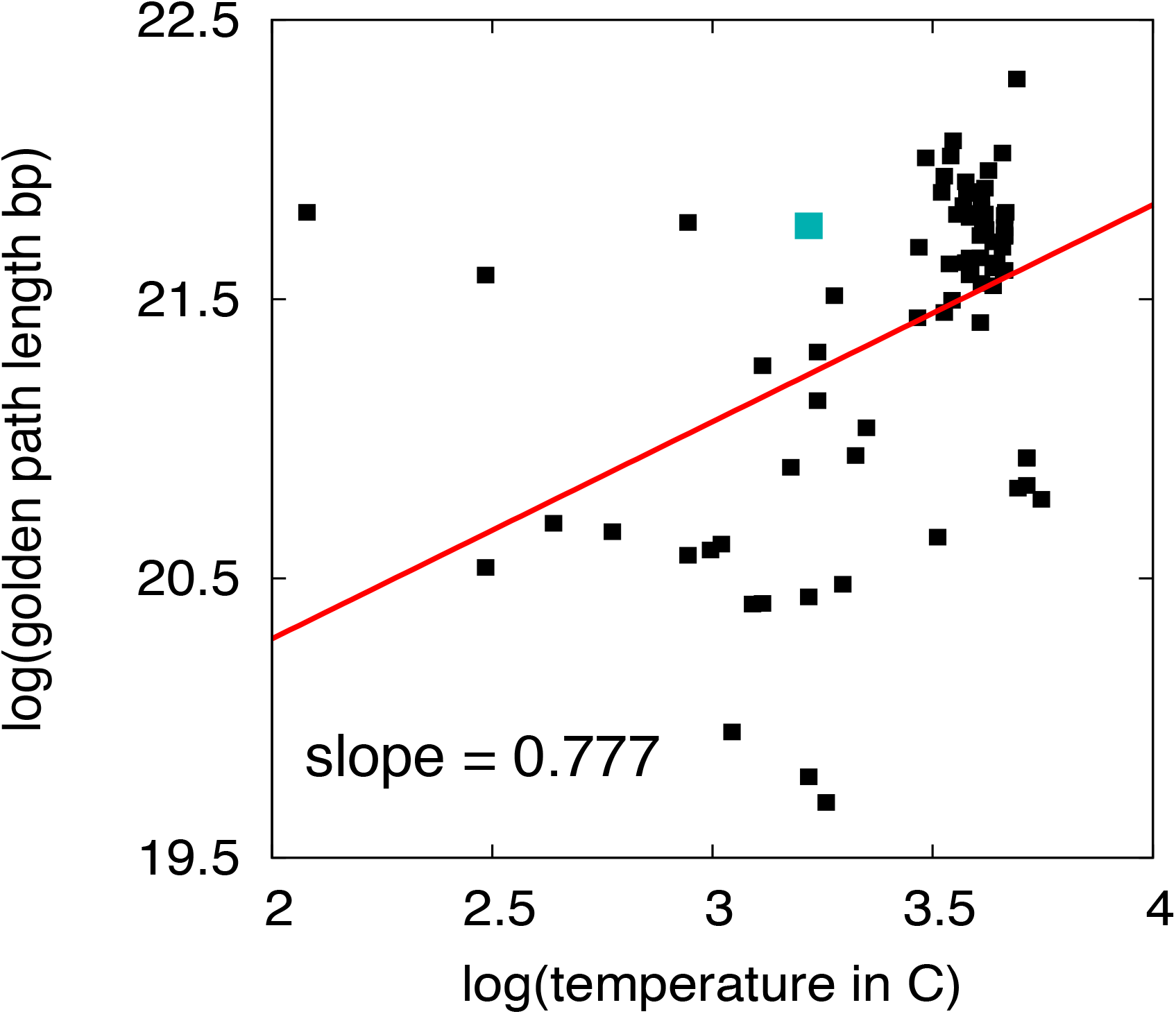
Scaling of genome size to temperature. Our regression analysis shows that, in 81 animal species, genome size (measured as golden path length) is positively scaled with temperature with an exponent B that is smaller than 1 (B = 0.777). Thus, genome size increases with temperature.

### 5.2 Scaling of neural genes to average gene lengths

Since several categories of neural genes are longer than average genes (Figure 4A, Figure S21), we examined whether their neural and average lengths obey a scaling relationship. Surprisingly, we found that neural genes are scaled to average genes with an exponent greater than 1 (B = 1.038, Figure S28), with the whale shark showing an extreme lengthening of neural genes.

**Figure S28.**
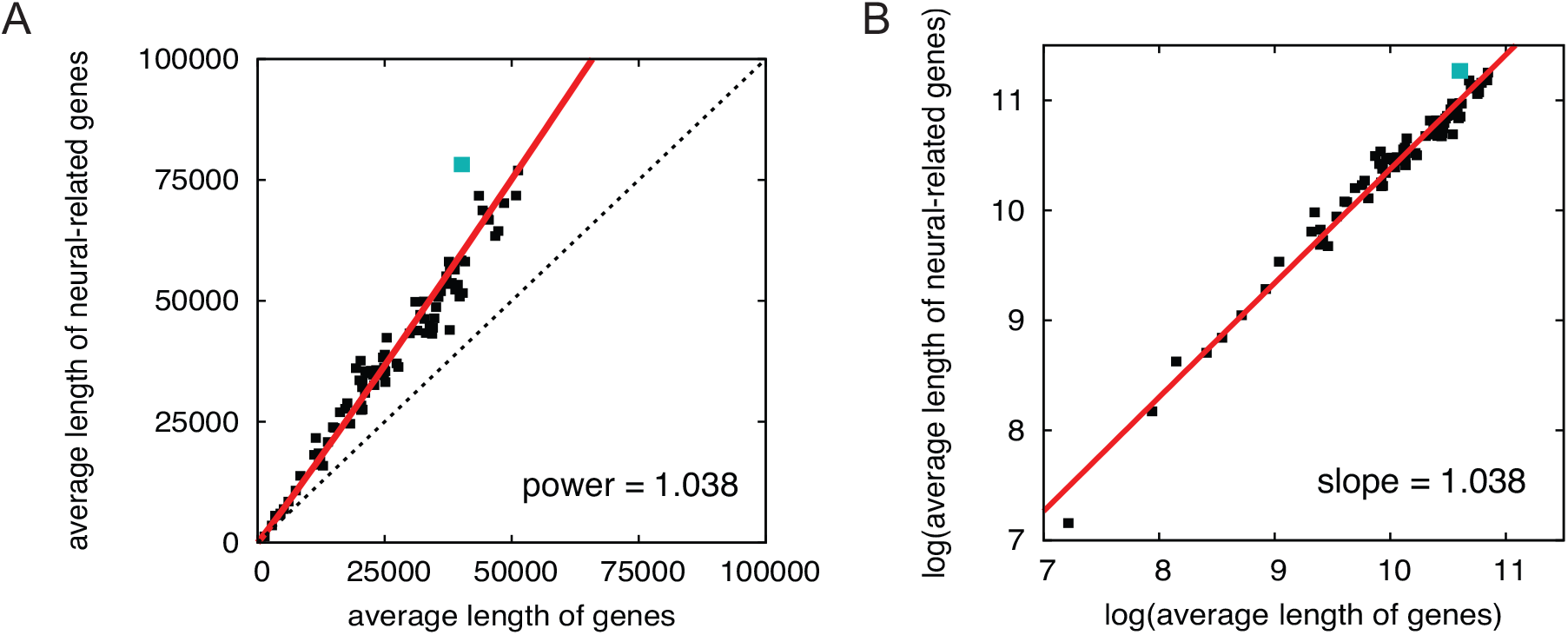
Scaling of neural genes to average gene size. Our regression analysis shows that, in 81 animal species, neural gene size is positively scaled with gene length (measured for every gene as the sum of exons and introns) with an exponent B that is bigger than 1 (B = 1.038). Thus, neural genes are longer than expected from gene size alone.

